# Structural mechanism of Gasdermin E-mediated mtDNA release from apoptotic mitochondria

**DOI:** 10.64898/2026.09.29.755391

**Authors:** Lisa Hohorst, Raed Shalaby, Nadine Gehle, Zakarya Benayad, Vincent Z Braun, Lars Richter, Sina Azami, KC Farrell, Asma Majoul, Shashank Dadsena, Eleonora G. Margheritis, Christina Kunz, Simone Prinz, Laurens Wachsmuth, Sabine Wilhelm, Shirin Kappelhoff, Alfredo Cabrera-Orefice, Ulrich Brandt, Manolis Pasparakis, Nieves Peltzer, Gerhard Hummer, Özkan Yildiz, Katia Cosentino, Ana J. Garcia-Saez

**Affiliations:** Institute of Genetics, CECAD, University of Cologne, Cologne, Germany; Department Membrane Dynamics, Max Planck Institute of Biophysics, Frankfurt am Main, Germany; Department of Biology/Chemistry and Center for Cellular Nanoanalytics (CellNanOs), University of Osnabrück, Germany; Department of Theoretical Biophysics, Max Planck Institute of Biophysics, Frankfurt am Main, Germany; IMPRS on Cellular Biophysics, Max-von-Laue-Straße 3, 60438 Frankfurt am Main, Germany; Centre for Molecular Medicine Cologne (CMMC) and Cologne Excellence Cluster on Cellular Stress Responses in Aging-Associated Diseases (CECAD), University of Cologne, Cologne, Cologne, Germany; Department of Structural Biology, Max Planck Institute of Biophysics, Frankfurt am Main, Germany; Electron Microscopy Facility, Max Planck Institute of Biophysics, Frankfurt am Main, Germany; Institute of Biochemistry, Faculty of Medicine, Justus Liebig University, Giessen, Germany; Research Institute for Medical Innovation, Radboud University Medical Center, Nijmegen, Netherlands; Department of Genome Editing, Institute of Biomedical Genetics (IBMG, University of Stuttgart, Stuttgart, Germany; Institute of Biophysics, Goethe University Frankfurt, 60438 Frankfurt am Main, Germany; Cluster of Excellence SubCellular Architecture of Life (SCALE), Goethe University Frankfurt, Frankfurt, Germany; Structural Biology Unit, Max Planck Institute of Biophysics, Frankfurt am Main, Germany; Department of Biomedical, Metabolic and Neural Sciences, University of Modena and Reggio Emilia, Italy

## Abstract

During apoptosis, the permeabilization of the mitochondrial inner membrane (MIM) through still-unclear mechanisms releases mtDNA into the cytosol, triggering the inflammatory cGAS/STING pathway under low caspase activity. Here, we report that, in apoptosis, the active form of the pore-forming protein Gasdermin E (GSDME-N) damages mitochondria before plasma membrane disruption. We visualize GSDME-N pore-like nano-assemblies in the MIM of apoptotic cells and of isolated mitochondria, which we bridge with the cryo-EM structure of the GSDME-N pore in mitochondria-like membranes. Deep membrane insertion of the anchor domain, which acts as a determinant of GSDME-N cardiolipin binding preferences, supports a role in pore formation. Notably, GSDME depletion results in a reduction in cristae swelling, MIM extrusion and mtDNA release during apoptosis. Subsequently, this decreases STING activation and inflammatory responses under caspase inhibition. Our findings reveal that GSDME mediates MIM permeabilization and mtDNA release during apoptosis and define the mechanism of GSDME-mediated membrane damage.

## Introduction

Apoptosis is a caspase-dependent form of regulated cell death with key roles in organ shaping, tissue homeostasis, and immune system function^1^. Dysregulation of apoptosis is associated with pathologies including cancer, neurodegeneration, or autoimmunity, which makes it an attractive therapeutic target^2^. The intrinsic pathway of apoptosis responds to intracellular stress and converges in the opening of pores at the mitochondrial outer membrane (MOM) by executioner members of the BCL-2 family, like BAX and BAK^3–5^. MOM permeabilization (MOMP) releases cytochrome c (cyt c) and SMAC into the cytosol, which activate apoptotic caspases, leading to the dismantling of cellular components and cell death acceleration^6,7^. BAX/BAK pores in the MOM also induce mitochondrial DNA (mtDNA) release into the cytosol^8,9^, which requires additional MIM permeabilization (MIMP) through a yet unknown mechanism.

The release of mitochondrial content downstream of mitochondrial BAX/BAK pores triggers competing signaling events. Cytosolic mtDNA initiates a type I interferon response via the pro-inflammatory cGAS/STING pathway. But activated apoptotic caspases following cyt c release in the cytosol inhibit this process ^6,7,10^. Furthermore, activated caspase-3 cleaves the pore-forming protein Gasdermin E and releases its active N-terminal fragment (GSDME-N), which can permeabilize the plasma membrane to induce a lytic form of apoptosis with features of pyroptosis^11,12^. GSDME-N has also been reported to target mitochondria to amplify cyt c release and caspase-3 activation through a positive feedback mechanism^13^, with a role in mitochondrial damage and neurodegeneration upon neurotoxin exposure^14^. GSDME is downregulated in many cancers and can activate anti-tumor immunity, likely associated with immunogenic cell death following its activation^15^.

GSDME belongs to the Gasdermin (GSDM) pore-forming protein family. Most of the structural and mechanistic understanding of GSDMs originates from studies of Gasdermin D (GSDMD) in the inflammasome pathways^16–20^. Active N-terminal GSDMD (GSDMD-N) assembles into ring-shaped complexes^21^ at the plasma membrane and reorganizes into a transmembrane β-barrel with a luminal pore, which permeabilizes the plasma membrane leading to cell death. This molecular mechanism is thought to be shared with other GSDM family members^22–24^, including GSDME^25^. Remarkably, GSDMD-N has been recently shown to induce mitochondrial damage and mtDNA release with consequences for the inflammatory outcome of pyroptosis^26^. Interestingly, GSDMD is inactivated by caspase-3 during apoptosis, indicating a specific role in mitochondrial damage during inflammasome-mediated cell death^27,28^.

However, little is known about GSDM structures in the membrane environment, the mechanism of action and function of active GSDME-N and its role in apoptosis. Here, we report the cryo-EM structure of the GSDME-N pore complex in mitochondria-like liposomes. Deep membrane insertion of the anchor domain is accompanied by membrane remodeling, explaining the molecular mechanism of how GSDME promotes pore formation. Molecular dynamics simulations, supported by experiments in model membranes and in cells, show preferential cardiolipin (CL) interactions with the anchor domain, which determines GSDME-N membrane targeting preferences. We also demonstrate that GSDME-N forms pore-like structures at the MIM, which we directly visualize. Importantly, we find that active GSDME can mediate MIMP and mtDNA release upon MOMP induction, which in presence of caspase inhibitors contributes to inflammatory signaling via the cGAS/STING pathway. Our findings reveal the structural mechanism for GSDME-N mitochondrial targeting and damage, and help consolidate the role of mitochondria in cell death-associated inflammation.

## Results

### GSDME-N can target and damage mitochondria independently of BAX and BAK

Like other GSDMs, GSDME-N shows higher affinity for the MIM lipid CL and localizes to mitochondria upon caspase-3 activation downstream of MOMP^13,26^. Furthermore, GSDME has been associated with mtDNA release from pyroptotic cells and with STING-driven inflammation in endothelial cells in atherosclerosis^29,30^. Based on this, we hypothesized that GSDME could be involved in MIMP and mtDNA release into the cytosol after MOMP in apoptosis. To test this, we first monitored GSDME cleavage and cellular distribution during apoptosis by immunoblotting of wild type (WT) mouse embryonic fibroblasts (MEFs) treated with 5 µM ABT737, 5 µM S63854 (AS) and/or 30 µM QVD (Q) for 2 h (Figure 1A). Subcellular fractionation indicated that cleaved GSDME-N accumulated in the mitochondria-enriched fractions isolated with anti-Tom22 magnetic beads (B), with cleavage only partially blocked under caspase inhibition. GSDME-N was also present in the 30,000 g pellet (crude lysosomal fraction, 30K), as well as in the microsomal 100,000 g pellet (100K) to a lesser extent. The cytosolic fraction (supernatant, SN) contained instead only full-length GSDME, suggesting that most cleaved GSDME is membrane-associated. We also detected similar GSDME cleavage and association with the crude mitochondrial fraction during apoptosis in human osteosarcoma (U2OS) cells, mouse small cell lung cancer (SCLC) cells and primary mouse lung fibroblasts (Suppl. Figure 1A-C). Experiments in MEFs deficient in caspases-9, -3 or -3/7 confirmed that GSDME-N generation during apoptosis was dependent on caspase-3 cleavage downstream of caspase-9 activation (Figure 1B and Suppl. Figure 1D). In line with Rogers et al. ^13^, recombinant GSDME-N activated *in vitro* by caspase-3 (Suppl. Figure 1E-G) bound to and permeabilized the MOM of isolated mitochondria, measured by cyt c release (Figure 1C). However, GSDME-N caused less cyt c release from mitochondria than cBID-activated BAK.

**Figure 1:**
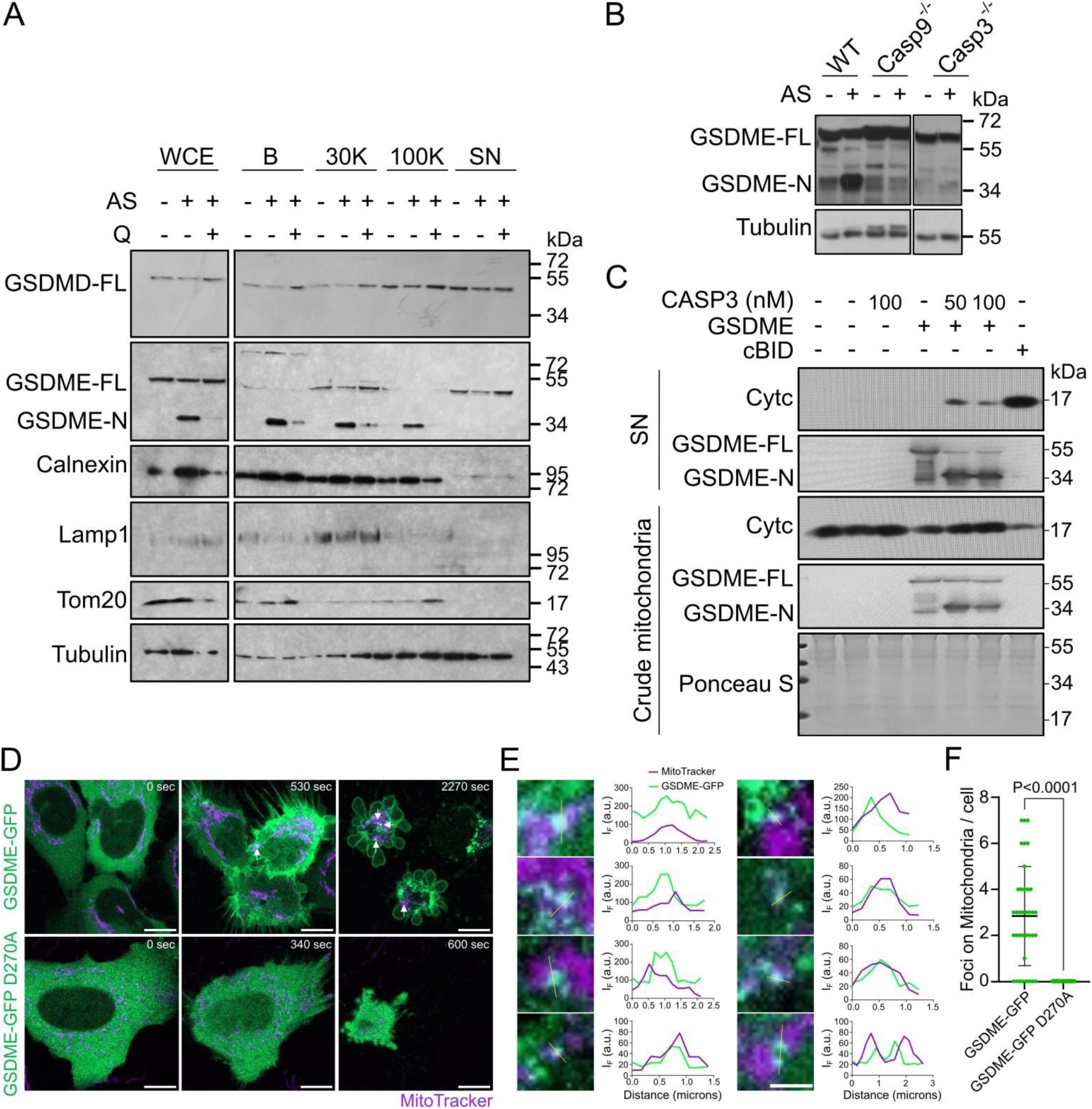
GSDME-N mitochondrial accumulation in apoptosis and damage. A) GSDMD and GSDME cleavage and subcellular localization during apoptosis were assessed by immunoblotting of whole cell extract (WCE), mitochondria isolated with Tom22 beads (B), 30,000 g pellet (30K), 100,000 g pellet (100K) and supernatant/cytosol (SN) fractions of MEF WT cells treated or not with 5 µM ABT737, 5 µM S63854 (AS) and 30 µM QVD (Q) as indicated, for 2h. B) Immunoblot of whole cell extracts showing caspase-3-dependent cleavage of GSDME downstream of caspase-9 activation. MEF cells of all genotypes were treated or not with 5 µM ABT737 and 5 µM S63854 (AS) for 2h. C) Immunoblot of cyt c release from crude isolated mitochondria from MEF WT cells treated or not with indicated proteins. Cyt c in the supernatant (SN) indicates release. D) Time-resolved imaging of hGSDME-GFP or hGSDME-GFP D270A co-transfected with Opto-hCaspase-9-mCherry in U2OS BAX/BAK DKO cells. Mitochondria were stained with MitoTracker Deep Red (magenta). hGSDME foci localized to mitochondria are indicated by white arrows. Scale bar: 10 µm. E) Representative magnified images from different cells showing hGSDME-GFP foci associated with mitochondria, together with the corresponding line profile analyses of the indicated regions. Line profiles display the fluorescence intensity of GFP and MitoTracker Deep Red along the indicated lines. Scale bar: 3 µm. F) Quantification of mitochondrial GSDME foci, as shown in D, throughout the entire acquisition period. Shown is the mean ± SD. n = 37 cells for hGSDME-GFP and n = 48 cells for hGSDME-GFP D270A. Statistical analysis by Welch’s t-test. All data were tested for normality.

To assess the intrinsic ability of GSDME-N to target mitochondria in cells independently of the apoptotic pore, we implemented an optogenetics system for light-controlled activation of GSDME in absence of apoptotic treatment, which we expressed in cells deficient in BAX and BAK. For this, Opto-hCaspase-9-mCherry was co-expressed in BAX/BAK-deficient cells (Suppl. Figure 1H) together with human GSDME-mEGFP, or with the Caspase-3 cleavage-site mutant hGSDME D270A-mEGFP (see Materials and Methods). The Opto-hCaspase-9-mCherry construct consists of hCaspase-9 fused to mCherry and Cry2 ^31^, a photo-inducible protein that oligomerizes upon blue light illumination, thereby triggering the oligomerization and activation of caspase-9, and thereby the apoptotic cascade downstream of MOMP (Suppl. Figure 1I). Before illumination, hGSDME-mEGFP, as well as the uncleavable mutant version, presented a diffuse cytosolic distribution. Upon light-induced activation of Opto-hCaspase-9-mCherry, hGSDME-mEGFP reorganized into a heterogeneous distribution, including discrete foci that co-localized with the mitochondrial signal, as well as with the plasma membrane, while hGSDME D270A-GFP remained cytosolic (Figure 1D,E). Quantification of the hGSDME-mEGFP foci co-localizing to mitochondria per cell upon light-inducing activation showed significant differences with the control sample, further supporting the evidence that cleaved GSDME can target mitochondria independently of MOMP (Figure 1F).

Together, these results indicate that, downstream of MOMP during apoptosis, GSDME-N accumulates at mitochondria and that it can damage the organelle also independently of BAX and BAK.

### Cryo-EM structure of GSDME in mitochondria-like liposomes

To understand the mitochondria-permeabilizing mechanism of GSDME, we solved the cryo-EM structure of the GSDME-N pore complex directly in CL-containing membranes. For this, we recorded cryo-EM movies of plunge-frozen proteoliposome samples and manually picked 285,000 particles of GSDME-N pores in vesicles (Figure 2A). Interestingly, we could detect GSDME-N pores occasionally formed in liposomes inside liposomes, which resemble the double-membrane topology of mitochondria. Analysis of 2D classes (Suppl. Figure 2) revealed a mixture of pore-complexes with 27-31 subunits, in line with the symmetry distributions observed for other pore structures of cleaved GSDMs^20,22,23^. Following iterative 2D averaging and 3D reconstruction, we achieved an optimized 3.6 Å map for the 30-subunit symmetry (Figure 2B). We fitted the model of soluble GSDME predicted by Alphafold^32,33^ to build an atomic model of the pore complex (Figure 2B, Suppl. Table 1). The resulting model shows the typical crown-like assembly with an inner diameter of 180 Å and an outer diameter of 300 Å (Figure 2B,D). Each GSDME-N subunit in the pore complex adopts a “hand-like” conformation similar to other GSDMs^20,22,23,34^, with the β-hairpins in the barrel forming the “fingers” and the globular domain the “palm”, in which the N-terminal α-helix and downstream loop would be the “thumb” (Figure 2C, Suppl. Figure 3).

**Figure 2:**
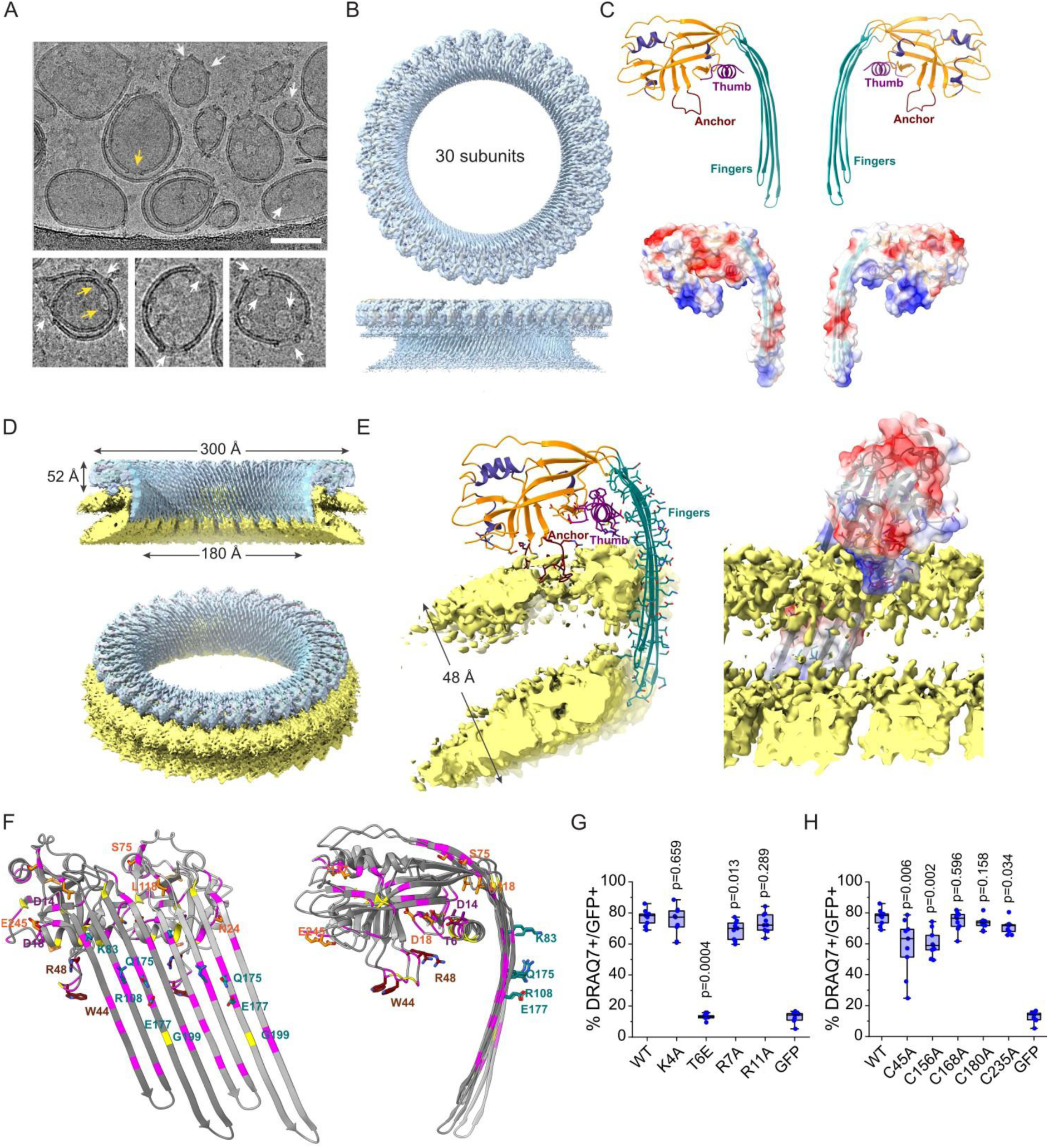
Cryo-EM structure of GSDME-N in CL-containing membranes. A) Cryo-EM micrograph of liposomes (PC:CL, 8:2) incubated with GSDME-N. Selected liposomes with GSDME-N pores (white arrows) on the outside are magnified in the lower row. GSDME-N can also form pores on the inner vesicles of multi-layered liposomes (yellow arrows). Scale bar: 100 nm. B) Overall structure of the GSDME-N pore complex. Single-particle cryo-EM map of GSDME-N with a resolution of 3.6 Å and a fitted model in ribbon representation. The ring-shaped pore complex with 30 subunits is shown from above (top) and parallel to the membrane (bottom). C) Membrane-inserted GSDME-N monomer seen from opposite sides. Top, the membrane-inserting β-strands (fingers) are colored in dark cyan, the N-terminal thumb helix in purple, and the anchor in brown. The other β-strands and helices in the subunit are shown in orange and slate blue, respectively. Bottom, the surface complementarity of neighboring subunits is shown in the surface potential presentation. Positively charged regions shown in blue and negatively charged areas in red. D) GSDME-N pore complex embedded in PC:CL membrane. Top, cross section through the ring parallel to the membrane with the overall dimensions indicated. The GSDME-N pores complex with 30 subunits has an inner diameter of 180 Å and an outer diameter of 300 Å. It protrudes 52 Å above the membrane. Bottom, full ring view. E) Interactions of GSDME-N with the lipid bilayer. A segment of the cryo-EM map corresponding to the headgroup region of the membrane (yellow) is shown with one GSDME-N subunit of the pore complex. The map for the lipid bilayer is shown in low contour level. On the left, GSDME-N is shown as a ribbon and stick model, while on the right it is rotated by 90° and presented as electrostatic surface. The hydrophobic tip of the GSDME-N anchor domain reaches the hydrophobic lipid core, while the positively charged residues located up- and downstream of the anchor domain and those of the thumb helix contact the lipid head groups of the membrane, contributing to the interactions of GSDME-N with the membrane. F) Location of GSDME-N mutations in cancer. Two subunits are shown in ribbon presentation from pore inside (left) and from the side (right). Residues mutated in cancer shown in magenta. Selected residues shown as stick model. Cysteine mutations tested in H highlighted in yellow. G,H) Functional analysis of selected GSDME-N-GFP mutants, overexpressed at similar levels. %DRAQ7 positive HeLa cells from cells transiently transfected with indicated GSDME-N-GFP mutants or with GFP at 24h. The center line of the box represents the median, while the box limits represent 25^th^ and 75^th^ percentiles. Data was analyzed with Mann-Whitney U test, with p-values compared to the WT sample.

The 30-subunit GSDME-N pore diameter is larger than that of GSDMA-N and GSDMB-N, with 24-27 subunits each ^20,22,23,24^. GSDMD-N with 33 subunits is slightly larger ^20,22,23^, while the 44-subunit GSDM from *Trichoplax adhaerens* (*Tricho*GSDM) is the largest (Suppl. Figure 4A-C)^34^. Comparison of the individual subunits shows structural similarity in the palm and slight differences in the anchor (Suppl. Figure 4D). The fingers of GSDME in the pore are longer than those of GSDMA, GSDMB, and GSDMD but comparable to those of *Tricho*GSDM, although they differ in the curvature, with the angle between palm and fingers being larger in *Tricho*GSDM (Suppl. Figure 4D). GSDMEN fingers are also straighter and less curved. The surface analysis of the pore complex shows predominantly negatively charged clusters on the solvent-exposed side of the globular domain and in the fingers facing the inside of the pore (Figure 2C,E and Suppl. Figure 5). In contrast, the membrane-facing side of the thumb helix and the tips of the fingers are mainly positively charged. The highly positive anchor region stands out in particular, although its tip is hydrophobic.

In contrast to previous detergent-solubilized Gasdermin pore complex structures^20,22,23,34^, our GSDME-N structure was solved in the intact lipid bilayer of liposomes, which allowed us to gain insight into the interactions with the membrane (Figure 2D,E). In the transmembrane β-barrel, the finger residues 88-96, 99-107, 176-186, 191-201 facing the membrane form a hydrophobic patch that directly contacts the lipid core of the bilayer. Further, the β-hairpins at the tip of the fingers are in a more extended conformation compared to other GSDMs (Suppl. Figure 4). Several, individually-unresolved lipid molecules are present between the hydrophobic patch of the fingers, the hydrophobic tip and basic residues of the anchor, and the basic residues of the thumb helix (Figure 2E), suggesting a direct role of these regions in membrane binding. Remarkably, the lipid bilayer is thinner in the region between anchor and the finger tips, indicating membrane thinning in the pore proximity.

The structure also revealed that the hydrophobic anchor tip (^43^FWCW^46^) inserts deep into the hydrophobic region of the bilayer and is stabilized there by direct van-der-Waals interactions with the lipids. These residues are flanked by mainly basic residues (^40^KKR^42^ and ^47^QRK^50^), which interact with the negatively charged lipid headgroups. It is striking that the lipid density below these residues is noticeably weaker, which could be explained by the fact that the anchor protrudes deep into the bilayer and displaces lipids laterally. The insertion of the anchor domain into the bilayer is asymmetric, adding material only to the outer leaflet, which could underlie the thinning observed in the pore proximity and thereby contribute to reducing the energy for pore formation. Accordingly, mutations in the anchor domain of *Tricho*GSDM reduce its pore-forming activity^34^.

The mapping of GSDME mutations identified in cancer patients^15^ to our GSDME-N structure reveals several clusters (Figure 2F). Mutations in the fingers appear preferentially in the luminal side of the pore, with most of them facing the pore lumen and others the membrane. Changing the hydrophilic residues in the pore lumen would impact the electrostatic processes during pore formation, while mutating the residues on the other side would impact the membrane binding. Mutations in the anchor would affect the interaction of GSDME with the membrane (see above). Furthermore, another cluster concentrates at the residues of the thumb α-helix, the downstream loop and its immediate surroundings.

To evaluate the role of the thumb, we performed functional studies with GSDME-N mutants overexpressed in cells (Figure 2G). Mutations R7A or R11A had a mild or no effect, suggesting that their positive charge is compensated by neighbouring residues and that they are individually dispensable. In contrast, phosphomimetic mutation T6E completely abolished GSDME-killing activity, indicating that a negative charge at this position is not tolerated. Together, these results indicate that the thumb contributes to GSDME-N pore activity, likely through electrostatic interactions with the membrane that might cooperate with the anchor domain to promote membrane thinning.

Unlike GSDMD, where Cys191/Cys192 (human/mouse) is palmitoylated for protein activation ^35–40^, GSDME-N does not contain an equivalent cysteine residue. Besides C45A in the anchor domain, substitution of each one of the cysteine residues in GSDME-N only had minor effects on its cell-killing activity. This suggests that individual cysteines are not essential for pore activity (Figure 2H), although GSDME palmitoylation was detected in previous work^36,41^.

### GSDME-N targets mitochondria prior to plasma membrane disruption

The results obtained in Figure 1 suggested that GSDME-N pores could potentially mediate MIMP during apoptosis, which we aimed to test by direct visualization. Considering the inner diameter of about 18 nm of the GSDME-N pore complex (Figure 2D), we reasoned that DNA-PAINT would provide the required spatial resolution^42^. Since apoptosis is a dynamic, irreversible process, we selected an appropriate time point for visual inspection of the fixed cell samples shortly after MOMP, but prior to robust plasma membrane permeabilization. For this, we visualized and quantified the kinetics of mitochondria fragmentation (indicated by a reduction in the number and geometry of branches) and of SMAC-mCherry release into the cytosol, as proxies for MOMP, in parallel to SYTOX uptake, as a marker for plasma membrane permeabilization, in U2OS cells transiently expressing full-length human GSDME tagged with GFP (hGSDME-mEGFP) upon apoptosis induction (Suppl. Figure 6A-H). We identified t=120 fulfilling these conditions. Of note, the absence of detectable SYTOX uptake does not exclude limited plasma membrane permeabilization at this stage. For these experiments, we analyzed cells with comparable GSDME expression levels, selecting cells with moderate GSDME expression and avoiding those with very high levels.

Using multi-color Airyscan live cell microscopy, we visualized the dynamics of the subcellular distribution of GSDME using a hGSDME-cpHalo construct in cells undergoing apoptosis at the single cell level. We stained mitochondria and the plasma membrane with AKAP1-mEGFP-Hpep3 (AKAP1-Hpep) and puro-leader-aALFAnb-TMD-mEGFP-Hpep3 (TMD-Hpep), respectively. As shown in Figure 3A-C (and Suppl. Figure 6I, J), we found that overexpressed hGSDME-cpHalo was initially cytosolic, and translocated first to mitochondria and later to the plasma membrane within the same individual cells during apoptosis progression. Because of technical limitations, we cannot distinguish whether the GSDME molecules localized to mitochondria and later to the plasma membrane correspond to the same population. Yet, this striking observation indicates that GSDME localizes to mitochondria early during apoptosis and later accumulates to the plasma membrane. Quantification of hGSDME-Halo localization in U2OS cells fixed at different time points after apoptosis induction showed that GSDME consistently localized to mitochondria at 120 minutes, and only at later stages appeared at the plasma membrane (Figure 3D,E, Suppl. Figure 6K,L). Importantly, by staining the MOM and MIM simultaneously, we could clearly detect individual GSDME puncta at both mitochondrial membranes, and especially co-localizing with MIM regions exposed to the cytosol through MOM openings (Figure 3F,G). Similar results were obtained with WT U2OS cells expressing hGSDME-mEGFP under the weak PGK promoter, which resulted in expression levels comparable to endogenous GSDME, and in WT HeLa cells transiently expressing hGSDME-GFP, with small differences in the timing of the events (Suppl. Figure 7 and 8A-G).

**Figure 3:**
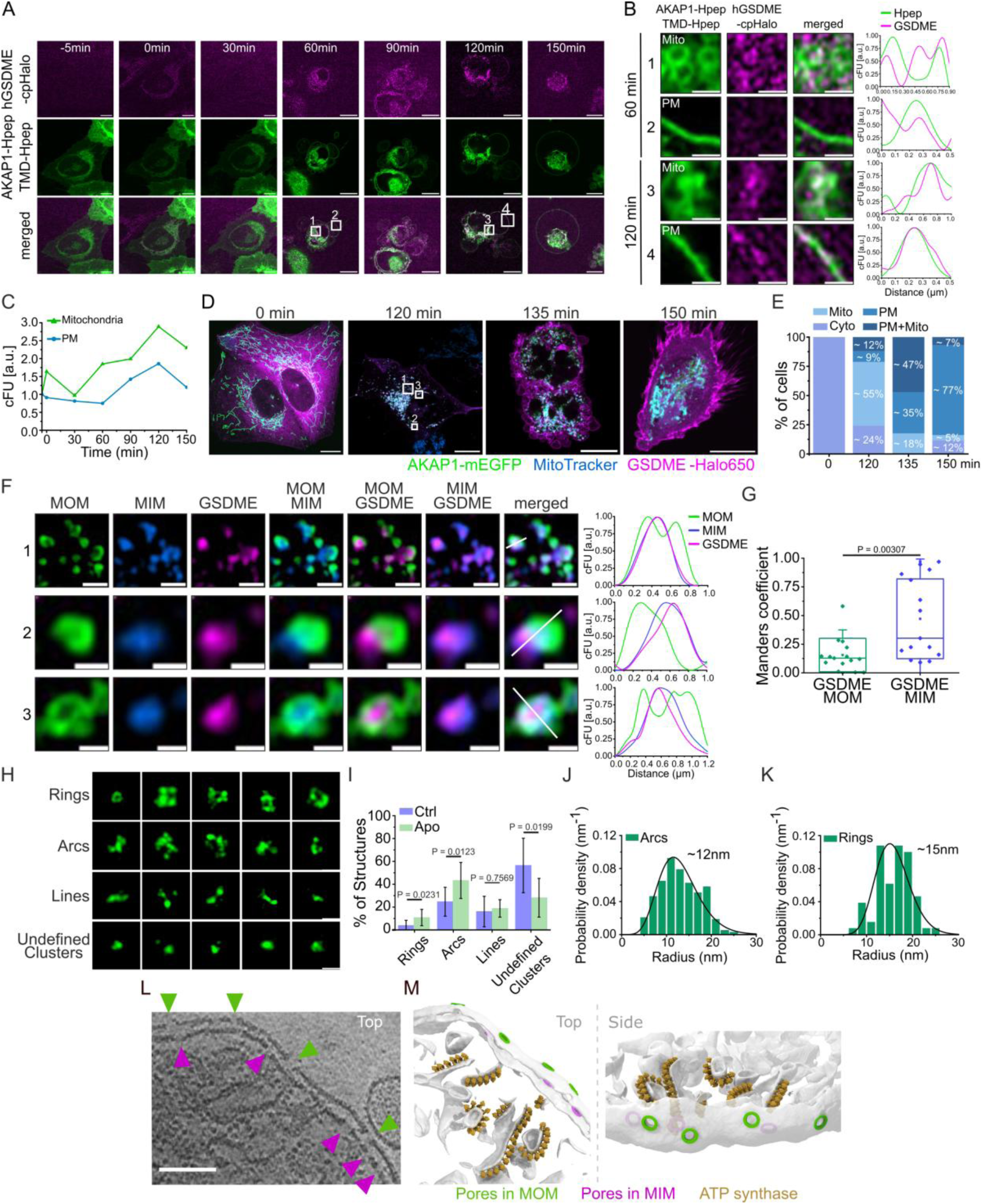
Visualization of GSDME-N pore-like assemblies at the mitochondrial inner membrane. A) Representative Airyscan time-lapse images of apoptotic U2OS WT cells expressing hGSDME-cpHalo (magenta), AKAP1-mEGFP-Hpep3 and puro-leader-aALFAnb-TMD-mEGFP-Hpep3 (green). The -5 min images are before HTLJFx650 addition and before apoptosis induction. Cells were then labeled with 50 nM HTLJFx650, and apoptosis was induced with 1 µM ABT-737 and 1 µM S63845. Scale bar: 10 µm. B) Airyscan microscopy images of zoomed regions in A at 60 min (Images 1 and 2) and 120 min (Images 3 and 4) after apoptosis induction. The line profiles of cross-sections show colocalization of hGSDME with mitochondria (Images 1 and 3) and PM (Images 2 and 4) in corrected fluorescence intensity (cFU). Scale bar: 1 µm. C) Quantitative analysis of hGSDME fluorescence intensity (corrected fluorescence intensity units, cFU) at mitochondria vs PM over time in apoptotic U2OS WT cells shown in (A). Data normalized to -5 min time point. Data are representative of two independent experiments (n = 2 with 8 cells). D) Representative Airyscan microscopy images of U2OS WT cells expressing AKAP1-mEGFP (green) and hGSDME-HaloTag (magenta) at 0 min, 120 min, 135 min and 150 min after apoptosis induction with 1 µM ABT-737 and 1µM S63854. Matrix/MIM stained with MitoTracker Red (blue). White boxes in the 120 min timepoint image correspond to zoomed regions in F. Scale bar: 10 µm. E) Quantification of hGSDME cellular distribution at different time points after apoptosis induction in experiments performed as in D. Cells were classified according to hGSDME localization as cytosolic (Cyto), mitochondrial (Mito), at the plasma membrane (PM) or both (PM + Mito) (n = 3 with >15 cells for each timepoint). F) Airyscan microscopy images of zoomed regions in D at 120 min after apoptosis induction. The line profiles of cross-sections show colocalization of GSDME with MIM in corrected fluorescence intensity (cFU). Scale bar: 1 µm for image 1 and 0.5 µm for images 2 and 3. G) Co-localization analysis of GSDME with MOM and MIM at 120 min after apoptosis induction. Data are presented as box plots with the center lines at the median, lower bound at 25th percentile, upper bound at 75th percentile, and individual data points represent cells (n = 3 with 16 cells). H) Gallery of hGSDME-mEGFP structures localized at the MIM of apoptotic U2OS WT cells resolved with DNA-PAINT microscopy. Scale bar: 50 nm. I) Distribution of mitochondrial GSDME structure types in apoptotic (n = 9 cells, 5 experiments, 498 structures) and untreated (n = 10 cells, 4 experiments, 361 structures) U2OS WT cells expressing hGSDME-mEGFP. J-K) Quantification of radius distribution of hGSDME-mEGFP ring- and arc-like structures at mitochondria of apoptotic U2OS WT cells. Ring average radius is 15 nm and arc average radius is 12 nm (Rings n = 56, Arcs n = 207; n = 9 cells; 5 experiments). L) Representative image of the top view of a central slice through a tomographic volume of mitochondria isolated from mouse liver and incubated with recombinant hGSDME-N. The MIM can be distinguished from the MOM by the presence of the ATP synthase. Scale bar: 100 nm. M) Top (left) and side (right) view of the segmentation of the region of interest in I) showing the inner and outer mitochondrial membrane (white), GSDME-N pores in the MOM (green), GSDME-N pores in the MIM (magenta), and the ATP synthase (gold). All statistics were measured by a two-tailed unpaired t-test with p>0.05 = non-significant. All data were tested for normality.

### GSDME-N forms pore-like structures at the mitochondrial inner membrane

We then resolved the nanoscale organization of the GSDME puncta localized at the MIM of healthy and apoptotic cells using dual-color DNA-PAINT (Suppl. Figure 8H-K). In agreement with the enrichment in crude mitochondria fractions (Suppl. Figure 1A-C), we also detected GSDME puncta at mitochondria of untreated cells (Suppl. Figure 8H,J). We could identify a mixture of distinct structures, shaped as rings, arcs and lines, and undefined clusters, with rings and arcs significantly enriched in apoptotic compared to healthy cells (Figure 3H,I). The size analysis revealed a radius of ∼15 nm for rings and ∼12 nm for arcs located to mitochondria (Figure 3J,K), which refer to the outer radius based on the label positioning on GSDME, in line with the ring outer radius of the cryo-EM pore structure (15 nm, Figure 2D).

To gain more insight into the topology of GSDME-N pore-like structures relative to the inner and outer mitochondrial membranes, we visualized GSDME-N pore structures in isolated mitochondria by cryo-electron tomography (cryoET). We incubated recombinant GSDME cleaved in vitro by caspase-3 with mitochondria isolated from mouse liver. The reconstructed tomograms clearly showed the presence of mitochondria with a double membrane system in which the ATP synthase complexes could be detected in only one of them, allowing an unambiguous distinction of the MOM and MIM (Figure 3L,M and Suppl. Figure 8L). Importantly, we could clearly identify GSDME-N pore complexes in the MIM, as well as in the MOM (Figure 3L,M, Suppl. Figure 8L). Similar results were obtained with mitochondria isolated from *Polytomella*, a unicellular organism with very characteristic mitochondria containing a high number of ATP synthase complexes in their MIM (Suppl. Figure 8M-N)^43^. This, together with the super-resolution imaging data, demonstrates that GSDME-N can form pores in the MIM, for which it has to first permeabilize the MOM and then pass through its own pores or, in apoptotic cells, reach the MIM once the MOM is permeabilized by BAX and BAK.

### The anchor domain determines GSDME-N mitochondrial targeting

To understand which molecular features in GSDME-N mediate CL and mitochondria membrane binding, we performed large-scale atomistic molecular dynamics (MD) simulations of 30-mer GSDME-N rings embedded in a lipid bilayer composed of 20% CL and 80% di-oleoyl phosphatidylcholine (DOPC) with explicit solvent (with systems of >4 million atoms). We also performed coarse-grained MD simulations at the Martini level of resolution. We consistently found that CL molecules bind preferentially to basic patches on the GSDME-N pore at the interface of the globular domain with the membrane and between monomers (Figure 4A-F, Suppl. Figure 9A-C and Suppl. Movie 1). Specifically, CL interacts with the basic regions (residues 39-42, 48, 49) and the hydrophobic tip (residues 43-46, including W44 and W46) in the anchor domain of one monomer, as well as with the inter-monomer region (residues 238–245) of the neighbouring monomer. These strong and preferential interactions emerge as likely drivers for mitochondrial targeting, with CL being a highly abundant lipid (∼20%) in the MIM^44^, accessible in apoptotic cells. In comparison, the concentration of phosphatidylinositol 4,5-bisphosphate (PIP_2_) at the plasma membrane, previously involved in interactions of membrane targeting of GSDMD-N, has been estimated at <1%^45^.

**Figure 4:**
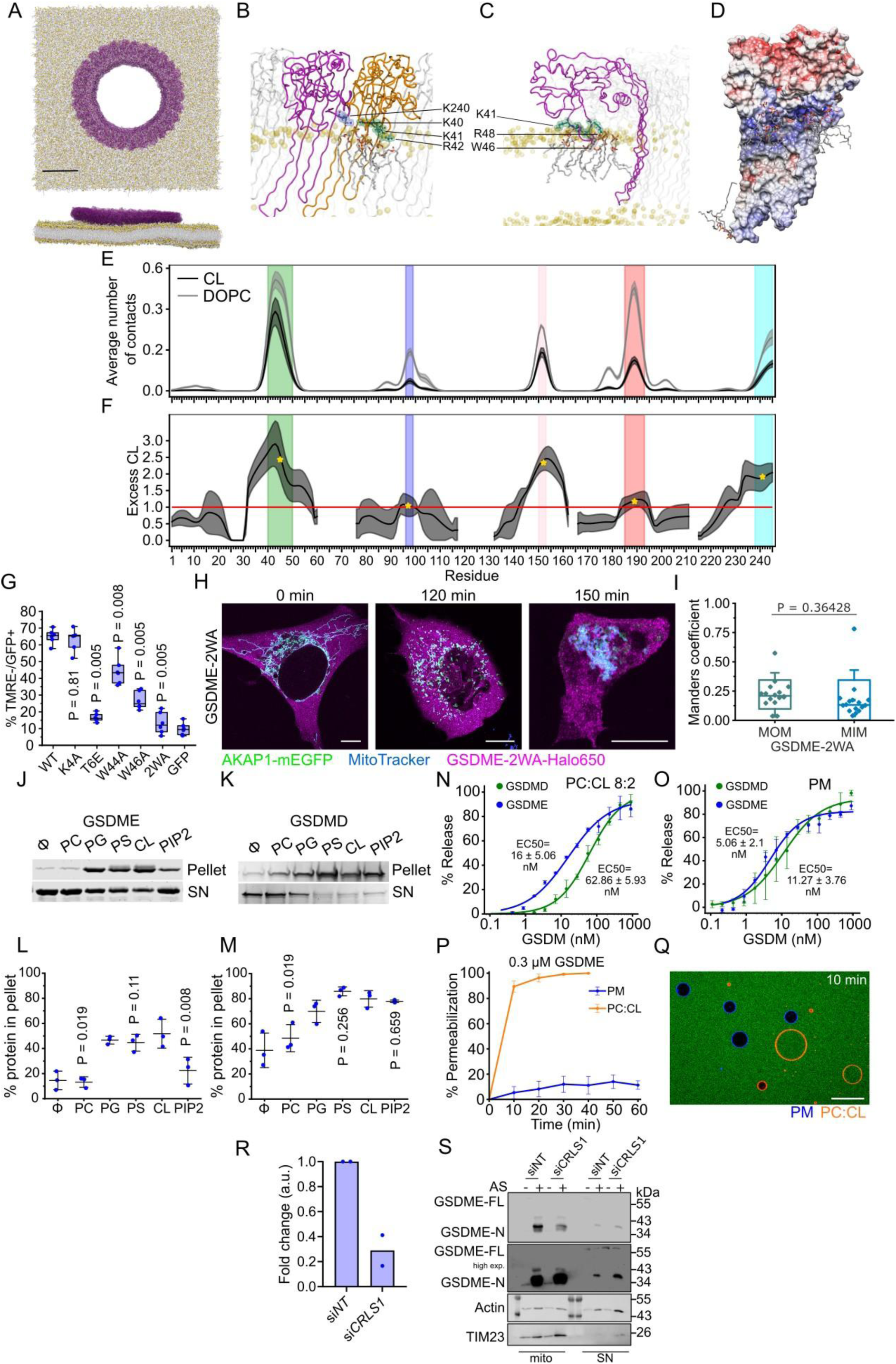
The anchor domain determines the lipid binding preferences of GSDME-N. A) Top and side views of the simulated system. GSDME-N pore: purple; lipid phosphate headgroups: yellow spheres; lipid tails: gray sticks; water and ions not shown for clarity; scale bar: 10 nm. B-C) Detailed interactions of cardiolipins with residues belonging to the basic regions (K40, K41, R42 and R48, highlighted in green), the hydrophobic tip (W46, highlighted in pink), and the inter-monomer region (K240, highlighted in blue). (B) Snapshot zooming in on the interface between two monomers, colored purple and orange, with cardiolipin lipids bridging them. Two cardiolipins are shown: one interacts with K240 of the purple monomer and K40 of the orange monomer, while the other interacts with K40 and R42 of the orange monomer. (C) Side view of a monomer, shown in purple, with two cardiolipins interacting with the basic residues K41 and R48. Residue W46, belonging to the hydrophobic tip, is embedded in the membrane and interacts with the cardiolipin lipids via their lipid tails. In both snapshots, the rest of the GSDME pore is shown in transparent white, membrane phosphorus atoms are shown as transparent yellow spheres, and water molecules, ions, and hydrogen atoms of the cardiolipins are not represented for clarity. The data shown are derived from the 3rd all-atom replica, at time point 180 ns. D) Electrostatic surface map of two neighboring GSDME-N monomers. The atomic coordinates were extracted from an all-atom molecular dynamics simulation of the GSDME-N pore. The electrostatic potential is shown on the protein surface, with a color scale ranging from red (negatively charged) to blue (positively charged) regions. Five interacting CL molecules are displayed, illustrating preferential binding of the phosphate headgroups to positively charged (blue) regions on the protein surface. E-F) Residue-resolved number of CL molecules (E) and local CL enrichment (F) averaged across 30 monomers and three simulation replicas (dark gray; only lipid headgroup contacts are considered). For comparison, DOPC contacts are shown in light gray in E. The CL enrichment was computed as the CL:DOPC contact ratio normalized by the overall lipid composition (20% CL, 80% DOPC). Curves were smoothed using a Gaussian filter (width=2 residues). Shaded areas around the curves represent the standard deviation across replicas. CL binding regions are indicated by colored shading as in B. Golden stars in F indicate average enrichment within each region. CL molecules bind preferentially to residues 40–50 (green) and, 150–153 (pink) from one GSDME-N monomer, and 238–245 (light blue) from the neighboring monomer, with significantly higher CL contact ratios in this bridging position than expected from lipid membrane composition (dashed red line). G) Functional analysis of selected GSDME-N-GFP mutants, overexpressed at similar levels. %TMRE-negative HeLa cells (mitochondrial depolarization) from cells transiently transfected with indicated GSDME-N-GFP mutants or with GFP at 24 h. The center line of the box represents the median, while the box limits represent 25^th^ and 75^th^ percentiles. Data was analyzed with Mann-Whitney U test, with p-values compared to the WT sample (n=3 independent experiments with 2 replicates each). H) Representative Airyscan microscopy images of U2OS WT cells expressing AKAP1-mEGFP (green) and hGSDME-W44A-W46A-HaloTag (magenta) after apoptosis induction with 1 µM ABT-737 and 1µM S63854 at 0 min, 120 min and 150 min. Matrix/MIM stained with MitoTracker Red (blue). Scale bar: 10 µm. I) Co-localization analysis of hGSDME-2WA with MOM and MIM at 120 min after apoptosis induction. Data are presented as box plots with the center lines at the median, lower bound at 25th percentile, upper bound at 75th percentile. n = 3 with 18 cells. J-K) Liposome sedimentation assay. The presence of GSDME-N (J) and GSDMD-N (K) in the pellet indicates binding to LUVs with indicated compositions (PC, 100%PC; PG, PC:PG (6:4); PS, PC:PS (6:4); CL, PC:CL (8:2); PIP,2 PC:PIP2 (9:1)). Φ, control without LUVs. L-M) Quantification of GSDME-N (L) and GSDMD-N (M) bound to LUVs in J-K as % protein in the pellet. Shown is mean ± SD. n=3. Statistical analysis was performed by two-tailed unpaired t-test, with p-values reported in comparison to the CL sample. N-O) GSDME-N and GSDMD-N concentration-dependent carboxyfluorescein release from LUVs made of PC:CL (8:2) (N) or PM (PC:PE:SM:Chol:PS:PI:PIP2, 18:6:10:29:10:5:2) (O), after 1 hour of incubation. Shown is the mean ± SD. n=3. P) Competitive assay of GSDME-mediated permeabilization of liposomes mimicking the mitochondrial (orange, 80% PC, 20% CL, 0.1% DiI) and the plasma membrane (blue, 50% PC, 30% PE, 10% Chol, 10% PS, 0.1% DiO) compositions. Normalized permeabilization of GUVs (indicated by uptake of Alexa488) over time upon addition of 0.3 µM of GSDME-N to a mixture containing both mitochondria- and plasma membrane-like GUVs. Shown is the mean ± SD. n=3 independent experiments. Q) Representative section of the tile set showing GUVs with mitochondrial membrane-mimicking lipid composition (PC:CL (8:2), orange) and plasma membrane-mimicking lipid composition (PM=PC:PE:Chol:PS (5:3:1:1), blue) 10 min after GSDME-N addition. Scale bar: 30 µm. R) Levels of RNA of CRLS1 after 72h of siRNA knockdown normalized to the non-targeting (si*NT*) control and to h*HPRT* (n = 2 independent experiments). S) Representative immunoblot of mitochondria-enriched (mito) and cytosolic (SN) fractions showing the reduction of GSDME-N in the mito fraction upon CRLS1 knockdown in apoptotic conditions (1 µM ABT-737+ 1 µM S63845; AS) (n = 2 independent experiments).

The anchor domain indeed presents several key features of CL binding domains^46^, in particular short stretches of aromatic residues flanked by positively charged amino acids. By systematic mutation of these residues in the anchor domain, we observed the most dramatic reduction in mitochondrial damage and loss of cell-killing activity with the “2WA“ double mutant W44A/W46A (Figure 4G and Suppl. Figure 9D. Accordingly, full-length GSDME-2WA activated during apoptosis remained mostly cytosolic, in contrast to the WT protein (Figure 4H,I and Suppl. Figure 9E) compared with Figure3D,E). Mutations in the lysine residues of the anchor domain to alanine also reduced the killing activity, though to a lesser extent (Suppl. Figure 9D). The least effect on the activity was observed when the arginines in the anchor domain were mutated to alanine (Suppl. Figure 9D). The CL-interacting residues identified in the MD simulations thus have a key role in membrane targeting.

We then compared the sequence and structure of the anchor domains of different GSDMs (Suppl. Figure 9F-K). Those of GSDME and GSDMD are similarly arranged with a hydrophobic tip flanked by basic residues, suggesting a common basis for their lipid binding preferences. However, the anchor domains of GSDMA, GSDMB and specially of GSDMC, are more divergent. They do not present the two W amino acids at the hydrophobic tip, and in the case of GSDMC, the tip is flanked by both negative and positively charged residues. In line with this, overexpression of a GFP-tagged construct corresponding to the fragment of GSDMC-NT equivalent to that of GSDME-NT and GSDMD-NT presented cytosolic localization and reduced cell-killing activity (Suppl. Figure 9L).

We next compared the relative ability of recombinant human GSDME-N and mouse GSDMD-N cleaved *in vitro* by caspases-3 and -4, respectively, to bind and permeabilize liposomes of different lipid compositions (Figure 4J-Q). We found that GSDME-N exhibited a slightly higher preference for CL-containing liposomes over those containing phosphatidyl serine (PS) and phosphatidyl glycerol (PG), and a lower binding affinity to PIP_2_ (Figure 4J,L). Instead, GSDMD-N seemed to bind with highest affinity to PS, closely followed by CL and PIP2, and slightly less to PG (Figure 4K,M). Of note, these results indicate lipid binding preferences, not selectivity, since both GSDMs could bind to negatively charged lipids, but not to neutral phosphatidyl choline (PC). With respect to membrane permeabilizing activity, GSDME-N presented higher activity in mitochondria-like membranes compared to GSDMD-N, and a similar activity in liposomes with a lipid composition mimicking the plasma membrane (Figure 4N,O and Suppl. Figure 9M-P). In addition, an important difference was in their permeabilization rate, with GSDME-N permeabilizing membranes, specially plasma membrane-like, slower than GSDMD-N (Suppl. Figure 9Q,R).

To further strengthen the evidence about the preference of GSDME-N for CL, we performed experiments of competitive liposome permeabilization by mixing giant unilamellar vesicles (GUVs) mimicking the mitochondrial and the plasma membrane compositions together with an external fluorescent probe in the same reaction sample. As shown in Figure 4P,Q (and Suppl. Figure 9S), when simultaneously exposed to membranes containing CL (mitochondria-like composition) and to a composition similar to that of the inner leaflet of the plasma membrane (without CL), GSDME-N permeabilized mitochondria-like liposomes faster and more efficiently, as indicated by the entry of the external probe into the GUVs, in line with the results obtained in Figure 3.

Finally, to explore the effect of cardiolipin on the ability of GSDME to target mitochondria, we knocked down CRLS1, the enzyme responsible for cardiolipin synthesis, in U2OS cells ^26,47^. Remarkably, we found that the accumulation of GSDME-N in mitochondria-enriched fractions upon apoptosis induction was reduced in these cells compared to control cells (Figure 4R,S). These results are consistent with a reduction of cardiolipin hindering the recruitment of GSDME to mitochondria. Importantly, they also strengthen the evidence that GSDME indeed targets mitochondria during apoptosis under endogenous conditions.

In summary, our findings provide evidence that different GSDMs present distinct membrane binding preferences and identify the anchor domain of GSDME as a key region for CL and mitochondrial targeting.

### GSDME mediates MIMP and mtDNA release during apoptosis

To evaluate the role of GSDME on MIMP during apoptosis, we compared the mitochondrial alterations in healthy and apoptotic wild-type (WT) versus *Gsdme*^-/-^ primary mouse lung fibroblasts. Treatment with BH3-mimetics induced a slightly reduced amount of cell death in the knockout cells compared to the WT cells, but mitochondrial potential loss, ROS production, and caspase activation, showed comparable kinetics (Figure 5A-D). We reasoned that GSDME-N pores at the MIM would cause matrix swelling as a result of water influx due to osmotic imbalance, enhancing MIM/matrix extrusion through the apoptotic pores. EM analysis of mitochondrial ultrastructure showed that apoptosis induction in WT cells caused cristae remodeling and swelling, as expected^47^. Remarkably, the mitochondria of apoptotic *Gsdme*^-/-^ cells presented lower circularity and remodeled cristae with reduced width compared to WT cells (Figure 5E-G and Suppl. Figure 10A). In addition, apoptosis induction in WT cells led to MIM extrusion from the MOM, measured by counting single extruded MIM events per cell as well as by a decrease in the overlap between MOM and MIM, quantified by co-localization analysis with Manders coefficient, in agreement with previous studies^7,8^. In contrast, in apoptotic *Gsdme*^-/-^ cells the extent of MIM extrusion was significantly reduced, although the MOM seemed permeabilized and the mitochondria fragmented, indicating that MIM extrusion but not MOMP was affected (Figure 5H-J and Suppl. Figure 10B).

**Figure 5:**
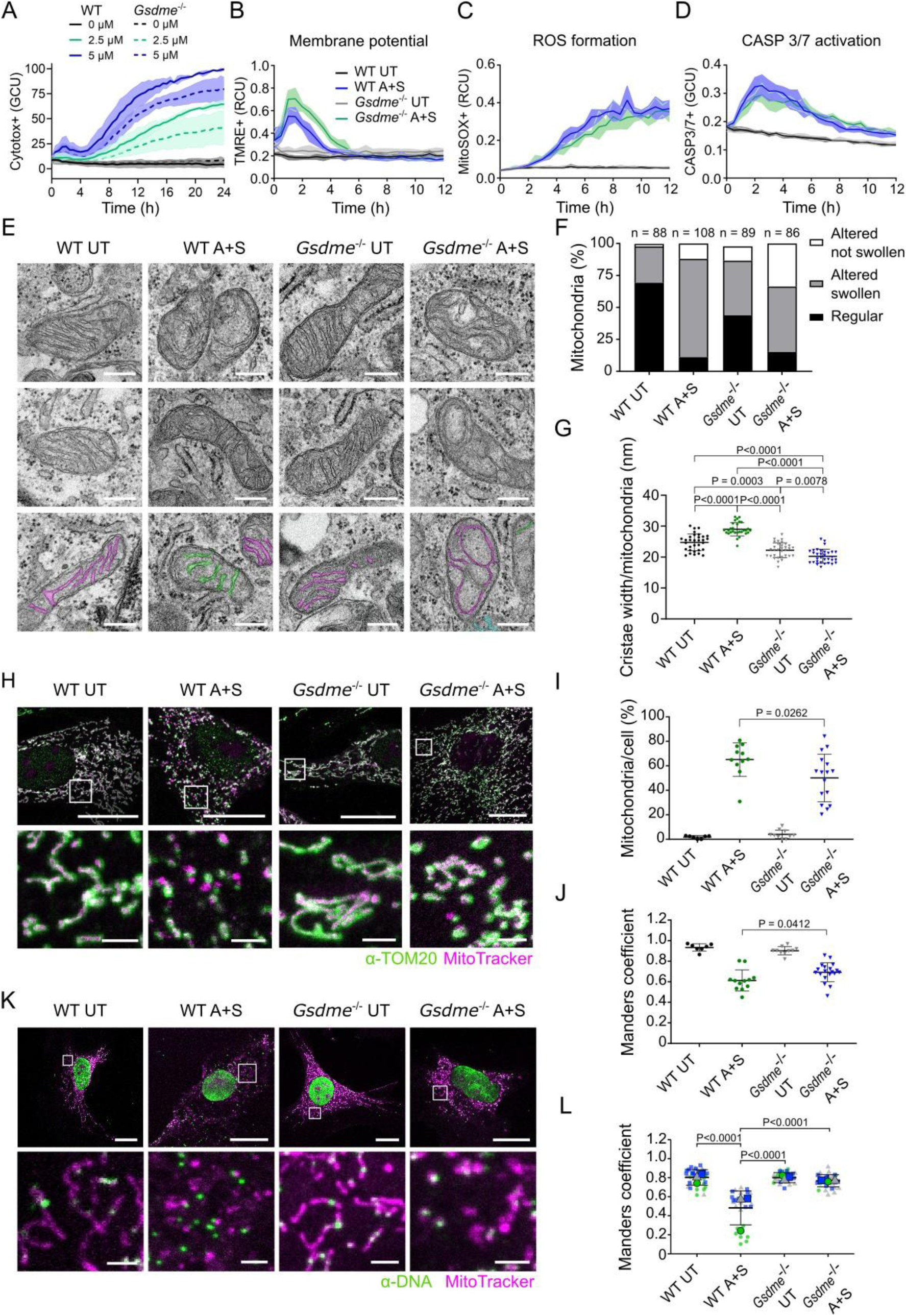
GSDME promotes MIM permeabilization and mtDNA release in apoptosis. A) Effect of GSDME depletion on cell death measured by Cytotox green uptake in UT cells with 2.5 µM and 5 µM A+S. Mean ± SD is shown. Graphs represent n = 3 independent experiments. B-D) Effect of GSDME depletion on mitochondrial membrane potential loss measured by TMRE (B), ROS formation measured by MitoSOX red (C), and Caspase 3/7 activation (D) in UT and A+S treated cells. Mean ± SD is shown. Graphs represent n = 3 independent experiments. E) Gallery of transmission electron microscopy images of mitochondria under indicated conditions. Scale bar: 250 nm. Bottom row, colored cristae indicate the masks created for width quantification. F) Quantification of cristae morphology per mitochondria, classified as altered not swollen (white), altered swollen (grey) and regular(black) cristae. Data represent n > 80 mitochondria from 4–5 individual cells per condition. G) Quantification of mean cristae width per mitochondria. Mean ± SD is shown, dots represent single mitochondria in individual experiments. Data represent n = 29-32 mitochondria from 4–5 individual cells per condition. Statistical analysis by Ordinary one-way ANOVA corrected for multiple comparisons with Tukey test with p>0.05 = non-significant. H) Visualization of MIM extrusion. Confocal images of MIM/matrix (labeled with MitoTracker orange, magenta) and MOM (labeled with anti-TOM20 antibody, green). Top, cell overview. Bottom, zoomed images of the areas in white boxes in the top. Scale bar: 20 µm, and 2 µm for zoomed areas. I) Quantification of MIM extrusion events per mitochondria per cell. Mean ± SD is shown, dots represent single cells in individual experiments. Data represent the status of the MIM counted for n=27-150 mitochondria from 6-16 independent cells per condition. Statistical analysis by two-tailed Welch’s *t*-test with p>0.05 = non-significant. J) Quantification of MIM extrusion by co-localization analysis of MIM and MOM. MitoTracker labels the MIM/matrix and TOM20 labels the MOM. Mean ± SD is shown, dots represent mitochondria of individual cells. n = 7-19 individual cells per condition. Statistical analysis by two-tailed Welch’s *t*-test with p>0.05 = non-significant. K) Visualization of mtDNA release by confocal microscopy. Mitochondria stained with MitoTracker orange (magenta), DNA labeled with anti-DNA antibody (green). Top, cell overview. Bottom, zoomed images of the areas in white boxes in the top. Scale bar: 20 µm, and 2 µm for zoomed areas. L) Quantification of mtDNA release. Co-localization analysis of mtDNA and MitoTracker signals. Mean ± SD is shown. Big symbols represent the mean of three independent experiments (n = 3) and small symbols represent single cells of one individual experiment. Statistical analysis by two-tailed Welch’s *t*-test with p>0.05 = non-significant. Experiments were performed in WT or *Gsdme*^-/-^ primary mouse lung fibroblasts untreated (UT) or treated with 5 µM A+S for 2h unless otherwise indicated. A, ABT-737; S, S63845 WT. All data were tested for normality.

To evaluate whether the reduced MIM alterations in apoptotic *Gsdme*^-/-^ cells were accompanied by altered permeability, we quantified mtDNA release. Apoptosis induction caused cytosolic relocalization of mtDNA in WT cells, quantified by a decrease in co-localization with mitochondria (Figure 5K,L and Suppl. Figure 10C). Importantly, this was largely inhibited in *Gsdme*^-/-^ cells, demonstrating GSDME’s role on mtDNA release into the cytosol during apoptosis.

As additional evidence for the functional role of GSDME on MIM permeabilization during apoptosis, we also quantified the activation of the STING pathway upon apoptosis induction under caspase inhibition using HeLa cells stably expressing a recently developed STING biosensor^48^ (Figure 6A,B). As expected, STING activation under these conditions was dependent on mtDNA, since it was blocked in Rho0 cells lacking mtDNA (Figure 6C and Suppl. Figure 10D). In line with a reduction in mtDNA release, GSDME depletion significantly reduced STING activation in this system, but not upon direct STING activation with the agonist diABZi (Figure 6D and Suppl. Figure 10E,F). Furthermore, the downstream induction of interferon β was significantly decreased in *Gsdme*^-/-^ primary mouse lung fibroblasts treated with AS+Q, compared to WT cells (Figure 6E-G). Interferon α and CXCL10 expression also showed a decreasing trend, although the reductions were less pronounced. Similar results were obtained in a different model of SCLC cells lacking GSDME (Suppl. Figure 10G-L).

**Figure 6:**
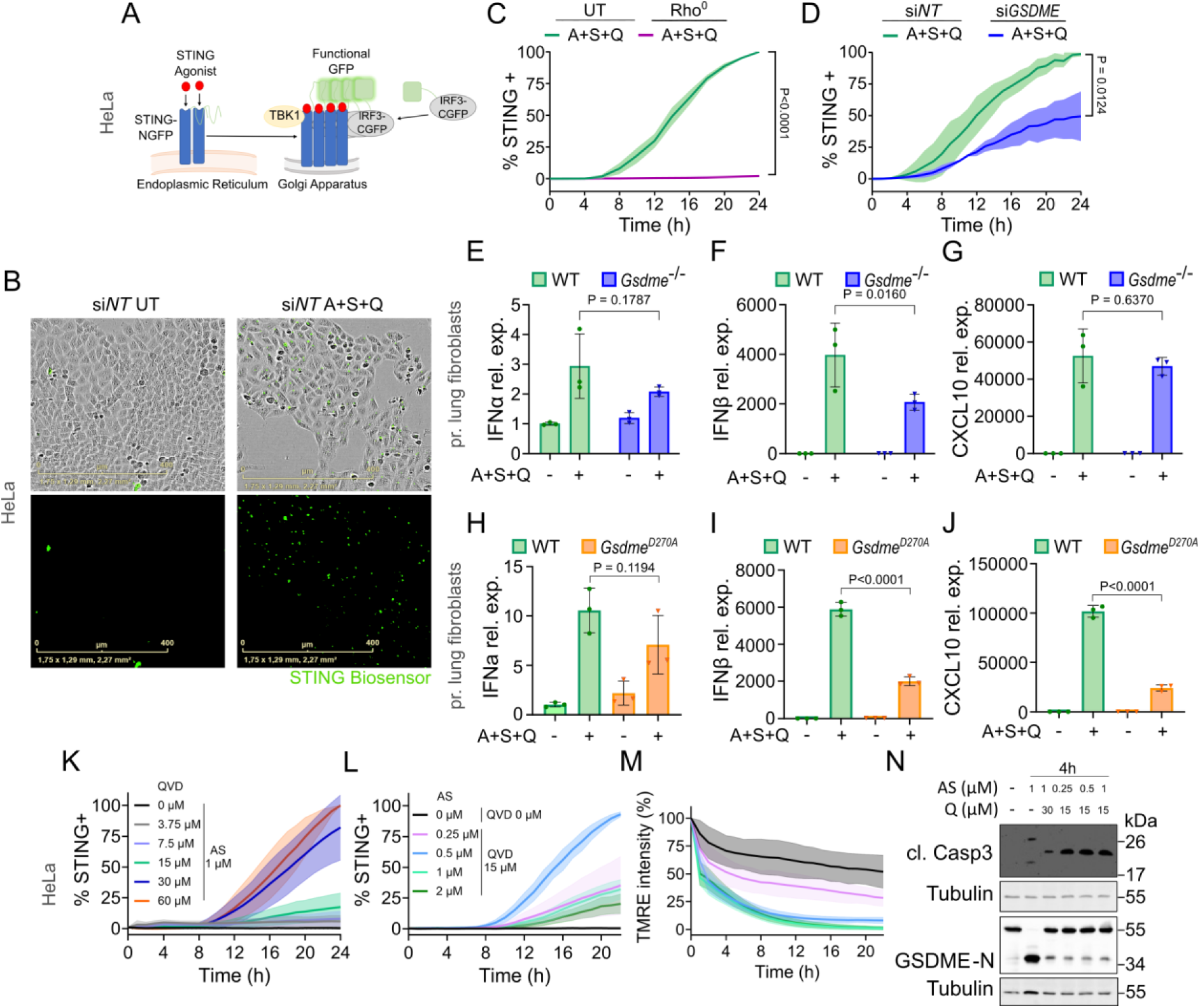
Partial caspase inhibition permits GSDME cleavage while failing to suppress STING signaling. A) Scheme of the STING biosensor. HeLa cells stably expressing STING fused to the N-terminal fragment of GFP and IRF3 fused to the C-terminal fragment of GFP, which are not fluorescent on their own. Binding of cGAMP or a STING agonist to STING induces its activation and translocation from the ER to the Golgi apparatus, where it interacts with IRF3, leading to complementation of the N- and C-terminal fragments of GFP, which becomes fluorescent. B) Representative IncuCyte images of HeLa cells expressing the STING biosensor transfected with non-targeting control siRNA either untreated (UT) or treated with 1 µM A+S and 10µM Q. The activation of STING can be observed by green spots (corresponding with Golgi) forming in the cell. Scale bar: 400 µm. C) Effect of mtDNA depletion (Rho^0^) in HeLa STING biosensor cells on STING activation after A+S+Q treatment normalized to time point 0 min. Mean ± SD is shown. n = 3 independent experiments. Statistical analysis by Wilcoxon test with p>0.05 = non-significant. D) Effect of GSDME knockdown in HeLa STING biosensor cells on STING activation after A+S+Q treatment normalized to time point 0 min. Mean ± SD is shown. n = 3 independent experiments. Statistical analysis by Wilcoxon test with p>0.05 = non-significant. E-G) qPCR analysis of *Ifnα* (E), *Ifnβ* (F), and *Cxcl10* (G) relative expression (rel. expr.) after 6 h treatment with A+S+Q in WT and *Gsdme^-/-^* mouse primary lung fibroblasts. ΔΔCt method. The data were normalized to untreated WT. n = 3 independent experiments. One-way ANOVA followed by Tukey’s post-hoc test with p>0.05 = non-significant. H-J) qPCR analysis of *Ifnα* (E), *Ifnβ* (F), and *Cxcl10* (G) relative expression (rel. expr.) after 6 h treatment with A+S+Q in WT and *Gsdme D270A* mouse primary lung fibroblasts. ΔΔCt method. The data were normalized to untreated WT. n = 3 independent experiments. One-way ANOVA followed by Tukey’s post-hoc test with p>0.05 = non-significant. K) Titration of the caspase inhibitor QVD at the indicated concentrations in combination with constant BH3-mimetics (AS, 1 µM) in HeLa STING Biosensor cells to measure STING activation. n = 3 independent experiments. L-M) Titration of BH3-mimetics (AS) at the indicated concentrations in combination with constant caspase inhibition (QVD, 15 µM) in HeLa STING Biosensor cells to measure STING activation (L) and mitochondrial membrane potential loss via TMRE (M). n = 3 independent experiments. N) Representative immunoblot of HeLa STING Biosensor cells showing caspase-3 and GSDME cleavage after 4 h of treatment at indicated BH3-mimetics (A+S) and caspase inhibitor (Q) concentrations. n = 3 independent experiments. A = ABT-737 and S = S63854 (A+S) and Q = QVD.

These results were intriguing because they suggest that caspase-3 activation downstream of MOMP would play a dual role. On the one hand, it would promote inflammatory signaling by cleaving GSDME: this would lead to mtDNA release and cGAS/STING-mediated interferon responses, in addition to triggering the known switch from apoptosis to a pyroptotic-like form of cell death. On the other hand, caspase-3 has been reported to dampen inflammatory signaling in apoptosis, among other by cleaving cGAS and inhibiting the STING pathway^10^. To explore the role of caspase activity on these effects, we used cells expressing a non-cleavable form of GSDME (GSDME D270A) to avoid the confounding effects of caspase activity on cGAS/STING (Suppl. Figure 10M,N). Importantly, the expression of interferon α, β, and CXCL10 was again reduced in *Gsdme^D270A^* primary lung fibroblasts to a similar extent as the *Gsdme*^-/-^ cells treated with AS+Q, compared to WT cells, demonstrating the contribution of GSDME cleavage to mtDNA-mediated interferon responses (Figure 6H-J).

To account for the effects of active GSDME on STING activation in presence of caspase inhibitors, we hypothesized that the activation level of caspase-3 downstream of MOMP could lead to scenarios with partial cleavage of cGAS and GSDME, sufficient to cause MIM permeabilization and mtDNA release, but insufficient to block STING activation. Consistent with this, we detected a weak band of cleaved GSDME under our apoptotic conditions with caspase inhibition, suggestive of residual caspase-3 activity (Figure 1A). To further explore this possibility, we searched for a level of caspase activity at which both events could take place (Figure 6K-N and Suppl. Figure 10O,P). We first titrated the concentration of the caspase inhibitor to determine the lowest concentration at which STING activation could be detected during apoptosis, identifying 15 µM of QVD as a threshold in our system (Figure 6K). Then, we titrated the concentration of BH3 mimetics to identify a concentration at which STING activation and MOMP were simultaneously induced in the presence of 15 µM QVD (Figure 6L,M, Suppl. Figure 10O,P). We also checked by western blot whether caspase-3 and GSDME were cleaved under these conditions. Remarkably, upon treatment of the HeLa STING Biosensor cells with low concentrations of BH3 mimetics and caspase inhibitor (15μM), both caspase-3 and GSDME appeared cleaved at 4 h of treatment conditions, under which we also detected strong STING activation (Figure 6N). Importantly, these results reveal that cells can undergo apoptosis under conditions in which caspase activity is sufficient to cleave and activate GSDME but insufficient to block the STING pathway.

Together, these results indicate that, under our experimental conditions, GSDME knockout hinders MIM permeabilization in apoptosis, reducing cristae swelling, MIM/matrix extrusion through the MOM and mtDNA release during apoptosis. This is accompanied by a reduction in STING activation and downstream transcriptional responses, under caspase inhibition. We identify experimental conditions of caspase activity levels compatible both with STING activation and with GSDME cleavage, which is necessary for cytokine induction. These findings uncover a role for GSDME in the inflammatory signaling associated with content release from the mitochondrial matrix downstream of MOMP in apoptosis within a window of caspase activity.

## Discussion

We report the cryo-EM structure of GSDME-N pore complex in liposomes, which provides mechanistic understanding of how GSDME (and likely other GSDMs) interacts with membranes and induce pore formation. We find that the aromatic residues in the anchor domain insert deeply into the hydrophobic region of the lipid bilayer and displace lipids only in one leaflet, which is associated with membrane thinning in the pore surroundings and indicates a contribution to the reduction of the energy cost for pore opening^49^. This reveals a mechanism how asymmetric anchor domain insertion into the membrane, likely in coordination with electrostatic interactions of the thumb and finger tips with the lipid headgroups, facilitates membrane remodeling and opening of pores by GSDME-N.

We also uncover the role of the anchor domain as a structural feature mediating the preferential binding of GSDME to CL and required for mitochondria targeting using a combination of MD simulations, reconstituted systems and cellular models. Although the anchor domain of GSDME shares structural similarities with that of GSDMD, the proteins present different lipid binding preferences. Both proteins bind to a range of negatively charged lipids. Yet, GSDMD-N binds equally well to plasma membrane and mitochondrial lipids, while GSDME-N binds mitochondrial CL preferentially. Consistent with the role of CL in the mitochondrial targeting of GSDME, we find that endogenous GSDME-N recruitment to mitochondria in apoptosis is reduced when the mitochondrial enzyme responsible for CL synthesis, CLS1, is knocked down.

Strikingly, we discover dynamic changes in the subcellular distribution of GSDME during apoptosis progression. Mitochondrial accumulation takes place at earlier stages and is followed by plasma membrane accumulation and permeabilization. This is in line with our results in giant vesicles showing that GSDME-N preferentially permeabilizes CL-containing membranes over membranes mimicking the plasma membrane, when simultaneously exposed to them. We hypothesize that GSDME lipid binding preferences could underlie their distinct rate of membrane targeting and pore formation, ultimately leading to distinct functional consequences in the membrane-damaging effects found in cells. The structural and mechanistic basis of this temporal shift in GSDME localization remains an open question that will deserve further investigation.

We also directly visualized GSDME-N pore-like nano-assemblies at the MIM of apoptotic cells and of isolated mitochondria with comparable sizes to the cryo-EM GSDME-N pore structure reported here. In addition to rings, we identified GSDME-N assemblies shaped as arcs and lines. Although these structures may partially result from the imaging of the assemblies at different tilt angles, we have previously observed them also in the context of a flat plasma membrane^21^, suggesting their existence in cells. Arc- and line-shaped assemblies of GSDMs and other pore forming proteins have also been found *in vitro*, in atomistic molecular simulations and in cells^3,4,50,51^. The inability to detect line- and arc-shaped GSDME-N structures in liposomes by cryo-EM may result from differences in experimental conditions or regulatory mechanisms of pore formation in mitochondria.

Importantly, GSDME depletion altered MIM remodeling and reduced MIM/matrix extrusion through the MOM and mtDNA release in apoptotic cells. This was accompanied by a decrease in STING activation and the induction of inflammatory transcriptional responses under caspase inhibition. These findings indicate that GSDME can mediate MIMP and mtDNA release, contributing to the activation of the cGAS/STING pathway and inflammatory signaling downstream of MOMP, in line with^6,7^.

We show that GSDME cleavage downstream of MOMP is required for its pro-inflammatory role in apoptosis, and we find experimental conditions of caspase activity levels in which GSDME cleavage and STING activation can happen simultaneously following apoptosis induction. We thus propose that partial caspase activation could be a realistic scenario under conditions of sublethal MOMP, or even lethal MOMP, in which the activation levels of caspase-3 are sufficient to produce partial cleavage of both GSDME and cGAS, sufficient for MIM permeabilization and mtDNA release, but insufficient for blocking the STING pathway. GSDME levels could then determine mtDNA-induced interferon responses, and thus the inflammatory or non-inflammatory outcome of cell death following apoptosis induction. In line with this, GSDME can dictate the inflammatory outcome of neutrophil death^52^ and promote interferon responses through the mtDNA/cGAS/STING axis during immune checkpoint inhibitor-induced myocarditis^53^. Our findings thus help explain the dual roles of caspase-3 as both pro- and anti-inflammatory and the biological implications of GSDME-N targeting the mitochondrial inner membrane and causing mtDNA release, with broader relevance for our understanding of cell death.

While immune-silent apoptosis is the desired outcome during programmed apoptosis in development or in remission of T cell populations after infection, it is reasonable to conceive that an inflammatory footprint of apoptosis could take place during apoptosis in response to stress, in the context of infection or cancer. The transition from apoptosis to pyroptosis-like death by activating GSDME downstream of caspase-3 illustrates this ^11^. Pathophysiological scenarios in which there is BAX/BAK-dependent MOMP, which activates caspases, causing mtDNA-induced STING signaling in absence of caspase inhibition have already been reported in minority MOMP during senescence, in the context of radiotherapy against breast cancer tumors, and more recently by VDAC depletion in tumors, which induces BAK-dependent cell death, STING-mediated interferon responses and inflammation dependent on mtDNA release, also in the absence of caspase inhibition^54–56^. In these studies, it remained unclear how MIM permeabilization and mtDNA is mediated downstream of BAX/BAK-induced MOM permeabilization. Our study sheds new light on this question.

Our results do not disentangle whether mtDNA, or other mitochondrial content like mtRNA^57^, is released through GSDME-N pores or through other openings that may form at the MIM as a consequence of GSDME-N-mediated MIMP. The relationship between MIM herniation, permeabilization, and mtDNA release is not fully understood. We propose that MIM extrusion and MIM permeabilization are distinct, yet interrelated processes. In Jenner et al. ^58^, we found that MIM swelling, likely following permeabilization, contributes to the extrusion of the inner membrane. Using membrane dynamics simulations, we found that cristae unfolding during apoptosis also contributes to the extrusion process independently of MIM permeabilization. This is consistent with the partial reduction in MIM extrusion reported here with the GSDME KO cells and could explain why GSDME depletion only partially reduces, rather than completely abolishes, MIM extrusion. Once extruded through BAX/BAK pores, the MIM may become susceptible to mechanical stress and rupture. In addition, the ion fluxes and matrix swelling induced by GSDME-N pores might trigger the opening of the mitochondrial permeability transition pore or interplay with other mitochondrial components, like VDAC, involved in mtDNA release under certain conditions of mitochondrial stress^59,60^.

Despite the requirement of GSDME for MIMP and mtDNA release under our experimental conditions, we cannot discard the existence of other cellular components capable of MIMP, including other GSDM family members^26^. We also detected GSDME-N assemblies at multiple intracellular localizations, suggesting that GSDME-N might permeabilize other organelles, although the functional relevance remains to be established. In agreement, GSDM-mediated damage in lysosomes was recently proposed^61^. Remarkably, GSDME-N and GSDMD-N presented different rates of membrane permeabilization, depending on lipid composition, and of cellular damage. This, together with differential expression and activation mechanisms^12,62^, suggests that GSDMs likely exhibit distinct pore properties to perform a variety of biological functions, especially in immunity, which remain yet to be defined.

In summary, here we report the structure of the GSDME-N pore complex in the membrane, which uncovers a mechanism for GSDME-N pore formation based on membrane remodeling by the asymmetric insertion of the anchor domain into the accessible leaflet of the bilayer. We identify the anchor domain as a driver for preferential cardiolipin binding, which defines mitochondria targeting. We validate the pore structure in mitochondria and in cells by visualizing GSDME pore-like assemblies at the mitochondrial inner membrane during apoptosis. Finally, we show that GSDME can mediate MIMP and mtDNA release into the cytosol downstream of MOMP. This is associated with cGAS/STING activation and inflammatory responses under caspase inhibition. Our findings link GSDME pores with mitochondrial-driven inflammatory signaling, further establishing Gasdermins as key players in innate immunity.

## Materials and methods

### Constructs

hGSDME-mEGFP, hGSDME-HaloTag, hGSDME-W44A-W46A-HaloTag, hGSDME-cpHalo, AKAP1-mEGFP, AKAP1-mEGFP-Hpep3, puro-leader-aALFAnb-TMD-mEGFP-Hpep3, Tom20-HaloTag, Tom20-HaloTag-AlfaTag, and CV(γ)-HaloTag-AlfaTag were cloned into pSems vector under the control of CMV promoter and used for transient transfection. hGSDME-mEGFP was additionally cloned into pSems vector under the control of PGK promoter to reduce GSDME expression and used for transient transfection. To visualize the localization of GSDME and its activated N-terminal fragment (GSDME-NT), hGSDME was tagged with either mEGFP, HaloTag or cpHaloTag. The tags were inserted into the linker region between the N-terminal (NT) and C-terminal (CT) domains, immediately upstream of the caspase-3 cleavage site. This design ensures that upon proteolytic activation by caspase-3, the fluorescence tag remains covalently attached to the GSDME-NT fragment, while the GSDME-CT domain is released, analogous to our previously described GSDMD construct^21^. Consequently, the observed fluorescence signals report the localization of both the full-length protein and the activated GSDME-NT. The N-terminal domains of hGSDMC and hGSDME were cloned in a pEGFP-N2 vector and were used for transient expression in mammalian cells. Full length hGSDME and mGSDMD bearing a C-terminal 8xhis tag were cloned in a pET21a vector and was used for recombinant protein production from E. coli. pET21b-caspase-3 plasmid was ordered from addgene (#90087).

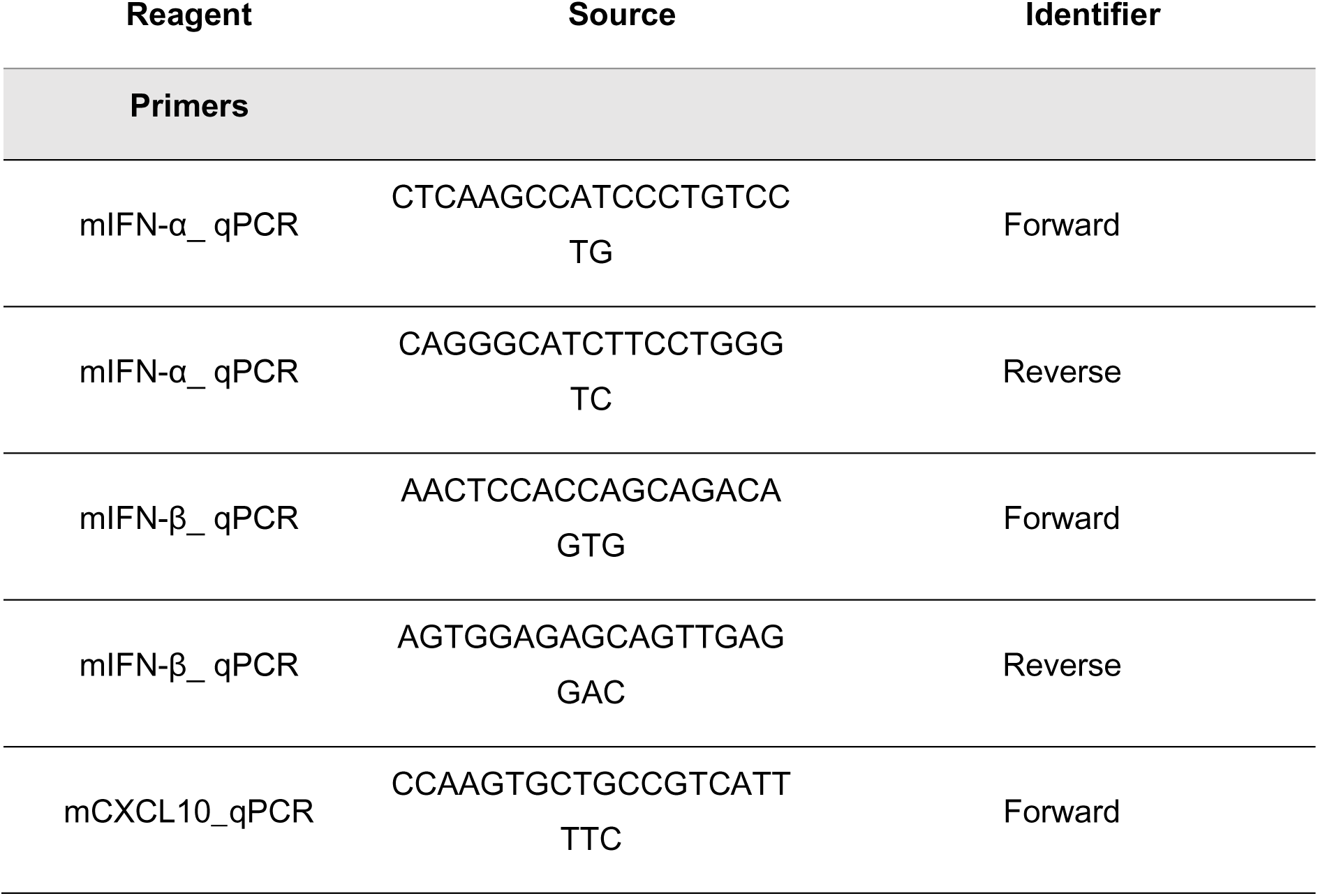
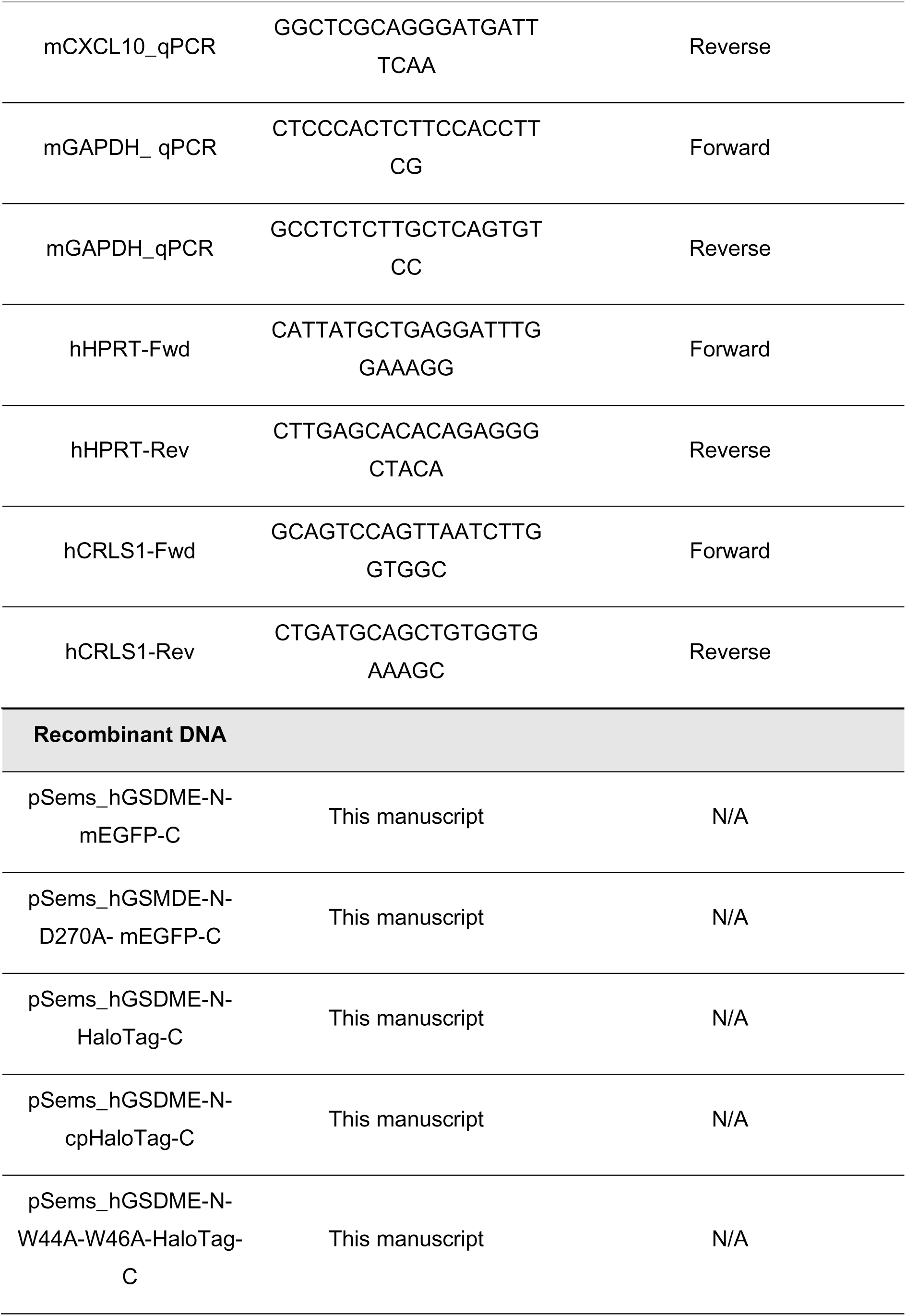
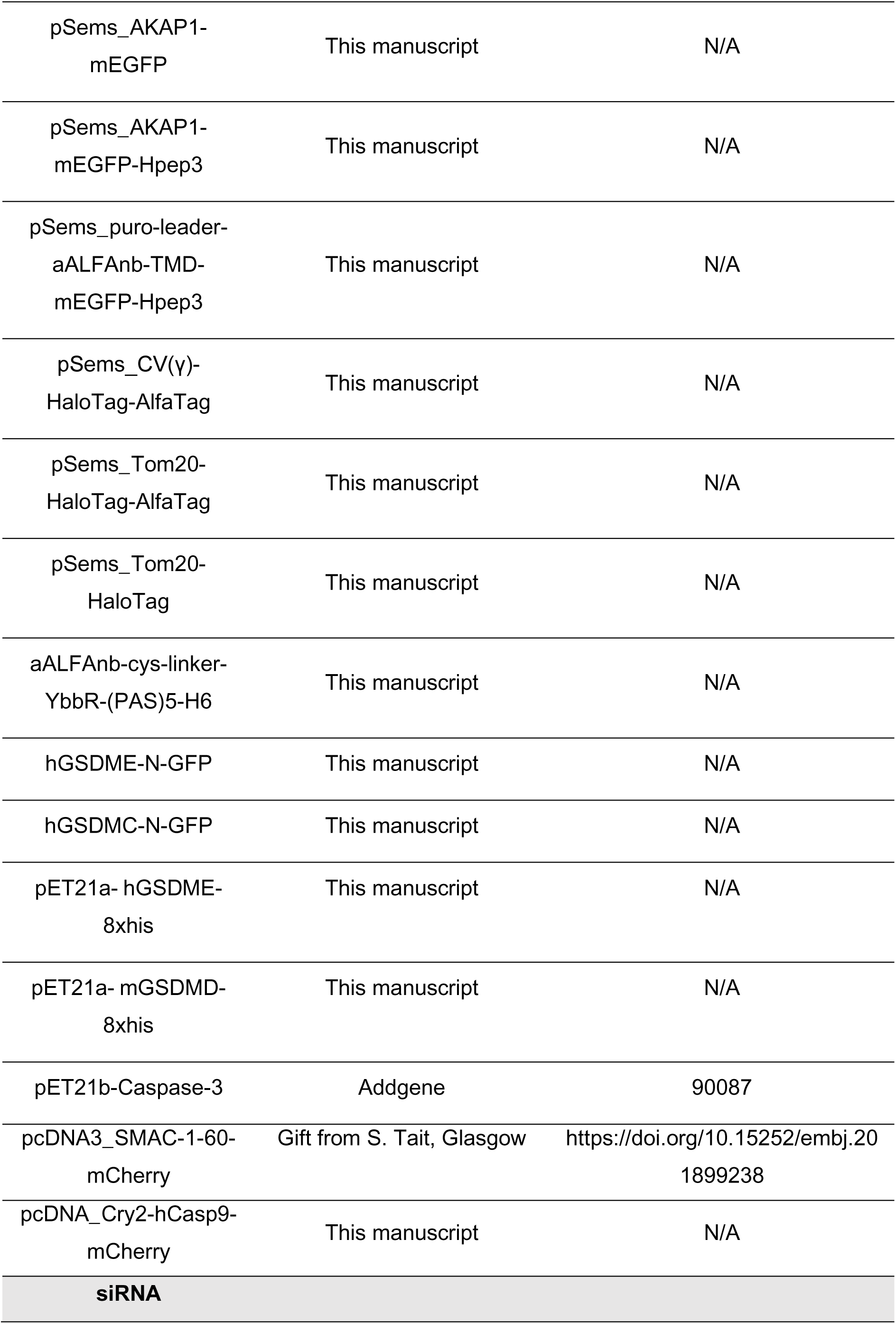
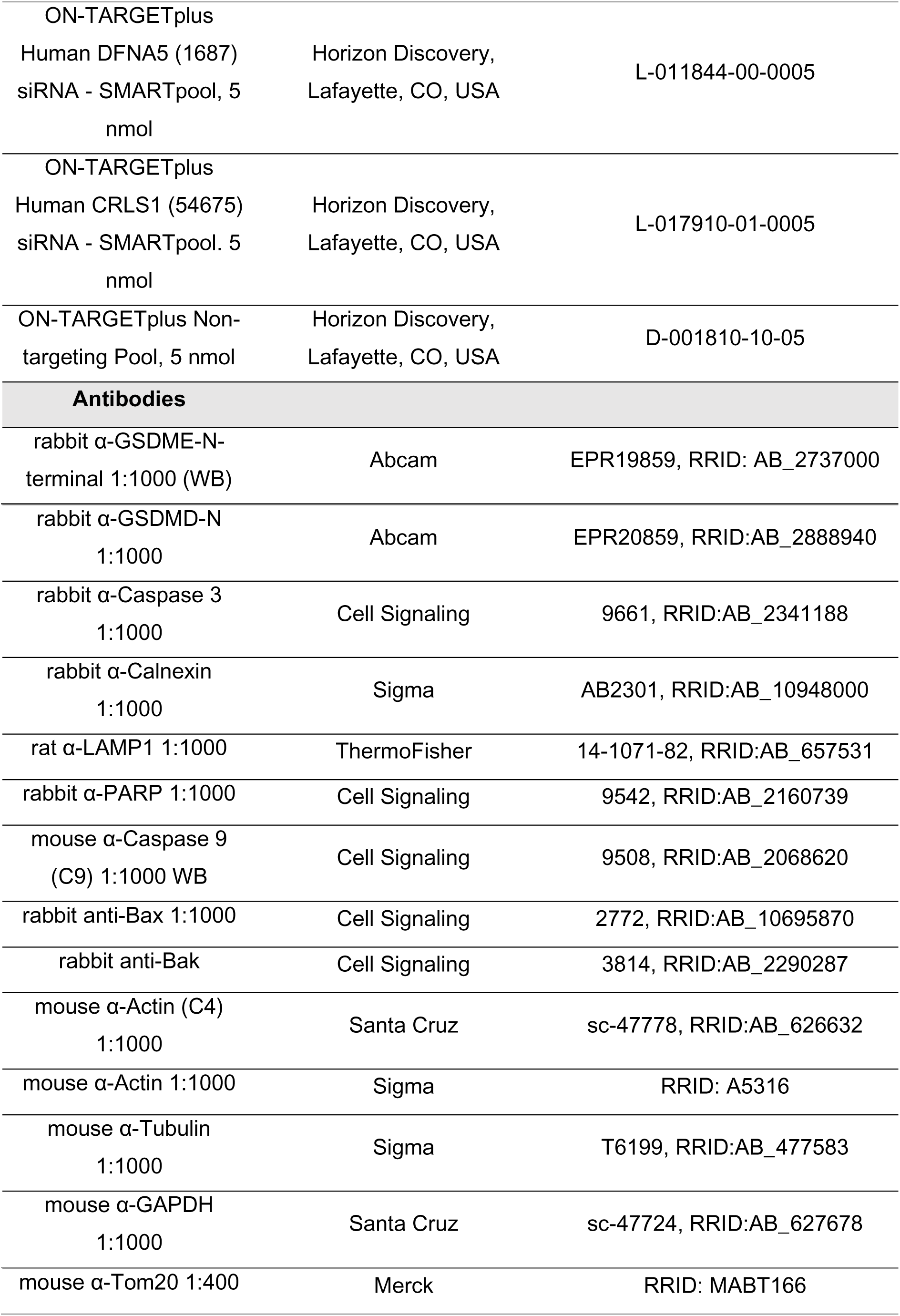
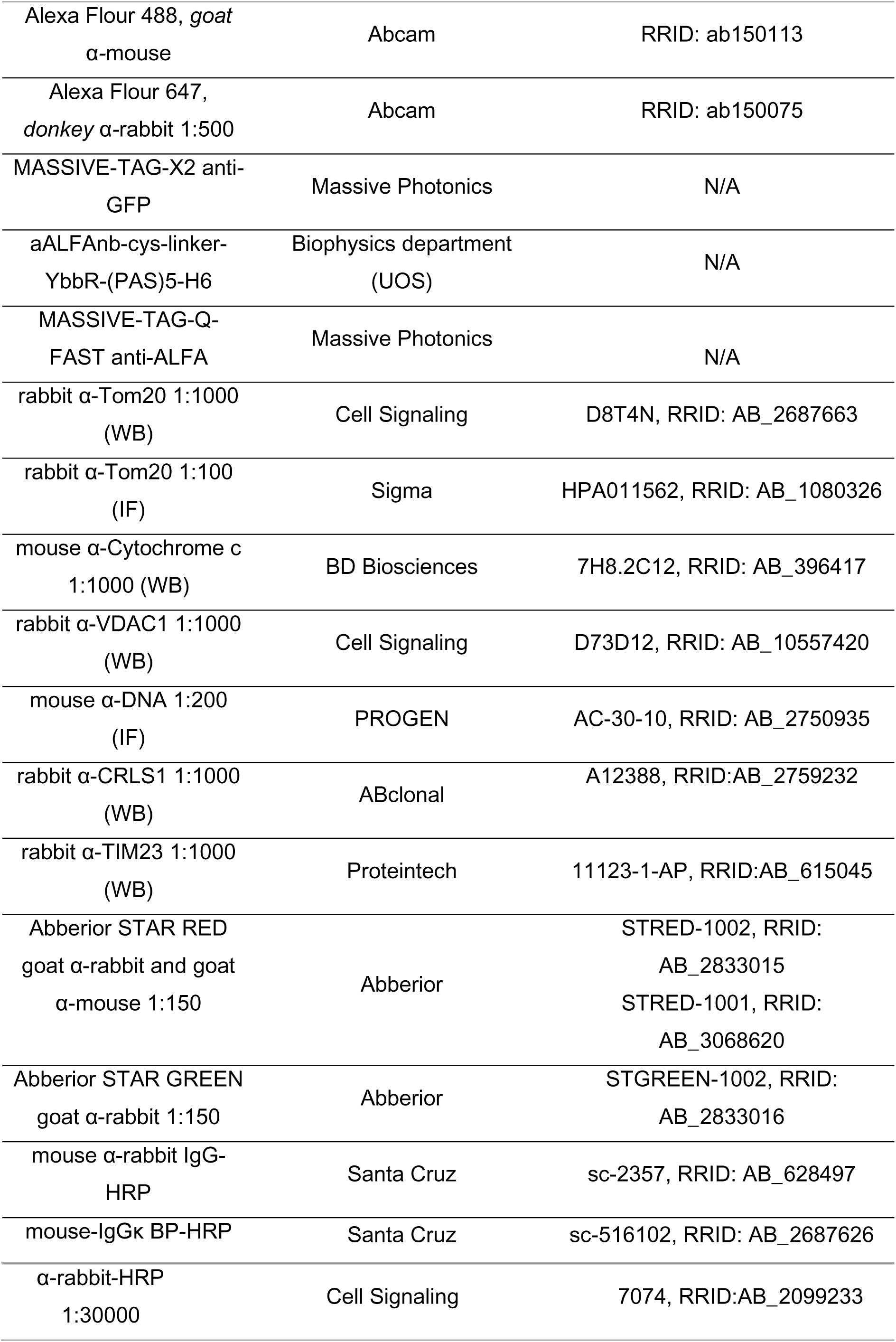
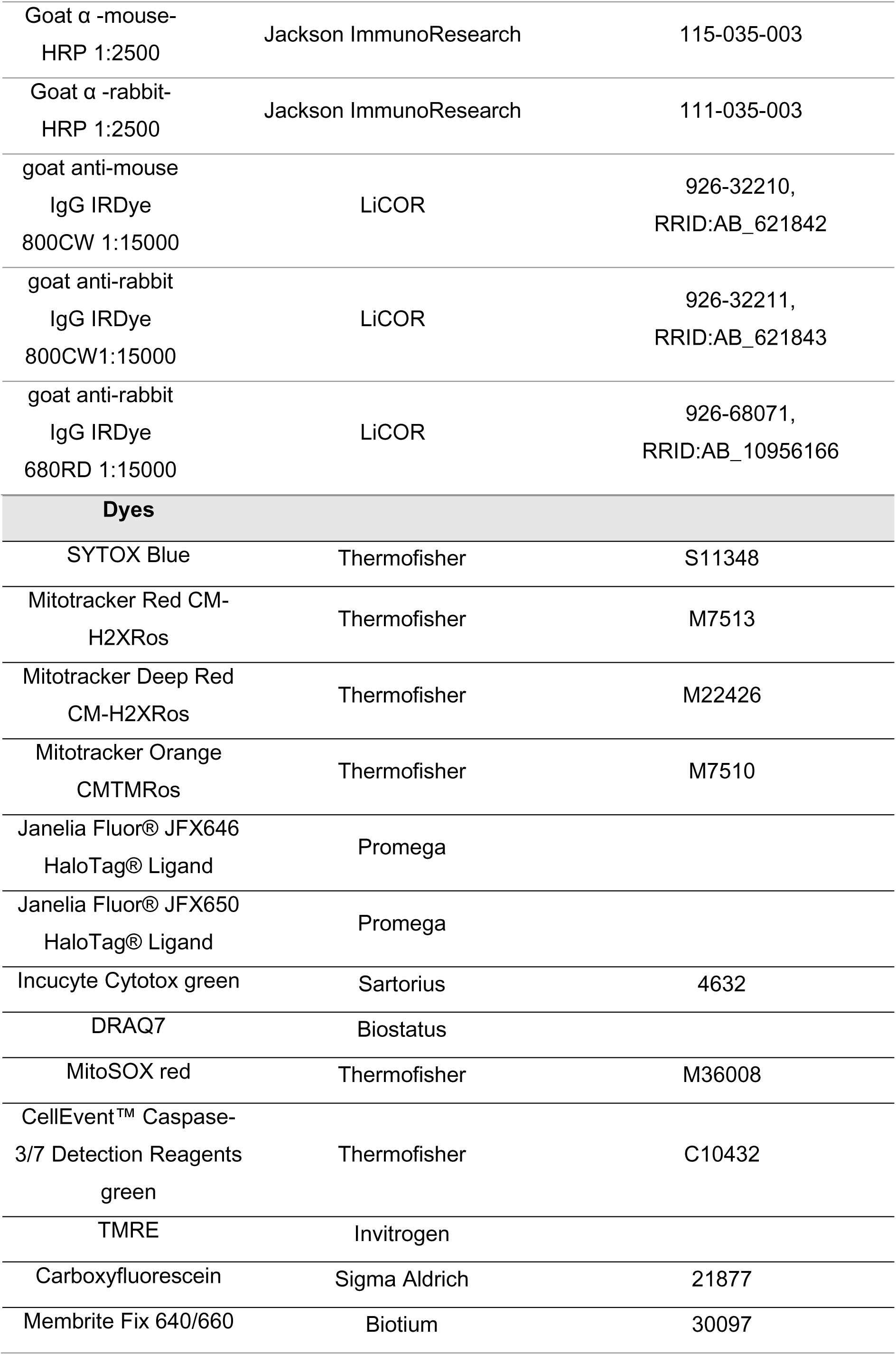
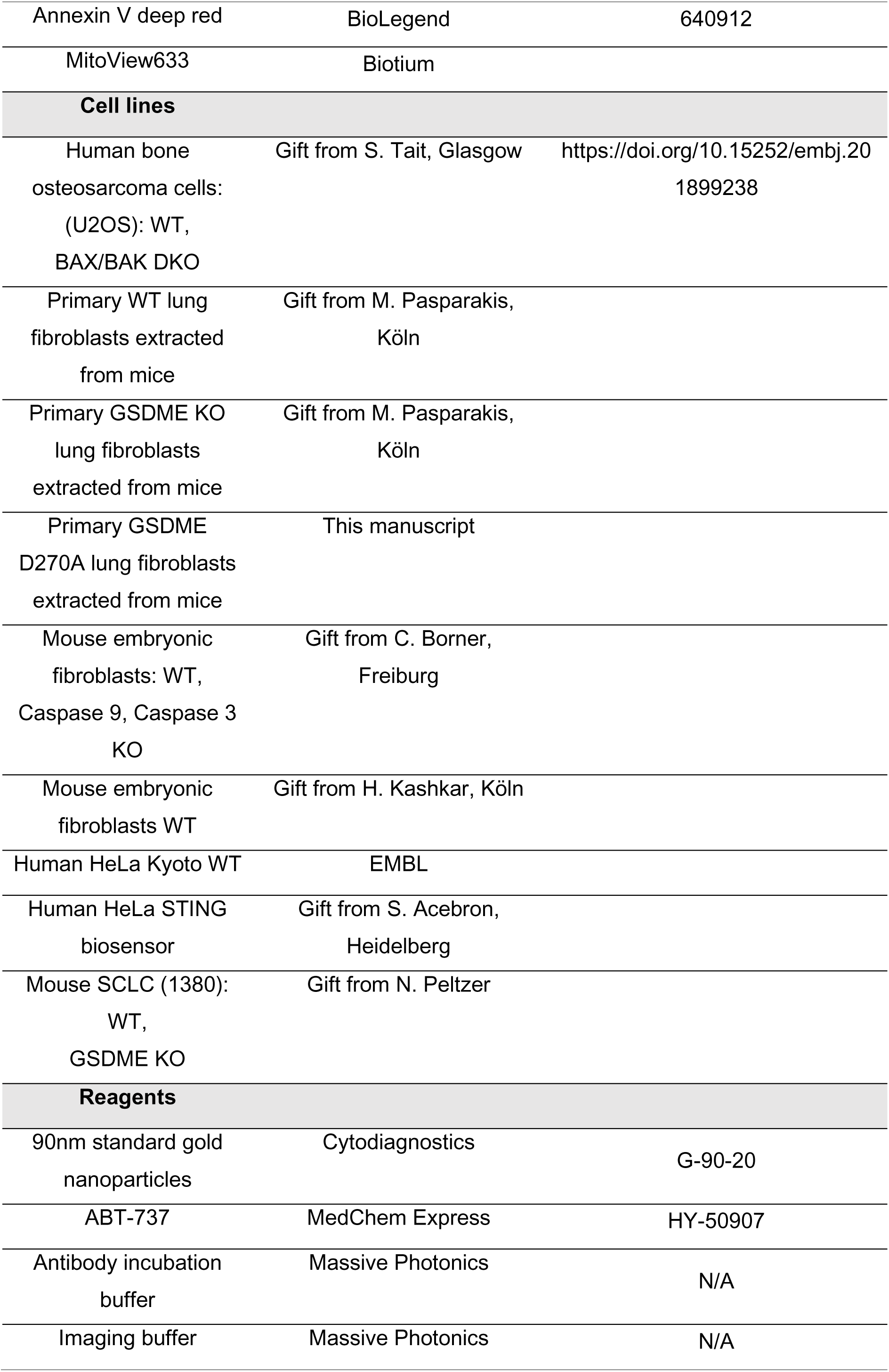
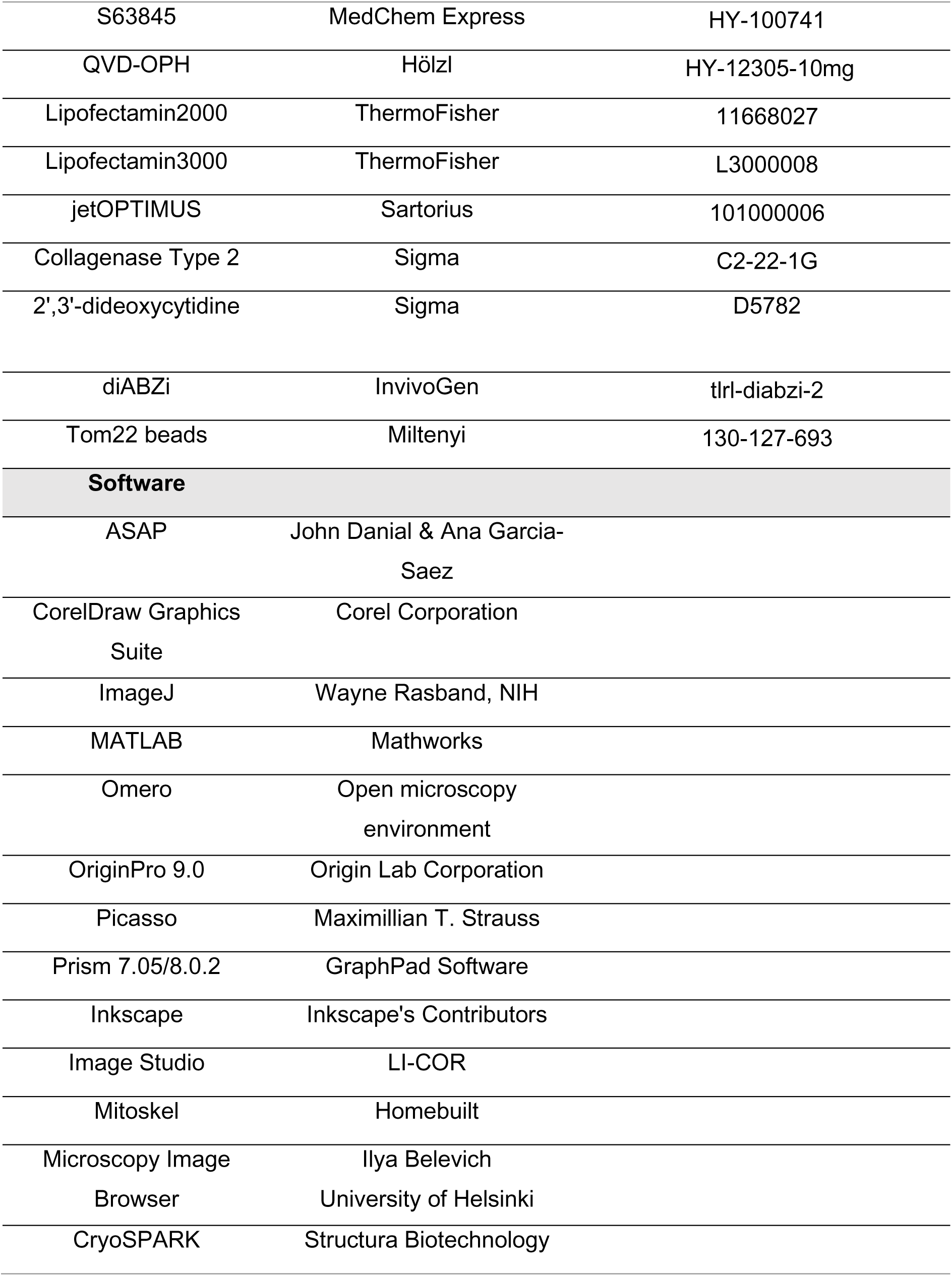

### Cell culture and isolation of primary lung fibroblasts

Unless otherwise indicated, human osteosarcoma U2OS, HeLa and MEF cells of all genotypes were cultured in DMEM supplemented with 10% FBS and 1% penicillin/streptomycin (Invitrogen, Germany). Primary lung fibroblasts were isolated from 10–32-week-old male or female *Gsdme^D270A^* or *Gsdme^-/-^* mice on a C57BL/6N background. Wild-type (WT) C57BL/6N mice were used as controls. Mice were housed in the SPF animal facilities of the Institute for Genetics and the CECAD Research Center of the University of Cologne with 12-hour light/dark cycles. All animal procedures were approved by responsible local authorities (Landesamt für Natur, Umwelt und Verbraucherschutz Nordrhein-Westfalen, Germany; ethical committee of Ghent University, Belgium). Lung fibroblasts of all genotypes were extracted from whole lung tissue using Collagenase Type 2 and were cultured in GlutaMax high glucose DMEM (Gibco) supplemented with 10% FBS and 1% penicillin/streptomycin (Invitrogen, Germany). Mouse 1380 SCLCs wt and *Gsdme* KO were cultured in RPMI supplemented with 10% FBS and 1% penicillin/streptomycin (Invitrogen, Germany). All cells were cultured at 37 °C with 5% CO2. All cell lines used in this study were regularly tested for mycoplasma.

For advanced microscopy, U2OS cells were cultured in MEM Eagle (PAN Biotech, P04-09500) supplemented with 10% FBS, 1% non-essential amino acids (NEAA – Pan Biotech, P08-32100) and 1% HEPES buffer (Pan Biotech, P05-01100). Cells were transfected at 80% confluence via calcium phosphate co-precipitation for 16 h. Two days before microscopy experiments, cells were detached by Trypsin/EDTA treatment (Capricorn Scientific, TRY-1B10) and seeded in microscopy supports.

### Immunoblotting

Samples were lysed in RIPA buffer (0.5% sodium deoxycholate, 150 mM NaCl, 1% (v/v) Triton™ X-100, 0.1% SDS, 50 mM Tris, pH 8.0) supplemented with Complete Protease Inhibitor Cocktail (Roche) and boiled in sample buffer (62.5 mM of Tris/HCl pH 6.8, 10% glycerol, 2% SDS, 0.005% ß-Mercaptoethanol, 0.01% bromophenol blue) for 5 min at 95°C. The protein concentration was determined with BCA Protein assays, for samples of the differential GSDME-GFP expression levels. Protein samples were separated by either by 12% or 4-15% gradient (BioRad) SDS-PAGE reducing gels (5% stacking gels) and transferred onto nitrocellulose membranes (Amersham) or PVDF membranes (Amersham) for caspase blots using the Turboblot system (BioRad) at 1-1.3 mA for 15-30 min or using a trans-blot SD semi-dry transfer cell (BioRad) at 0.2 A for 1h. Blots were blocked with 3% w/v BSA or 5% w/v non-fat dried milk and incubated overnight at 4°C with primary antibodies diluted in 2-3% BSA in TBS-T (TBS + 1% Tween-20; see table), probed with secondary antibodies diluted in TBS-T (see table) for 1h at RT. For HRP antibodies, blots were developed using SuperSignal™ West Pico PLUS or FEMTO PLUS chemiluminescent substrate (Thermo Scientific) and fluorescent antibodies were imaged on a BioRad Gel doc, ChemiDoc or a LiCor Odyssey system or an AZURE 600 gel imager (Biozym).

### Subcellular fractionation and cytochrome c release from isolated mitochondria

In order to track the translocation of GSDME-N to mitochondria 2.5-3x10^6^ cells were seeded per 10 cm dish and treated with 1 µM ABT-737 and 1 µM S63845 (U2OS cells), 5 µM per drug (MEFs, primary lung fibroblasts) or 10 µM per drug (SCLCs) to induce apoptosis for indicated time points. The supernatant (SN) of the cells was spun down to collect detached cells and the cells were harvested via scraping in mitochondria stabilization buffer (MSB; 250 mM Sucrose, 10 mM Tris/HCl pH 7.4; 1 mM EDTA) with freshly added protease inhibitor cocktail (Roche). The cells were passed through a 30 G needle (∼25 times) when used for protein translocation assays. Debris and unbroken cells were removed by centrifugation at 800 g for 10 min at 4°C (2x). The SN was spun down at 10,000 g for 15 min at 4°C to pellet crude mitochondria. The whole cell extract (WCE), mitochondria, and the SN fractions were separated by SDS-PAGE and analysed via immunoblot.

To further fractionate MEF WT cells, the SN of the mitochondria pellet was spun down at 30,000 g for 30 min at 4°C to pellet an ER-rich fraction. Then the SN was spun down at 100,000 g for 90 min at 4°C to pellet the microsome and separate it from the cytosol. Ultracentrifugation pellets were resuspended and vortexed in in urea lysis buffer (8M urea, 150 mM NaCl, 50 mM Tris-HCl pH 7.4, and 5mM EDTA) containing freshly added protease inhibitors (Roche) followed by an equal volume of 2X RIPA (300 mM NaCl, 1% v/v Triton X, 1% v/v NP40, 1% w/v sodium deoxycholate, 0.2% w/v SDS, and100 mM Tris pH 8), also containing freshly added protease inhibitors (Roche). In the meantime, the crude mitochondrial pellet was resuspended in 1 mL MSB and incubated rotating with 25 µl of Tom22 beads (Miltenyi, Germany) for 1h at 4°C. LS columns (Miltenyi, Germany) were attached to a QuadroMACS™ Separator (Miltenyi, Germany) and equilibrated with 2 mL protein extraction buffer (PEB; 2 mM EDTA; 0.5% BSA in PBS pH 7.2). The mitochondria/Tom22 beads suspension was added to the LS column which was washed 3x with 2 mL PEB buffer. Afterwards, the columns were removed from the magnet and the beads were eluted 2x with 2 mL PEB buffer into 15 mL tubes using the plunger of the column. All fractions were separated by SDS-PAGE and analysed via immunoblot.

To assess the role of cardiolipin in GSDME recruitment to mitochondria-enriched fractions, U2OS WT cells were reverse transfected with 40 nM siNT (non-targeting siRNA) and siCRLS1 using Lipofectamin2000 for 72 h. Afterwards, apoptosis was induced with 1 µM ABT-737 and 1 µM S63845 in McCoy medium containing 10% delipidated FBS (Bio&SELL, FCS.LFS.0100) for 2 h, which was followed by crude mitochondria isolation and immunoblotting. For the cytochrome c release assay, the cells were harvested via trypsinization, cell pellets resuspended in isolation buffer (300 mM Trehalose, 10 mM KCl, 1 mM EDTA, 10 mM HEPES–KOH pH 7.7 and 0.1% BSA) and mechanically broken using a glass homogenizer on ice (∼30 strokes). Debris and unbroken cells were removed by centrifugation at 800 g for 10 min at 4°C (2x). The SN was spun down at 10,000 g for 15 min at 4°C to pellet crude mitochondria. The cytosolic fraction was removed, the protein concentration was determined by Bradford assay (BioRad) and 35 µg of isolated mitochondria were incubated with Caspase 3 (50-100 nM), GSDME (2 µM) or cBID (10 nM) in release buffer (300 mM trehalose, 80 mM KCl, 1 mM EDTA, 10 mM HEPES–KOH pH 7.7 and 0.1% BSA) for 30 min at 30°C. For combined treatments of Caspase 3 and GSDME, GSDME was pre-activated with Caspase 3 for 10 min at 37°C. Afterwards, mitochondria were pelleted by centrifugation at 14000 g for 10 min at 4°C. The mitochondria and the supernatant fractions were separated by SDS-PAGE and analysed via immunoblot.

### Recombinant protein expression and purification

For the production of recombinant human GSDME and mouse GSDMD, BL21-(DE3)-RIPL E. coli competent cells (Agilent Technologies) were transformed with the plasmids and were grown in LB media containing Ampicillin and Chloramphenicol. Protein expression was induced at OD = 0.8, using 0.5 mM isopropyl-β-D-thiogalactoside (IPTG) for 3 h at 25 °C. The bacteria were then harvested by centrifugation, resuspended in Lysis buffer (50 mM Tris, 150 mM NaCl, 1 mM DTT, pH 7.5) and were then lysed by sonication. Afterwards, the lysate was cleared by centrifugation at 20,000 g for 1 h. The supernatant was incubated with Ni-NTA agarose beads (Qiagen) and was eluted with 250 mM imidazole. The eluted fractions were then concentrated and injected on a Superdex 200 Increase 10/300 GL Size exclusion column (Cytiva). Protein purity was examined by SDS–polyacrylamide gel electrophoresis. For the production of caspase 3, BL21-(DE3)-RIPL E. coli competent cells (Agilent Technologies) were transformed with the plasmid and were grown in LB media containing Ampicillin and Chloramphenicol. Protein expression was induced at OD = 1, using 0.2 mM IPTG for 5 h at 30 °C. The bacteria were then harvested by centrifugation, resuspended in Lysis buffer (50 mM Tris, 100 mM NaCl, 20 mM Imidazole, pH 8) and were then lysed by sonication. Afterwards, the lysate was cleared by centrifugation at 20,000 g for 1 h. The supernatant was incubated with Ni-NTA agarose beads (Qiagen) and was washed with washing buffer (50 mM Tris, 500 mM NaCl pH 8) and then eluted with 250 mM imidazole. The eluted fractions were then concentrated and dialyzed in (50 mM Tris, 100 mM NaCl, pH 8). Protein purity was examined by SDS–polyacrylamide gel electrophoresis.

### Liposome permeabilization assay

Lipids were purchased from Avanti Polar Lipids, and were dissolved in chloroform and mixed with the desired molar ratios. Chloroform was then evaporated under vacuum for 3 h. Large unilamellar vesicles (LUVs) were prepared using the extrusion method as described before^63^. Briefly, the lipid film was hydrated with 80 mM solution of Carboxyfluorescein, pH 7 for a final lipid concentration of 5 mg/mL followed by five cycles of freezing and thawing. The lipid solution was then extruded through a polycarbonate membrane with a pore size of 100 nm using glass syringes. Carboxyfluorescein-loaded LUVs were separated from free Carboxyfluorescein using Sephadex-G50 beads and the lipid concentration was adjusted to 100 μM. Full-length hGSDME and mGSDMD were incubated with caspase3 and caspase4 respectively at 37°C for 10 min to obtain the active hGSDME-N and mGSDMD-N. Then, LUVs were incubated with serial dilutions of the active protein in a 96-well plate and Carboxyfluorescein release was monitored by fluorescence emission at 520 nm with excitation at 490 nm for 1 h using a microplate reader (Spark, TECAN). 0.1% Triton X-100 was used as a positive control (100% Carboxyfluorescein release) and the percentage of Carboxyfluorescein release was calculated as follows:

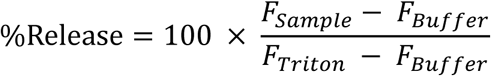

### Liposome co-sedimentation assay

Experiments were adapted from ^64^. Lipid films were resuspended in (20mM HEPES, 140mM NaCl, 1mM EDTA, pH=7) to give a final concentration of 0.5 mg/mL, followed by five cycles of freezing and thawing. Full-length hGSDME and mGSDMD were incubated with caspase3 and caspase4 respectively at 37°C for 10 min to obtain the active hGSDME-N and mGSDMD-N, and the active proteins were incubated with the LUVs for 30 min at 25 °C. Afterwards, centrifugation at 35,000 g for 30 min was used to pellet down the liposomes. The pellet and supernatant were fractionated by SDS-PAGE and stained with Coomassie. Intensity of protein bands was quantified using Image Studio software.

### Two-colour GUV permeabilization assay

Giant unilamellar vesicles (GUVs) were prepared by electroformation. Lipid mixtures were prepared in chloroform containing either 80 mol% 1,2-dioleoyl-sn-glycero-3-phosphocholine (DOPC) and 20 mol% bovine heart cardiolipin (CL), mimicking the mitochondrial membrane, or 50 mol% DOPC, 30 mol% 1,2-dioleoyl-sn-glycero-3-phosphoethanolamine (DOPE), 10 mol% cholesterol, and 10 mol% 1,2-dioleoyl-sn-glycero-3-phospho-L-serine (DOPS), mimicking the plasma membrane. Mitochondrial and plasma membrane GUVs were labelled with 0.5 mol% DiI and DiO, respectively. For electroformation, 3 μL of lipid solution were deposited onto each of two platinum wires and dried completely. The chamber was assembled and filled with 300 mM sucrose. An alternating electric field of 1.5 V at 10 Hz was applied for 2 h, followed by a maturation step at 1.5 V and 2 Hz for 40 min. Electroformed GUVs were harvested and stored at room temperature until use. Both GUV populations were mixed immediately before imaging. Alexa Fluor 655 (1 μM) was included in the external PBS as a membrane-impermeable tracer to monitor membrane permeabilization. GSDME-N, generated by caspase-3 cleavage, was added to final concentrations of 0.3 or 1 μM. Samples were incubated for 5 min before image acquisition. Time-lapse imaging was performed on a Zeiss LSM 980 confocal microscope equipped with a 60× objective. For each experiment, 4 × 5 tiled images containing both GUV populations were acquired. Images were recorded for 80 min to monitor Alexa Fluor 655 influx into individual vesicles. Image analysis was performed using GUVprofiler, available at https://github.com/jaufdermauer/GUVprofiler. To account for baseline dye accumulation observed in the absence of protein, permeabilization values were normalized to the corresponding negative control according to:

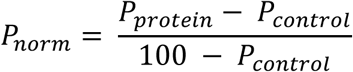

Where *P_protein_* denotes the permeabilization measured in the presence of GSDME-N and *P_control_* denotes the corresponding value obtained from the negative control. Normalized permeabilization values were expressed as percentages by multiplying *P_norm_* by 100. At least two independent biological replicates were performed for each condition, each including a minimum of two technical replicates. Identical analysis parameters were applied to all experimental conditions.

### Optogenetics experiments

BAX/BAK double-knockout (DKO) U2OS cells were seeded at a density of 50,000 cells per well in removable eight-well chambers (catalog no. 80826; ibidi) fitted with glass coverslips suitable for confocal microscopy (VWR). After 6 h, cells were co-transfected with 150 ng of hGSDME–GFP or hGSDME-D270A–GFP and 150 ng of Opto-Caspase-9–mCherry. Cells were incubated for 16 h in the dark before imaging. Optogenetic experiments were performed using an LSM 980 confocal laser-scanning microscope equipped with Airyscan 2 (Carl Zeiss Microscopy) and operated using ZEN 3.5 software (Zeiss). Images were acquired using a 63×/1.2 NA water-immersion C-Apochromat objective. Diode lasers with wavelengths of 488, 561, and 638 nm, each with a maximum output of 30 mW, were used together with GaAsP and transmitted-light photomultiplier tube (T-PMT) detectors. The emission filters used to collect images were 490–534 nm, 410-623 nm, and 641–694 nm, respectively. During imaging, cells were maintained at 37°C and 5% CO₂ using an integrated incubation module (Carl Zeiss Microscopy). The following laser settings were used: 45 µW for the 488 nm laser (GFP), 95 µW for the 561 nm laser (mCherry) and 11 µW for 639 nm laser (MitoTracker far red). Brightfield images were acquired using the 561 nm laser with 95 µW and a transmitted light (T-PMT) detector. Photoactivation was performed by acquiring GFP fluorescence using the 488 nm laser at 10-second intervals until the cell either died or moved out of the imaging field. To assess mitochondrial translocation, videos were analyzed throughout the entire acquisition period, and quantified foci were annotated as ROIs in Fiji.

### Spinning disk confocal microscopy

For monitoring the kinetics of mitochondrial fragmentation and SYTOX Blue uptake after apoptosis induction, U2OS cells were transfected with pSems-hGSDME-mEGFP and pSems-Tom20-HaloTag. A day before the experiment, cells were seeded into an 8-well plate (80821, ibidi), and directly before the experiment 50 nM JFX646 dye and 1 µM SYTOX Blue were added. Cells were imaged before apoptosis induction and every 30 min after induction using 1 µM ABT-737 and 1 µM S63845 with a fully motorized inverted spinning disc microscope based on a Zeiss Cell Observer.Z1 and a CSU-X1 spinning disc unit (Yokogawa) with a 63x, oil immersion objective (Alpha Plan-Apochromat, NA 1.46). The microscope was equipped with a home-built incubation chamber and a PID-controller heater (The Cube, Live Imaging Services). Samples were imaged at 37°C in a humidified (Zeiss humidifier module S1) 5% CO2 atmosphere (Zeiss CO2 module S1), a motorized XYZ-stage (PZ-2000 XYZ, Applied Scientific Instrumentation and a Definite Focus System (Zeiss). The morphology of the cells was monitored by BF. To monitor PM permeabilization, SYTOX Blue was excited with a 405 nm diode laser (max. p. 50mW), while to monitor mitochondrial fragmentation, the mitochondrial marker Tom20-HaloTag was checked with a 635 nm diode laser (max. p. 30mW). hGSDME-mEGFP expression and localization was checked with a 488nm optically pumped semiconductor laser (max. p. 100mW). Appropriate filters were used before detection by a Hamamatsu OCA Flash V3. Imaging was performed with the software Zeiss Zen 2.6. Analysis of mitochondria fragmentation was done using ImageJ and the plugin Mitochondria Analyzer (Chaudhry 2020). The settings for thresholding in the Mitochondrial Analyzer plugin were adjusted, by setting the block size to 1.05 microns and C-value to 5, before analysis with default settings.

### Airyscan high-resolution microscopy

To observe GSDME localization at mitochondria apoptosis U2OS cells were transfected with pSems-hGSDME-HaloTag or pSems-hGSDME-W44A-W46A-HaloTag and pSems-AKAP1-mEGFP or with pSems-PGK-hGSDME-mEGFP and Tom20-HaloTag. HeLa cells were transfected with pSems-hGSDME-HaloTag and pSems-AKAP1-mEGFP. Cells were seeded into an 8-well plate (80821,ibidi), 100 nM Mitotracker Red (Thermofisher) and 50 nM JFX650 dye were added. Apoptosis was induced for varying timepoints using 1 μM ABT-737 and 1 μM S63845 and cells were then fixed with 4% PFA for 20 min. Samples were imaged with a fully motorized Zeiss Cell Observer 7 confocal laser scanning microscope (cLSM) 880 with a fast Airyscan detector. Imaging was done with a C Plan-Apochromat 63ximmersion oil objective, a xy scanning stage SCAN IM 130x100, and a high precision z-axis piezo stage. DIC was used to observe the morphology of the cells. The localization of hGSDME-HaloTag or the outer mitochondrial membrane marker Tom20-HaloTag were checked by a 633 nm helium-neon laser. The outer mitochondrial membrane marker AKAP1-mEGFP or PGK-GSDME-mEGFP were excited by a 488 nm argon-ion laser. The inner mitochondrial membrane marker Mitotracker Red (ThermoFischer) was observed by a 561 nm diode-pumped solid-state laser. Z-stacks were acquired for every timepoint. Imaging and image analysis were done by the Zen2.1 software. For assessing the co-localization of GSDME at mitochondria Z-stacks were transformed into maximum-intensity projections in Fiji. Colocalization calculations were performed using Fijis BIOP JACoP plugin according to the developers’ instructions. Intensity Line-Plots were done with Fiji.

### DNA-PAINT

For visualizing the structures of GSDME at mitochondrial membranes U2OS cells were transfected with pSems-hGSDME-mEGFP and either pSems-Tom20-HaloTag-AlfaTag or pSems-ComplexV-HaloTag-AlfaTag. A day before the experiment cells were seeded onto glass coverslips. Directly before the experiment 50 nM JFX646 dye was added. Apoptosis was induced using 1 µM ABT-737 and 1 µM S63845. Cells were fixed at 0 min and 150 min.

For DNA-PAINT sample preparation, chips with fixed cells were reduced with 1 mg/mL sodium borohydride at RT for 7 min and washed 4 times for 5 min with 1 mL washing buffer (Massive Photonics). Cells were permeabilized for 2h with 3 % BSA and 0.25 % Triton X-100, and then incubated with 50 nM of the anti-GFP nanobody (MASSIVE-TAG-X2 anti-GFP, massive Photonics),10 nM Alfa-nanobody (aALFAnb-cys-linker-YbbR-(PAS)5-H6) or 50nM of the anti-ALFA nanobody (MASSIVE-TAG-Q-FAST anti-ALFA) dissolved in antibody incubation buffer (Massive Photonics) at 4°C overnight or at RT for 2h. After incubation, samples were washed with 1 mL washing buffer and treated with fiducial markers by using a 1:1000 dilution of 90 nm gold nanorods (Cytodiagnostics, G-90-20). After incubation for 5 min at RT, samples were washed with washing buffer and imaging buffer (Massive photonics) to remove unspecific bound nanobodies. Before imaging, samples were treated with 250-500 pM Cy3b-conjugated imager strand-3 (Massive Photonics) dissolved in the imaging buffer and after imaging, washed five times and treated with the second imager Cy3b-conjugated imager strand-3 (Massive Photonics). DNA-PAINT experiments were performed using an inverted Olympus IX-81 microscope equipped with a motorized xy-stage (IM 120x80, Märzhäuser), a motorized quad-line TIR-illumination condenser (cellTIRF-4-Line, Olympus), a 100x oil immersion objective (UAPON 100x TIRF, NA 1.49, Olympus), an incubator with temperature control (TempController 2000-2, CellVivo) and a CO_2_ controller (CO_2_ controller 2000, CellVivo). The signals were monitored by a 488 nm, a 561 nm, and a 640 nm diode-pumped solid-state lasers (Olympus). Laser lines were filtered by clean-up filters: BrightLine HC 482/18 for 488nm, BrightLine HC 561/14 for 561 nm, and BrightLine HC 640/1 for 640 nm (Semrock). Fluorescence emission was filtered by a bandpass filter (BrightLine HC 446/523/500/677) and single bandpass filters for each channel (BrightLine HC 390/40, 482/18, 561/14 and 640/14, Semrock) before detection with a sCMOS camera (ORCAFlash 4.0 V3, Hamamatsu).

For imaging, the temperature was kept at 27°C, and buffer evaporation was prevented by humidification. DNA-PAINT laser was set to HiLo mode at an angle optimized for mitochondrial illumination. The imagers were excited with 561 nm lasers adjusted to a power density of approx. 100W/cm2 in the focal plane stabilized with a hardware autofocus system (IX2-ZDC2, Olympus). 35,000-50,000 frames of each recording were acquired with 150 ms exposure time (for fast alfa variant 50 ms exposure time) and 2x2 pixel binning.

### DNA-PAINT data processing

Each collected dataset was processed, after the removal of the first 5,000-10,000 frames to reduce unspecific signal and drift, with Picasso software suite^42^. *Picasso Localize* was used to identify and localize DNA-PAINT signals. The box size was set to 5 px, and the unwanted background signal as well as the signals with a low signal-to-noise ratio were filtered by adjusting the min net gradient to 2500-5000. Photon conversion parameters used for the analysis were set at: EM Gain = 1, Baseline = 400, Sensitivity = 0.46, Quantum efficiency = 0.9, and Pixel size =130. *Picasso Filter* was used for filtering the fitted signals for localization precision with a threshold of 0.05-0.1 px /6 nm. *Picasso Render* was used for the drift correction of the dataset by cross-correlation and with gold nanorod fiducials. Localizations were linked with a localization precision threshold of 0.05 px. Super-resolved structures were picked with a peak diameter of 0.5 px ± 1 and filtered for specific structures with a frequent sampling pattern with the “pick similar” function.

After processing, 50 random overview images were saved with a zoom of 200 as 8-bit tiffs for the final structural classification with the software Automated Structures Analysis Program (ASAP)^65^. This software allowed the identification of structures by “connectivity” by setting the following parameters: threshold method = fixed, threshold multiplier = 0.6, no cleaner, identification range size = 400-20000. A radial profile analysis with a max. ring size of 10 px was done on the exported files with a pixel size of 0.65 nm. The software was then trained to automatically classify the identified super-resolved GSDME structures into 4 groups based on the shape of the structure: rings, arcs, lines, and undefined clusters. Quantification of the radius of ring- and arc-like structures was done for GSDME particles combined at the MOM and the MIM.

### High-content analysis system CQ1

To monitor PM permeabilization and SMAC release, U2OS WT and HeLa WT cells were transfected with pSems-hGSDME-mEGFP and pcDNA3-SMAC-1-60-mCherry.. To examine PM permeabilization with lower GSDME expression, U2OS cells were transfected with pSems-PGK-hGSDME-mEGFP and pcDNA3-SMAC-1-60-mCherry. One day before imaging, cells were seeded into 8-well plates (80821, ibidi). Immediately before imaging, cells were incubated with 1 µM SYTOX Blue. Apoptosis was induced with 1 µM ABT-737 and 1 µM S63845 for 5-7 h and images were acquired before treatment, and every 30 min. Imaging was done via the high-content analysis system CQ1 from Yokogawa and using a 40x objective. The samples were imaged at 37°C. SMAC-mCherry was excited with a 561 nm laser, hGSDME-GFP with a 488 nm laser and SYTOX Blue with a 405 nm laser. Brightfield images were acquired simultaneously to monitor cell morphology. The CQ1 software was used for imaging. Image analysis was done with ImageJ. PM permeabilization was quantified by the intensity increase over time of the SYTOX Blue signal using manually defined ROIs in the nucleus. The mean intensity per cell was normalized to the first time point. SMAC release was quantified by manually drawing ROIs around the cytosol. The standard deviation of the mean intensity per cell was normalized to the first time-point.

### Live cell imaging of GSDME-N subcellular distribution

To visualize the distribution of GSDME-N between mitochondria and PM after apoptosis induction, we generated GSDME reporter constructs using the same split-tag design principle as described by Lin et al.^66^ U2OS cells were transfected with the GSDME construct pSems-hGSDME-N-cpHaloΔ2-C, together with the outer mitochondrial membrane marker pSems-AKAP1-mEGFP-Hpep3 and the PM marker pSems-puro-leader-aALFAnb-TMD-mEGFP-Hpep3. One day before imaging, U2OS cells were seeded into 8-well glass-bottom chamber slides (80821,ibidi). Immediately before imaging, cells were labeled with 50 nM HTL-JFx650 and images were acquired before and after dye addition. Apoptosis was induced using 1 μM ABT-737 and 1 μMS63845 and cells were imaged at multiple time points. Live-cell imaging was performed using a fully motorized Zeiss Cell Observer 7 confocal laser scanning microscope (cLSM) 880 with a fast Airyscan detector,a C Plan-Apochromat 63ximmersion oil objective, a xy scanning stage SCAN IM 130x100, and a high precision z-axis piezo stage. The hGSDME-cpHalo labeled with HTL-JFx650 was excited by a 633 nm helium-neon laser. AKAP1-mEGFP-Hpep and TMD-mEGFP-Hpep were excited by a 488 nm argon-ion laser. Z-stacks were acquired for every timepoint. Imaging and image analysis were done by the Zen2.1 software (Zeiss). For quantitative analysis, background was subtracted from the GSDME-cpHalo channel in Fiji using the rolling ball (radius of 50 pixel) and sliding paraboloid. Then Mean Intensity of GSDME at mitochondria vs PM was measured for all timepoints by manually drawing ROIs around mitochondria or PM. Data was normalized to images before HTLJFx650 addition. Fluorescence intensity line profiles were generated in Fiji.

### Grid preparation, cryo-EM data collection, processing, and model building

LUVs (PC:CL 8:2) were incubated with GSDME (+20 nM caspase3) for 40 min. The protein:lipid molar ratio was adjusted to 1:100 or 1:10000. 3 µl of liposomes were applied to glow-discharged QF R2/1 300 mesh grids (Quantifoil Micro Tools, Jena, Germany), blotted for 4 – 5 seconds at 10°C and 100% humidity and plunge-frozen using a Vitrobot (Thermo Fischer). Movies were collected on a KriosG3 microscope (Thermo Fisher Scientific) equipped with a Gatan K3 camera at 105kx magnification corresponding to a pixel size of 0.837 Å. The total dose for the 60 frames of the 3sec movies was 65 e^-^/Å^2^ and the defocus ranged from -1.6 to -2.4 µm. Motion-correction, CTF estimation and manual particle picking were performed in CryoSPARC^67^. Particles were extracted with a box size of 500 pixels (419 Å) and 2D-classified in multiple rounds. Ab-initio reconstruction with C1 and C27-33 symmetry applied. Subsequent heterogenous refinements were performed with C29-31 symmetry. NU-refinement of the best class with imposed C30-symmetry yielded the final maps. The 3D reconstructions with symmetry were sharpened using negative B-factors that were calculated in CryoSPARC. The nominal resolutions were estimated using Fourier-shell correlation (FSC) with the gold standard criterion of 0.143. Initial models were obtained using Alphafold^68^ and individual domains were rigid body fitted into the map and manually corrected in Coot^69^. Supplementary Table 1 summarizes data acquisition and refinement statistics. Figures were created with ChimeraX^70^.

### Cryo-ET of isolated mitochondria

The cultivation of *Polytomella* cells and isolation of its mitochondria were performed as previously described by Blum et al. (2019)^71^. At the end of the isolation procedure, mitochondria were flash-frozen in liquid nitrogen and stored at -80°C at a protein concentration of 43 mg/mL for further use. Mitochondria were isolated from mice liver as previously described^72^. At the end of the isolation procedure, mitochondria were flash-frozen in liquid nitrogen and stored at −80 °C at a protein concentration of 25 mg/mL until further use.

For cryo-electron tomography experiments, approximately 0.5 g of mitochondrial pellet was thawed on ice. 20 µl of mitochondria was diluted in 80 µL of ice-cold freezing buffer (250 mM trehalose, 20 mM HEPES/KOH pH 7.4). To induce pore formation, 1 μL of diluted mitochondria was added to 10 μl of 5 μM caspase-3-cleaved GSDME-N and incubated at 37 °C for 1 h. Following incubation, 6 μL of sample was applied to freshly glow-discharged R3.5/1 Au 200-mesh holey carbon grids (Quantifoil Micro Tools). Grids were plunge-frozen in liquid ethane using a Vitrobot Mark IV plunge-freezer (Thermo Fisher Scientific) operated at 4 °C and 100% relative humidity. Samples were blotted for 16–17 s using grade 597 filter paper (Whatman) with a blot force setting of −1 before vitrification.

Cryo-electron tomography data were acquired using a Titan Krios G4 transmission electron microscope (Thermo Fisher Scientific) operating at 300 kV and equipped with a SelectrisX energy filter and a Falcon 4i direct electron detector. Automated data acquisition was performed using SerialEM. Tilt series were collected from -60° to 60° using a dose-symmetric acquisition scheme with 3° increments. Images were recorded at a nominal magnification of 53,000×, corresponding to a calibrated pixel size of 2.414 Å. A total electron dose of approximately 120 e⁻/Å² was distributed evenly across 41 tilt images. Target defocus values ranged from −2.0 to −4.5 μm.

Tilt-series alignment and tomographic reconstruction were performed using AreTomo v1.3.3. Reconstructed tomograms were binned fourfold prior to further analysis. Tomograms were visually inspected and processed using IMOD. For visualization of mitochondrial membranes, tomograms were automatically segmented using MemBrain-seg. GSDME-N pores and ATP synthase complexes were manually positioned and annotated in ChimeraX based on the tomographic data. Density representations of the structures were generated using the *molmap* command on RCSB PDB:5ara for ATP synthase and RCSB PDB:9H5M for the GSDME-N pore in ChimeraX for illustrative visualization.

### Molecular dynamics simulations

We first performed a coarse-grained molecular dynamics simulation of the GSDME pore embedded in a lipid bilayer using the Martini 3 force field^73^. The cryo-EM structure of the 30-mer ring was coarse-grained using Martinize^74^, and embedded into a lipid membrane using insane^75^. The membrane composition was 20% CL and 80% DOPC. The system was solvated in water, and NaCl ions were added to a final concentration of 0.15 M to neutralize and buffer the system. The system size after equilibration was 50 × 50 × 18 nm³, containing a total of 393959 beads. The membrane bilayer consisted of 4529 DOPC lipids and 1132 cardiolipins.

Energy minimization was carried out using the steepest descent algorithm for 15000 steps. Following the protocol from^76^, the system was equilibrated through a series of NVT and NPT stages, during which restraints on lipid headgroups and the protein atoms were progressively released. NVT equilibration was performed for 1.5 ns at 310 K using a Berendsen thermostat ^77^ with a coupling time constant of 1 ps. This was followed by 60 ns of NPT equilibration at 310 K and 1 bar using a Berendsen barostat^77^ (compressibility of 3×10⁻⁴ bar⁻¹).

The production run was performed for 10 microseconds at 310 K using a velocity-rescaling thermostat (time constant: 1 ps) and a Parrinello–Rahman semi-isotropic barostat^78^ (x and y directions coupled together; time constant: 1 ps; compressibility: 3×10⁻⁴ bar⁻¹). The integration time step was 20 fs.

Three structures from the coarse-grained trajectory at 6, 8, and 10 microseconds, respectively, were selected as starting points for all-atom simulations, in order to sample different lipid configurations. Backmapping of the protein and lipid bilayer (excluding solvent) was performed using cg2at^79^. The backmapped protein was aligned to the original cryo-EM structure, and then replaced with the cryo-EM coordinates. To prevent steric clashes, lipids within 1.5 Å of any heavy protein atom were removed using a custom Python script and the MDAnalysis package^80^. The system size after equilibration was 50 × 50 × 18 nm³ for all three replicas, with a total of 4,422,364, 4,412,827, and 4,414,537 atoms for replicas 1, 2, and 3, respectively. The membrane bilayer consisted of 4315 DOPC lipids and 1091 cardiolipins in replica 1, 4364 DOPC and 1087 cardiolipins in replica 2, and 4343 DOPC and 1096 cardiolipins in replica 3.

This yielded three distinct starting configurations with different lipid arrangements. All-atom simulations were run using the CHARMM36m^81^ force field and TIP3P^82^ water model. Electrostatics were treated using Particle Mesh Ewald^83^ (PME), with a 1.2 nm cutoff for real-space electrostatic and van der Waals interactions. Each system underwent energy minimization using the steepest descent algorithm for 15000 steps. NVT equilibration was performed for 0.250 ns using the v-rescale thermostat with a time constant of 1 ps, followed by 1.625 ns of NPT equilibration using the same thermostat and a Parrinello–Rahman semi-isotropic barostat (time constant of 5 ps, compressibility = 4.5×10⁻⁵ bar⁻¹). Production runs included three replicas: one of 250 ns and two of 100 ns.

Contact analysis was performed using a custom Python script with the MDAnalysis library. For all-atom simulations, the first 50 ns were discarded for equilibration; for the coarse-grained simulation, the first 2 microseconds were discarded. Only contacts involving phosphate headgroups of the lipids were considered. Cardiolipins contain two phosphate headgroups. Since both can interact with the protein, at most one contact per CL was counted per residue (Figure 4E and Suppl. Figure 7B) per frame, even if both phosphate groups were involved. The contact cutoff distance was set to 5 Å between heavy atoms of lipid headgroup and protein. To smooth the contact profiles, we applied a Gaussian filter using the gaussian_filter1d function in Python (width=2 residues). CL enrichment at each residue i was calculated as: 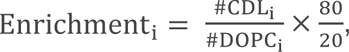, where #CDL and #DOPC represent the smoothed number of CL and DOPC contacts with residue i, respectively. The factor 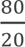 normalizes the ratio by the relative membrane composition (20% CL, 80% DOPC).

The electrostatic surface map in Figure 4D was computed using pdb2pqr^84^ for a GSDME dimer taken out of the ring structure and visualized in ChimeraX ^70^. Structural snapshots in Figure 4A-C and Suppl. Figure 7A were generated using VMD^85^. Coarse-grained simulations were performed with GROMACS^86^ version 2022.4, and all-atom simulations with GROMACS^86^ version 2024.4.

### Transmission electron microscopy (TEM)

To analyse the ultrastructure of mitochondria during apoptosis in primary WT and GSDME KO lung fibroblasts 0.2x10^6^ cells were seeded per well on top of 3 aclar foil discs in a 12-well plate. The cells were left untreated or treated with 5 µM ABT-737 and 5 µM S63845 for 2h and fixed for 1 h in 2% Glutaraldehyde with 2.5% Sucrose and 3mM CaCL2 in 0.1M HEPES buffer pH 7,4. Samples were washed three times with 0.1M Cacodylate buffer and incubated with 1% Osmiumtetroxid and 1% Potassium hexacyanoferrate for 1 h at 4°C. After 3x 5 min wash with 0.1M Cacodylate buffer, samples were dehydrated at 4°C using ascending ethanol series (50%, 70%, 90%, 3x100%) for 7 min each. Infiltration was performed with a mixture of 50% Epon/ethanol for 1h, 70% Epon/ethanol for 2h and with pure Epon overnight at 4°C. Samples were embedded into TAAB capsules and cured for 48 h at 60°C. Ultrathin sections of 70 nm were cut using an ultramicrotome (Leica Microsystems, UC6) and a diamond knife (Diatome, Biel, Switzerland) and stained with 1.5 % uranyl acetate for 15 min at 37°C and 3% Reynolds lead citrate solution for 4 min. Images were acquired using a JEM-2100 Plus Transmission Electron Microscope (JEOL) operating at 80kV equipped with a OneView 4K camera (Gatan). Individual mitochondria (n > 80) from 4-5 independent cells per condition were analyzed in a blinded way. The cristae morphology was classified into three categories: regular, altered not swollen and altered swollen. Additionally, the cristae width was quantified using Microscopy Image Browser (MIB). For n = 29-32 mitochondria, individual cristae were outlined with a B/W threshold (110), manually filled and saved as a mask. Using the Plug-in “Plasmodesmata” -> “Cell wall thickness” a central axis was automatically generated for each traced crista and the cristae width was measured at every point perpendicular to the axis. The median width per crista was obtained. For each mitochondrion, the mean of the median widths of all measured cristae was calculated to represent the average width per mitochondrion. Only well-resolved and clearly discernible cristae were included in the analysis. Statistics were determined using Ordinary one-way ANOVA corrected for multiple comparisons with Tukey test in GraphPad Prism. Galleries of representative images were created with Omero.

### Visualization of MIM extrusion and mtDNA release

Primary WT and GSDME KO lung fibroblasts 0.2x10^6^ cells were seeded per well on top of a 15 mm coverslip in a 12-well plate and stained with Mitotracker orange on the following day for 30 min. Then the cells were treated with 5 µM ABT-737 and 5 µM S63845 for 2h and fixed with 4% paraformaldehyde for 10 min. The cells were washed 3x with PBS at RT and quenched with 50 mM of NH4Cl in PBS for 15 min at RT and washed 3x. After permeabilisation with 0.25% TritonX-100 in PBS for 8 min, the cells were washed 3x with PBS and blocked for 1h with 2% BSA in PBS. In order to visualize the extrusion of the inner mitochondrial membrane out of the outer membrane during apoptosis, the cells were immunoblotted with α-TOM20 antibody and a secondary Abberior STAR green antibody. The samples were mounted and confocal images were acquired with a TCS SP8 gSTED 3X microscope (DFG-INST 2016/742-1 FUGB, Leica Microsystems) equipped with PL APO 63x/1.40 oil immersion objective and the LASX 3.5.7.23225 (Leica) software. Excitation lasers were 550 nm and 488 nm. Individual mitochondria (n = 27-150) from at least 7 cells per genotype and condition were evaluated and counted for MIM extrusion events. Statistics were measured via Welch’s *t*-test in GraphPad Prism. Mitotracker overlap with the TOM20 signal was measured using Fijis BIOP JACoP plugin to obtain the Manders coefficient. Statistics were measured via Mann-Whitney test. Circularity of Mitotracker signal (MIM) per cell was measured with ImageJ/Fiji and statistics were measured via Brown-Forsythe and Welch ANOVA test using GraphPad Prism. Galleries of representative images were created with Omero.

For the detection of mtDNA release during apoptosis, the cells were immunoblotted with α-DNA antibody and a secondary Abberior STAR red antibody. The samples were mounted and confocal images were acquired with a TCS SP8 gSTED 3X microscope (DFG-INST 2016/742-1 FUGB, Leica Microsystems) equipped with PL APO 63x/1.40 oil immersion objective and the LASX 3.5.7.23225 (Leica) software. Z-stacks were acquired. Excitation lasers were 550 nm and 647 nm. The respective Z-stacks were transformed into maximum intensity projections in Fiji/ImageJ to assess mtDNA release. We used a home-built software to detect mitochondria and create masks of them. The nuclei were cropped out and the mtDNA overlap with the mitochondria masks was measured using Fijis BIOP JACoP plugin to obtain the Manders coefficient. n = 3 independent experiments were performed. Levels of significance were determined using the Welchs’ *t*-test in GraphPad Prism. Galleries of representative images were created with Omero.

### Generation of non-cleavable *Gsdme^D270A^* mice

To generate *Gsdme^-/-^* mice and *Gsdme^DA/DA^* mice, a single guide RNA (Gsdme gRNA: 5′-TTTCTGGACATGCTGGATGG -3′) targetting exon 7 was in vitro-transcribed and co-electroporated into fertilized wild-type oocytes together with Cas9 protein (IDT) and a ssDNA HR template carrying the desired point mutations (5’TAGACTCTGTCTACTTGGACCCCCTGGCCTACAGAGAGTTCGCCTTTCTGGA CATGCTGGccGGaGGTCAAGGGATCTCTTCGCAGGACGGGCCCCTACGAGTTGTAAAACAAGGTATTGCCTGT3’). After confirmation of the correct mutations by genomic DNA sequencing analysis, founder mice carrying the targeted mutations were backcrossed to C57BL/6N mice to establish independent mouse lines. Mice were maintained at the specific-pathogen-free (SPF) animal facility of the CECAD Research Center of the University of Cologne.

### IncuCyte experiments

A total of 0.01x106 primary WT and GSDME KO lung fibroblasts were seeded per well of a 96-well plate. The cells were stained with TMRE (150 nM) to observe mitochondria membrane potential changes. Cytotox green (1:3000) was used to detect cell death kinetics, MitoSOX red (5 µM) to detect ROS formation and Caspase 3/7 detection reagent (1:500) to detect Caspase 3/7 activation in the cells. Apoptosis was induced with 5 µM ABT-737 and 5 µM S63845 and the respective fluorescence intensity was measured as high red or mean total fluorescence per well by acquiring 4 images per well every 30 min for 24 h using the IncuCyte S3 (Sartorius) at 37°C with 5% CO_2_. Data were normalized to the 0-time point.

To determine the killing activity of GSDME-N-GFP mutants in mammalian cells, the cellular uptake of DRAQ7 was monitored as an indication of plasma membrane integrity. HeLa WT cells were first transfected at 80% confluence with plasmids encoding GFP, WT GSDME-N-GFP or the indicated GSDME-N-GFP mutants using jetOPTIMUS. Subsequently, the medium was replaced with medium containing DRAQ7 (1:3000). To assess mitochondrial membrane potential loss, cells transfected with GFP, WT GSDME-N-GFP or the indicated GSDME-N-GFP mutants were incubated with TMRE (150 nM). Four to nine images per well were acquired every 1 h for 24 h. For DRAQ7 analysis, cells were classified as DRAQ7-positive or -negative as a measure of plasma membrane permeabilization. For TMRE analysis, red fluorescence intensity was quantified per cell, and cells were classified as either retaining or having lost mitochondrial membrane potential using the same cell-by-cell analysis parameters across all constructs and conditions. At least three independent experiments were performed. Images were analysed using the IncuCyte cell-by-cell analysis software module and OriginPro.

To determine the activation of the cGAS/STING pathway, we used a STING-Biosensor stably expressed in HeLa cells^48^. For GSDME knockdown, cells were either transfected with 40 nM of DFNA5-targeting or a non-targeting control siRNA using Lipofectamin2000 for 48 h. Afterwards, apoptosis was induced with 1 µM ABT-737 and 1 µM S63845, and co-treatment with 10 µM of the pan-caspase inhibitor QVD-OPH (QvD) was used to inhibit caspase-dependent deactivation of cGAS. Treatment with diABZi was used as a positive control for STING activation. HeLa STING-Biosensor cells were treated with DDC (2’,3’-dideoxycytidine) for 6 days to deplete mtDNA and create Rho^0^ cells as a control to show that the resulting STING activation is triggered only by mtDNA. To determine kinetics allowing GSDME activation but also STING activation, first QVD was titrated as indicated with a stable A+S concentration and afterwards A+S were titrated as indicated with 15 µM of QVD in the HeLa STING Biosensor cell line. The cells were stained with TMRE (50 nM) to observe mitochondrial membrane potential changes, Annexin V to measure PS exposure (1:500), and DRAQ7 (1:3000) was used to detect cell death kinetics. Four images per well were acquired every 1 h for 24 h. At least n = 3 independent experiments were performed. The images were analysed using the IncuCyte basic analysis software module and Prism. Data were normalized to the 0-time point. Levels of significance were determined using the Wilcoxon test in GraphPad Prism.

### qPCR

To determine the RNA levels after the CRLS1 knockdown, a part of the total lysate was used for qPCR. RNA was isolated using the NEB Monarch RNA isolation kit (NEB) as per manufacturer instructions. RT-qPCR was performed directly from the RNA using Luna Universal One-Step RT-qPCR Kit (New England Biolabs) as per manufacturer instructions except that 10μL reactions were performed in 384 well plates. Cycling conditions for Applied Biosystems was used and were: 55C for 10 min followed by 95C for 1 min, followed by 40 cycles fo 10s at 95C and 1 min at 60C, followed by the melt curve stage. Fold change was calculated as 2-ΔΔCt. Small cell lung cancer cells (SCLCs) of all genotypes were seeded at a density of 4 x 10^5^ cells per well in a 6-well plate and incubated at 37°C for 2 h to let cells attach, which was confirmed under the microscope before drug treatment. Cells were treated with 10 µM ABT-737 and S63845 (A+S) and 20 µM QVD-OPH (Q) for 24h. Primary lung fibroblasts of all genotypes were seeded at a density of 4 x 10^5^ cells per well in a 6-well plate and treated on the next day with 5 µM A+S and 20 µM Q. After treatment, cells were washed twice with DPBS, trypsinized and collected in RNase-free tubes with centrifugation at 300 x g for 4 min. RNA extraction was performed following the manufacturer’s instructions (QIAwave RNA Plus Mini Kit, QIAGEN, cat no: 74636). RNA concentration was measured with the NanoPhotometer. 200 ng RNA were synthesized to cDNA with the LunaScript® RT SuperMix (New England Biolabs, cat no: M3010X). RT-qPCR was performed using the Luna^®^ Universal qPCR Master Mix (New England Biolabs, cat no: M3003E) with 20 ng cDNA per reaction. Relative gene expression (RE) was calculated using ΔΔCt method.

### Data availability statement

All data necessary to evaluate our findings are present in the paper or the Supplementary Information. The 3D cryo-EM density map of GSDME-N have been deposited to the Electron Microscopy Data Bank under the accession number EMD-51889. The atomic coordinates have been deposited to the Protein Data Bank under the accession code 9H5M. All other data supporting the findings of this study are available from the corresponding author upon reasonable request.

## Acknowledgements

We thank W. Kühlbrandt for providing the *Polytomella* mitochondria and for sharing his expertise in single particle cryoEM, M. Strater and L. Faber for technical support and A. Villunger for critically reading the manuscript. We thank the CECAD Imaging Facility and A. Schauss, K. Seidel and C. Jüngst for their support in EM and gSTED microscopy. We also thank B. Zevnik and the CECAD Transgenic Core Facility for CRISPR/Cas9-assisted gene targeting in mice. This work has been funded by the European Research Council (Grants Nr. APOSITE -817758 and MITOPORE -101195637) and by the Deutsche Forschungsgemeinschaft (CRC1218, A09 to A.J.G.S; CRC1403, A02 and B07 to A.J.G.S. and A10 to N.P.; CRC1507, P21 to A.J.G.S. and P12 to G.H.; CRC1399-413326622 and CRC1530-455784452 to N.P; CRC1557, P5-467522186SFB to K.C.; and EXC-3094 – 533751785 to A.J.G.S. and G.H.), as well as by the University of Cologne and the Max Planck Society. S.A. was supported by Nachlass Johann und Therese Röhig Project ID-No. 43-2024 of the Medical Faculty, Univ. Cologne.

