## Supplemental data for "Structural mechanism of Gasdermin E-mediated mtDNA release from apoptotic mitochondria"

**Supplementary information**

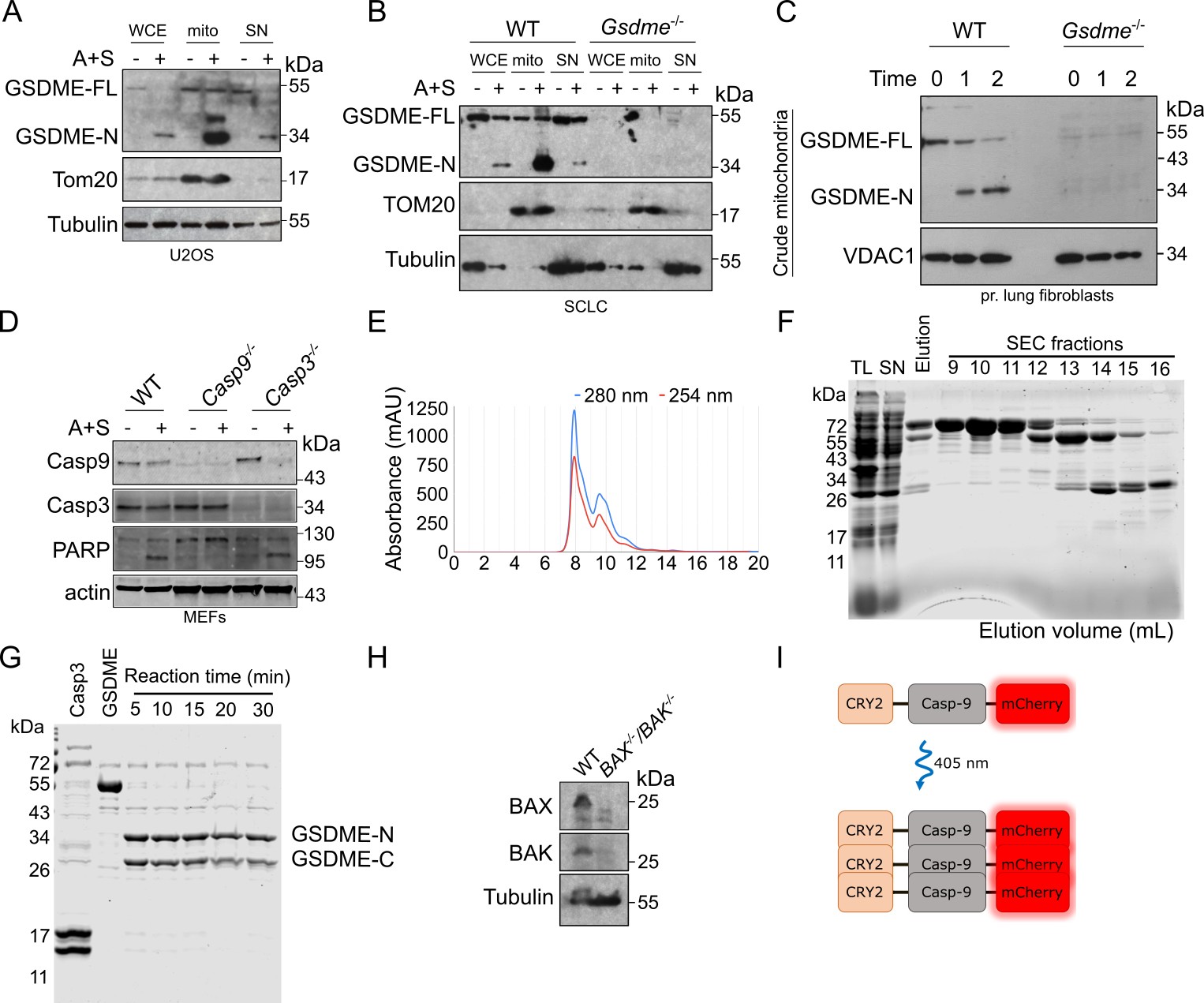

**Suppl. Figure 1: GSDME mitochondrial targeting, purification and scheme of Opto-GSDME-N construct**

A-C) Representative Immunoblots of GSDME cleavage and accumulation at mitochondria of U2OS cells (A), WT and *Gsdme^-/-^* mouse SCLC cells (B), and WT and *Gsdme^-/-^* primary mouse lung fibroblasts (C) treated or not with ABT737 and S63845 (A+S) for 2 h. Whole cell extract (WCE), crude mitochondria (mito), and supernatant (SN) fractions are shown. U2OS cells were treated with 1 µM A+S, SCLCs with 10 µM A+S, and primary lung fibroblasts with 5 µM A+S.

D) Representative immunoblot confirming indicated caspase knockouts in healthy and apoptotic MEF cells treated with 5 µM A+S for 2 h.

E) Size-exclusion (SEC) Chromatogram from a Superdex75 column after injection of the elution fraction of purified recombinant GSDME.

F) Representative Coomassie-stained SDS-PAGE gel for the steps of the purification process of recombinant GSDME. TL: total lysate, SN: supernatant after lysis and centrifugation, SEC: size-exclusion chromatography fractions from 9 to 16 ml. Fraction-13 was used for further experiments.

G) Representative Coomassie-stained SDS-PAGE gel for the cleavage of recombinant GSDME by caspase 3.

H) Representative immunoblot confirming BAX and BAK double knockout in U2OS cells.

I) Representative schematic illustration of the CRY2-Caspase-9-mCherry construct. UV light-induced CRY2 oligomerization leads to the activation of Caspase-9-mCherry, which subsequently activates apoptotic executioner caspases.

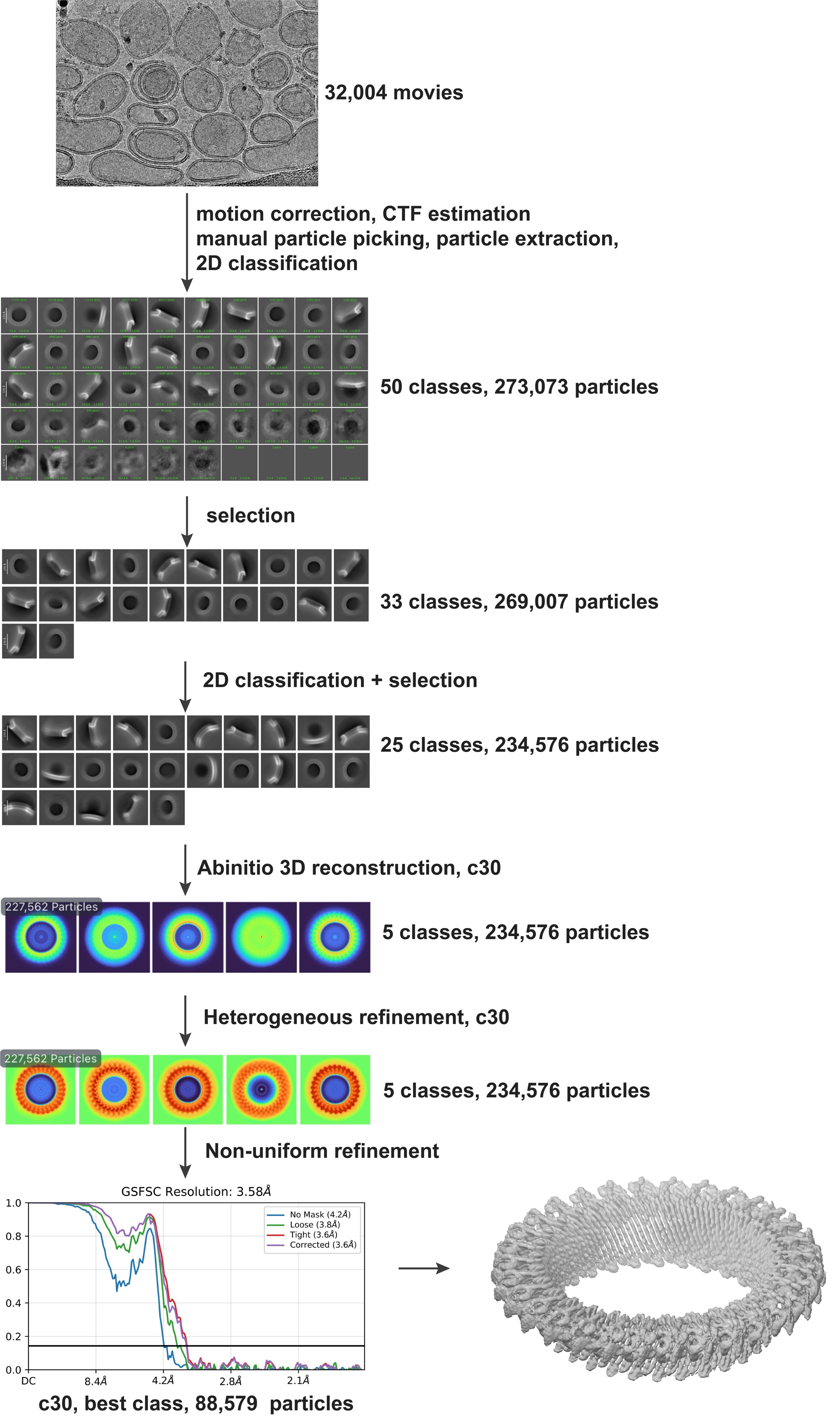

**Suppl. Figure 2: Cryo-EM processing of GSDME-N.**

Flowchart of cryo-EM image processing. Representative cryo-EM micrograph, 2D class averages, and 3D reconstructions of GSDME-N on liposomes. The final 3D refinement (non-uniform refinement) is performed with 88,579 particles. The FSC curves for the final map with applied c30 symmetry indicate a resolution of 3.58 Å.

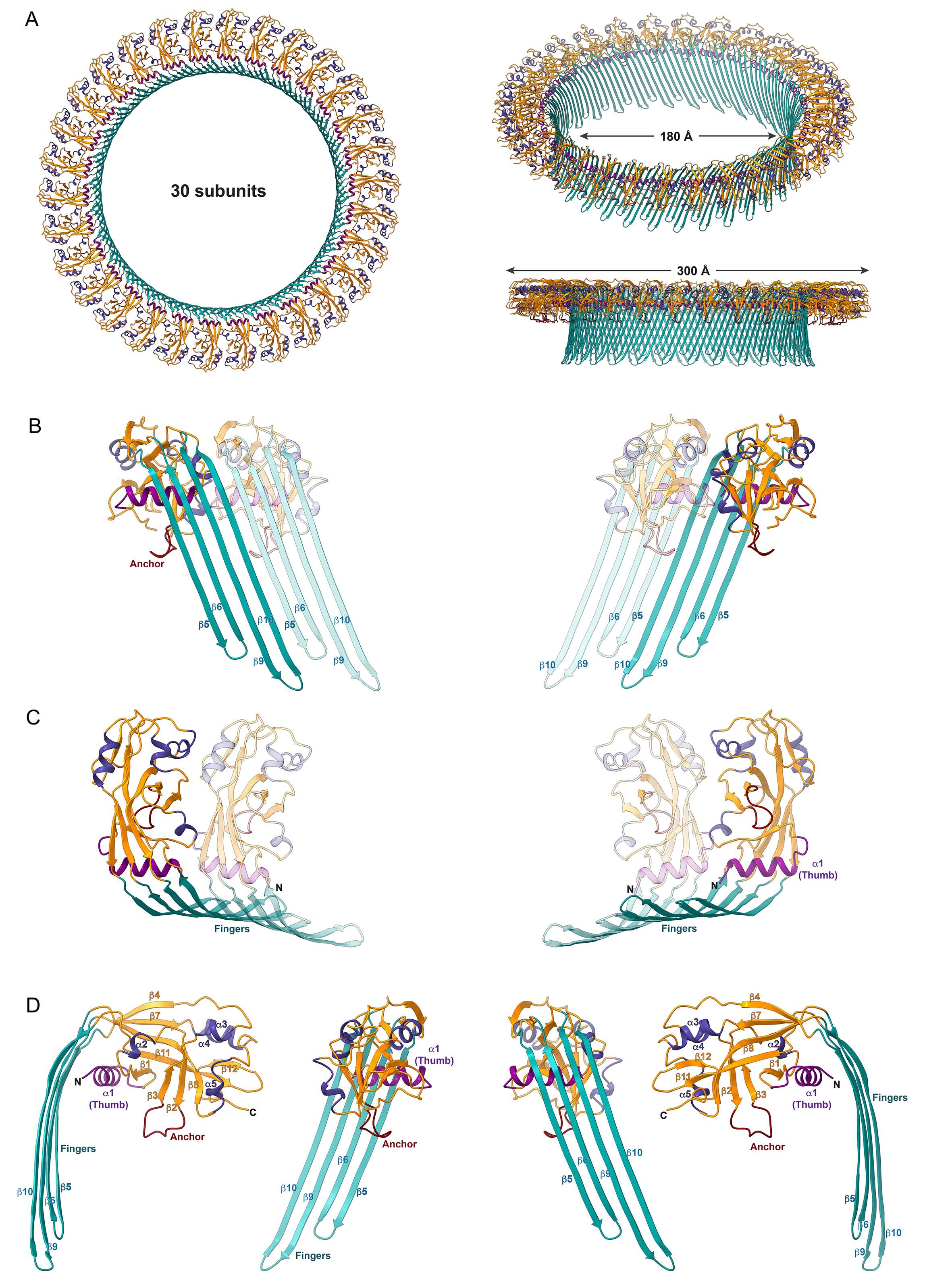

**Suppl. Figure 3: Structural details of the GSDME-N pore complex.**

A) The overall structure of the GSDME-N pore complex with 30 subunits is shown from different perspectives with details of the inner and outer diameter.

B) Two neighboring subunits shown from inside (left) and outside (right) the pore. The main interactions for the oligomerization of the subunits are those between β-strand β5 and β10.

C) Interactions between two subunits in the globular domain of GSDME-N. The thumb helix is positively charged at its N-terminus and negative on the other side (Suppl. Figure 5F,I) and forms a bipolar interface for oligomerization.

D) The structural elements for one subunit of GSDME-N in the pore state are shown from different perspectives.

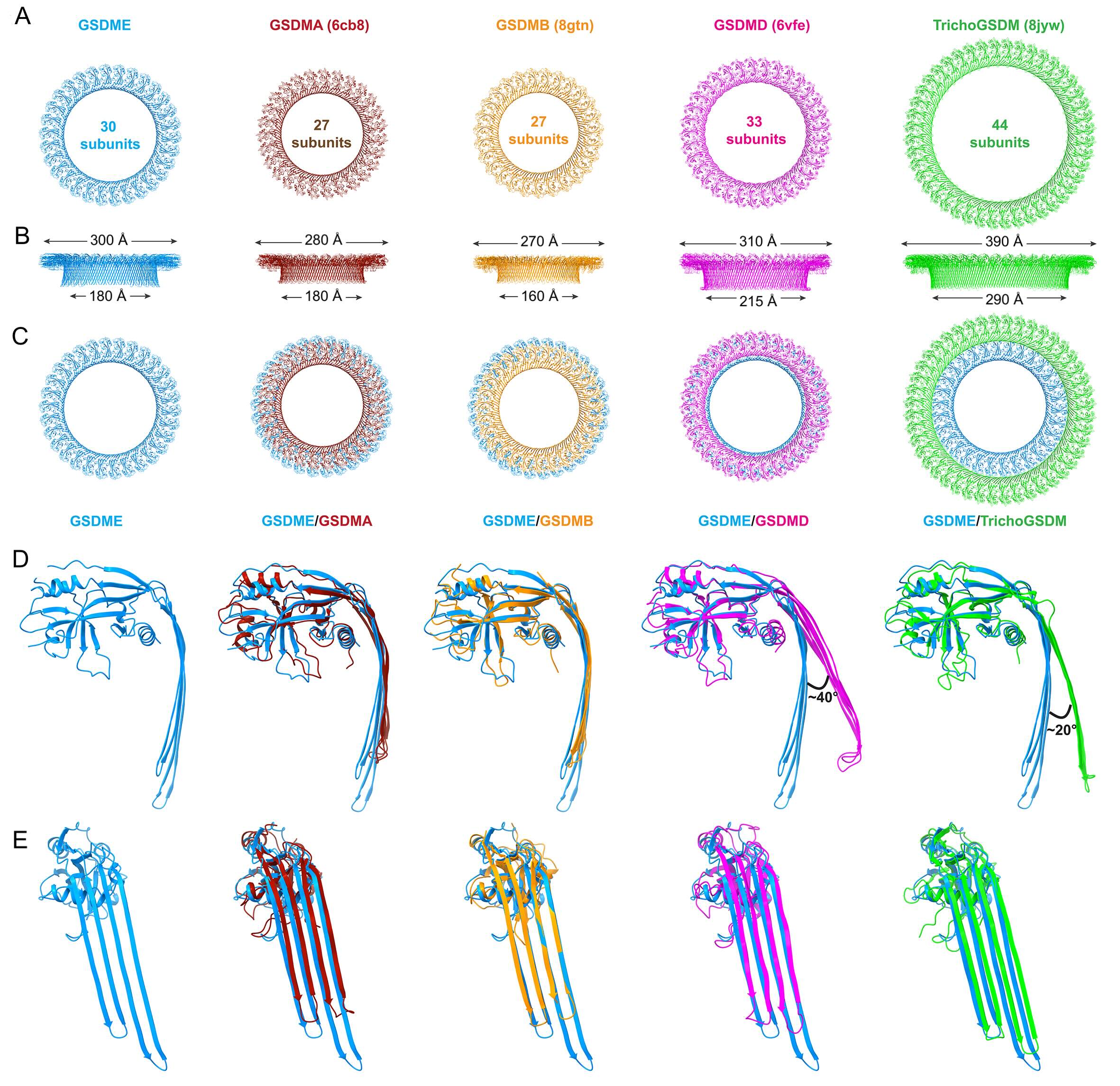

**Suppl. Figure 4: Comparisons of GSDME with structurally known GSDMs.**

GSDME (blue) is compared with GSDMA (brown), GSDMB (orange), GSDMD (magenta), and *Tricho*GSDM (green). In the first and second rows, the individual pore complexes are shown from above and along the membrane plane, respectively. The top-view shows the number of subunits in the pore complex in the ring. The side-view (second row) shows the size of the outer diameter above, and that of the inner diameter is shown below the molecule. The superposition of GSDME-N with the other GSDMs is shown in the third row. In the fourth row, the superpositions of the individual subunits are shown from the side. The curvature of the GSDMA and GSDMB fingers is very similar to that of GSDME, although the fingers of GSDME are considerably longer. The fingers of GSDMD and *Tricho*GSDM are similar in length to those of GSDME, but they are less curved. As a result, the angle in relation to the globular domain is greater in these two pore complexes, which have more subunits. The fifth row shows the superposition of the subunits seen from the pore center.

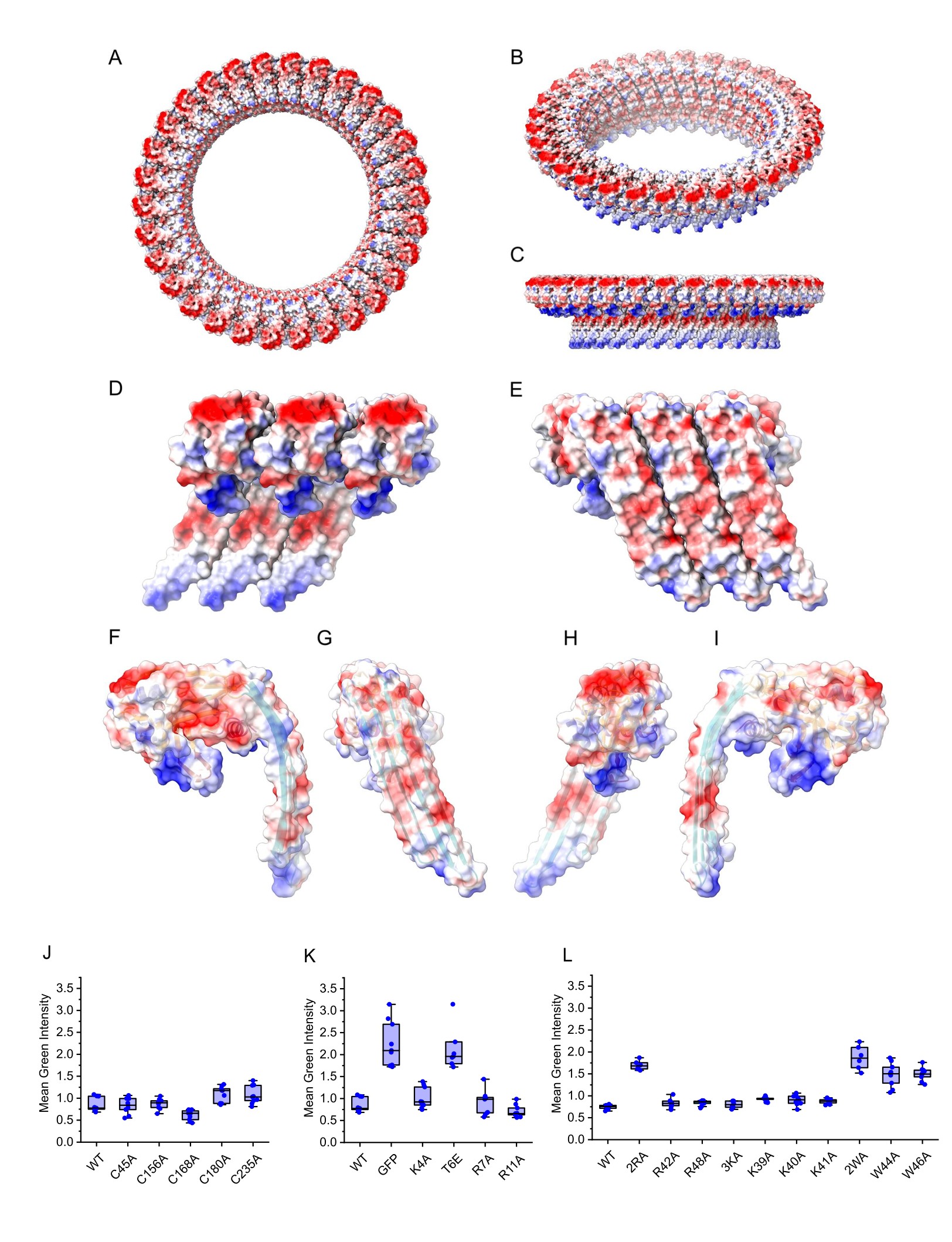

**Suppl. Figure 5: Charge distribution on the GSDME-N pore complex.**

A-I) Surface potential presentation for the entire GSDME-N pore complex (A-C), three subunits side-by-side (D, E), and for one subunit (F-I). Overall, positive charges (blue) dominate on the membrane-facing side of the globular domain (palm), while the opposite, solvent-exposed side is mainly negatively (red) charged (A-C). The fingers (D, E) are slightly negative within the pore. A hydrophobic patch on the membrane-facing side of the finger is flanked by positive charges on the tip of the fingers and negative charges on the other side. Contact surfaces of adjacent GSDME-N monomers in the pore complex show charge complementarity. Here, the thumb α-helix, which forms a continuous ring around the β-barrel, is negatively charged on one side (F) and positively charged on the other. The positively charged region of the hydrophobic tip of the anchor stands out in particular.

J-L) Green fluorescence intensity of HeLa cells transiently transfected with GSDME-N-GFP and indicated mutants or with GFP at 14 h. Data show similar expression levels amongst most of the constructs compared to the WT. Mutants displaying reduced cytotoxicity showed higher fluorescence intensities than WT. The center line of the box represents the median, while the box limits represent 25th and 75th percentiles.

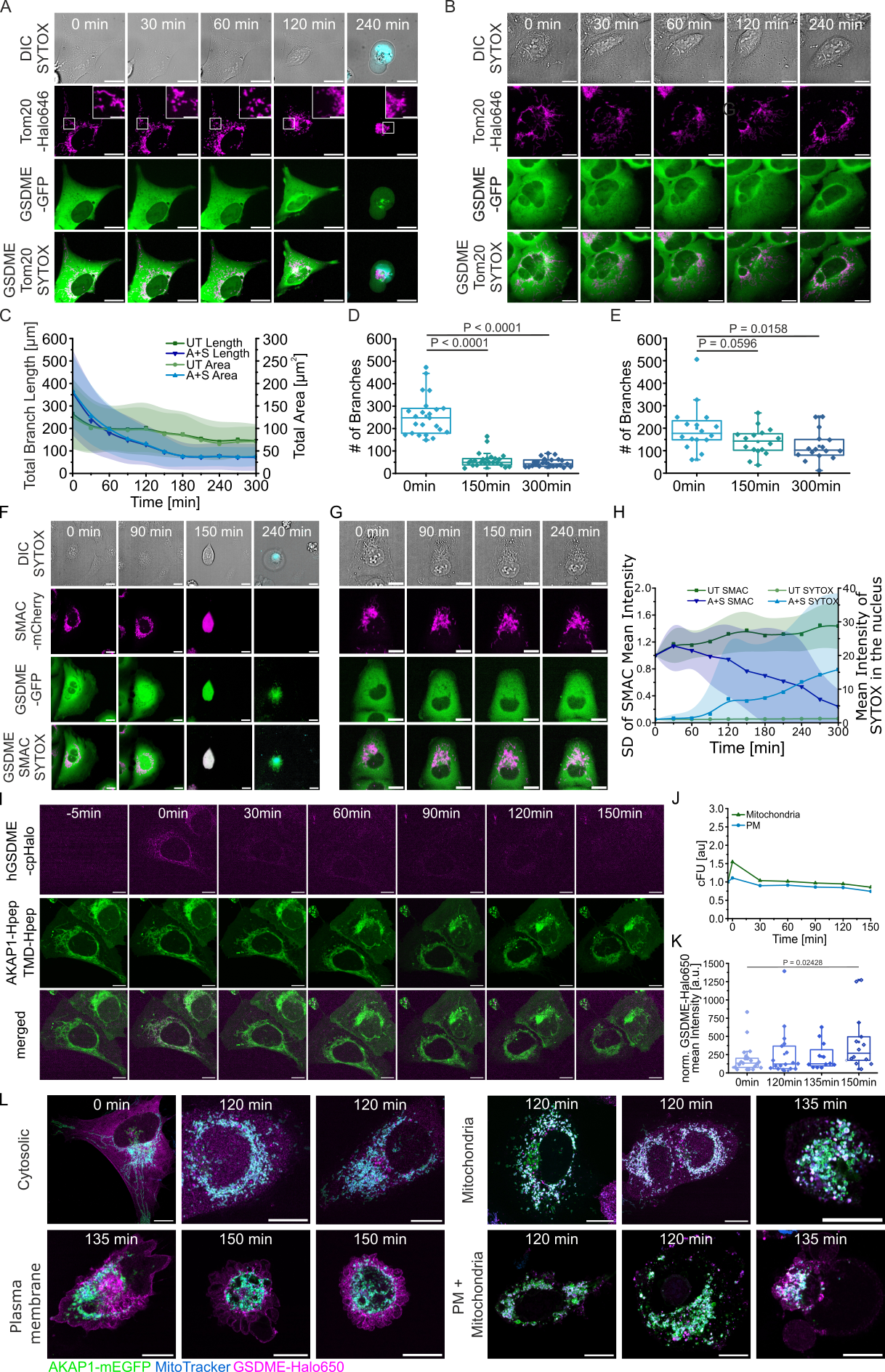

**Suppl. Figure 6: Super-resolution microscopy of GSDME in mitochondria in fixed and live cells.**

A-B) Confocal time-lapse images of apoptotic (A) and untreated B) U2OS WT cells expressing hGSDME-mEGFP (green) and Tom20-HaloTag (magenta). Plasma membrane permeabilization was monitored by SYTOX Blue uptake (cyan) and morphological changes by brightfield microscopy. Zoomed images show mitochondrial fragmentation characteristic of MOMP. Scale bar: 20 µm, for zoomed images 5 µm (apoptotic) and 10 µm (untreated).

C-E) Quantitative analysis of mitochondrial fragmentation in A and B, calculated as Total Branch Length (C), Total Area of mitochondria (D) and Number of Branches of mitochondria € per cell. Shown is mean ± SD. (C,D). Data are presented as box plots with the center lines at the median, lower bound at 25th percentile, upper bound at 75th percentile. (n = 3 with 16 cells for untreated and n = 3 with 23 cells for treated).

F-G) Confocal time-lapse images of apoptotic (F) and untreated (G) U2OS WT cells expressing hGSDME-mEGFP (green) and SMAC-mCherry (magenta). Plasma membrane permeabilization monitored by SYTOX Blue uptake (cyan) and morphological changes by bright field microscopy. Scale bar: 20 µm.

H) Quantitative analysis of SMAC release and SYTOX uptake in treated and untreated U2OS WT cells in (F-G). Data normalized to time point 0 min of either SMAC-mCherry or SYTOX Blue. Shown is mean ± SD. n =3 with 45 cells for untreated and treated.

I) Representative Airyscan time-lapse images of untreated U2OS WT cells expressing hGSDME-cpHalo (magenta), AKAP1-mEGFP-Hpep3 and puro-leader-aALFAnb-TMD-mEGFP-Hpep3 (green). The -5 min images are before HTLJFx650 addition and before apoptosis induction. Cells were then labeled with 50nM HTLJFx650 and apoptosis was induced with 1 µM ABT-737 and 1 µM S63845. Scale bar: 10 µm.

J) Quantitative analysis of hGSDME fluorescence intensity (corrected fluorescence intensity units, cFU) at mitochondria *vs*. PM over time in untreated U2OS WT cells shown in (I). Data normalized to -5 min time point. Data are representative of two independent experiments (n = 2 with 8 cells).

K) Quantification of the mean intensity of GSDME-Halo650 in the whole cell of Airyscan microscopy images (Figure 3D). Data are presented as box plots with the center lines at the median, lower bound at 25th percentile, upper bound at 75th percentile. Shown is mean ± SD. n = 3 with 20 cells for 0 min, n = 3 with 16 cells for 120 min, n = 3 with 19 cells for 135 min, and n = 2 with 24 cells for 150 min.

L) Representative Airyscan microscopy images of GSDME distribution in U2OS WT cells expressing AKAP1-mEGFP (green) and hGSDME-HaloTag (magenta) at 0 min, 120 min, 135 min and 150 min after apoptosis induction with 1 µM ABT-737 and 1 µM S63845. Matrix/MIM stained with MitoTracker Red (blue). Scale bar: 10 µm.

All statistics were measured by a two-tailed unpaired t-test with p>0.05 = non-significant. All data were tested for normality.

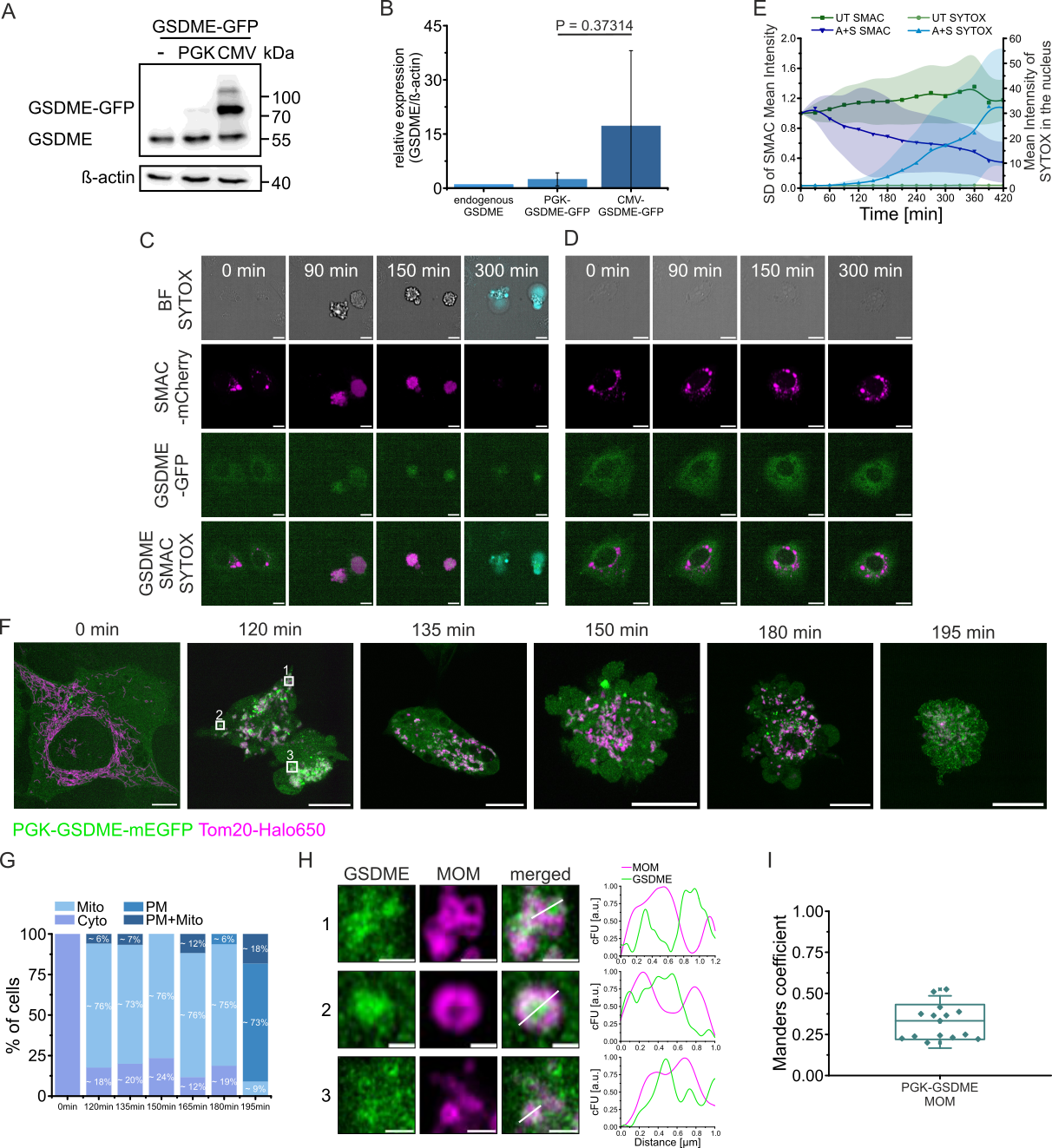

**Suppl. Figure 7: Super-resolution microscopy of PGK-GSDME-GFP in mitochondria.**

A) Representative immunoblot of whole cell extracts showing expression of endogenous (-) GSDME, PGK-hGSDME-mEGFP (PGK) and CMV-hGSDME-mEGFP (CMV) in U2OS WT cells. Representative of three independent experiments.

B) Quantification of GSDME expression relative to ß-actin. Data normalized to endogenous GSDME. Shown is the mean ± SD. n = 3.

C-D) Confocal time-lapse images of apoptotic (C) and untreated (D) U2OS WT cells expressing PGK-hGSDME-mEGFP (green) and SMAC-mCherry (magenta). Plasma membrane permeabilization was monitored by SYTOX Blue uptake (cyan) and morphological changes by bright field microscopy. Scale bar 10 µm.

E) Quantitative analysis of SMAC release and SYTOX uptake in treated and untreated U2OS WT cells in (C-D). Data normalized to 0 min time point for both SMAC-mCherry and SYTOX Blue. Shown is mean ± SD (n =3 with 45 cells for untreated and treated).

F) Airyscan microscopy images of U2OS WT cells expressing PGK-hGSDME-mEGFP (green) and Tom20-HaloTag (magenta) at different time points after apoptosis induction with 1 µM ABT-737 and 1 µM S63845. White boxes in the 120 min timepoint image correspond to zoomed regions in H. Scale bar 10 µm.

G) Quantification of the cellular distribution of hGSDME at varying time points after apoptosis induction, classified as cytosolic (Cyto), mitochondrial (Mito), at the plasma membrane (PM) or both (PM + Mito). n = 3 with >15 cells for each timepoint.

H) Airyscan microscopy images of zoomed regions in F at 120 min after apoptosis induction. The line profiles of cross-sections show colocalization of GSDME with MOM in corrected fluorescence intensity (cFU). Scale bar 1 µm for images 1 and 3 and 0.5 µm for image 2.

I) Co-localization analysis of GSDME with MOM at 120 min after apoptosis induction. Data are presented as box plots with the center lines at the median, lower bound at 25th percentile, upper bound at 75th percentile (n = 3 with 16 cells).

All statistics were measured by a two-tailed unpaired t-test with p>0.05 = non-significant. All data were tested for normality.

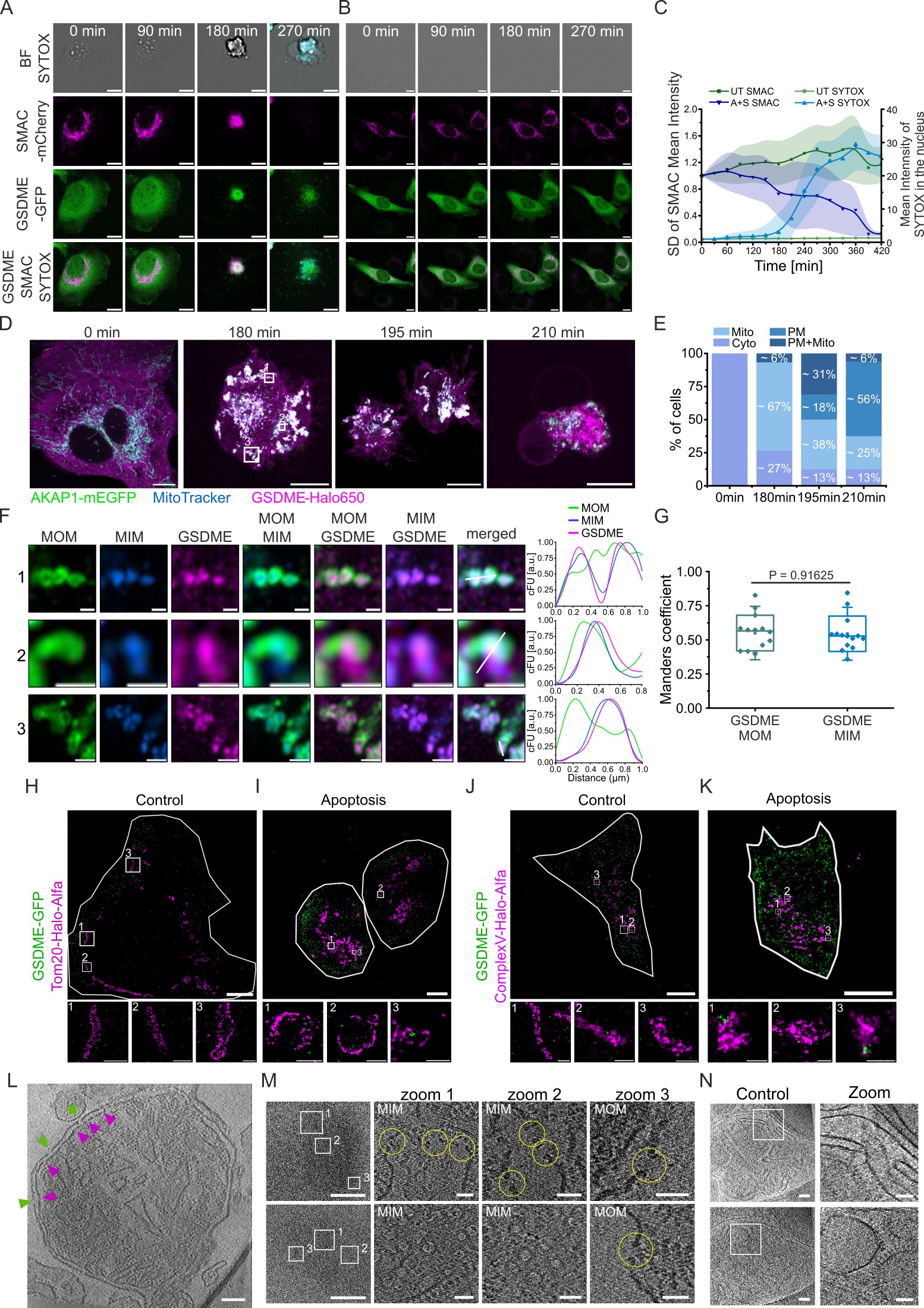

**Suppl. Figure 8: Super-resolution microscopy of HeLa cells and cryoET of GSDME in mitochondria.**

A-B) Confocal time-lapse images of apoptotic (A) and untreated (B) HeLa WT cells expressing hGSDME-mEGFP (green) and SMAC-mCherry (magenta). Plasma membrane permeabilization monitored by SYTOX Blue uptake (cyan) and morphological changes by BF. Scale bar: 10 µm.

C) Quantitative analysis of SMAC release and SYTOX uptake in treated and untreated HeLa WT cells in (A-B). Data normalized to time point 0 min of either SMAC-mCherry or SYTOX Blue. Shown is mean ± SD (n =3 with 45 cells for untreated and treated).

D) Airyscan microscopy images of Hela WT cells expressing AKAP1-mEGFP (green) and hGSDME-HaloTag (magenta) at 0 min, 180 min,195 min and 210 min after apoptosis induction with 1 µM ABT-737 and 1µM S63845. Matrix/MIM stained with Mitotracker Red (blue). White boxes in the 180 min time point image correspond to zoomed regions in F. Scale bar: 10 µm.

E) Quantification of the cellular distribution of hGSDME at varying time points after apoptosis induction, classified as cytosolic (Cyto), mitochondrial (Mito), at the plasma membrane (PM), or both (PM + Mito). n = 3 with 15 cells.

F) Airyscan microscopy images of zoomed regions in D at 180 min after apoptosis induction. The line profiles of cross-sections show colocalization of GSDME with MIM in corrected fluorescence intensity (cFU). Scale bar: 0.5 µm.

G) Co-localization analysis of GSDME with MOM and MIM at 180 min after apoptosis induction. Data are presented as box plots with the center lines at the median, lower bound at the 25th percentile, upper bound at the 75th percentile (n = 3 with 15 cells).

H-I) Representative images of post-processed two-color DNA-PAINT microscopy experiments of untreated (n = 2 experiments with 4 cells) (H) or apoptotic (n = 3 experiments with 6 cells) (I) U2OS WT cells expressing hGSDME-GFP and Tom20-HaloTag-AlfaTag. White boxes in the main image (left) correspond to zoomed regions (right). Scale bar: 5 µm, zoomed 1 = 1 µm, zoomed 2-3 = 500 nm (H) and 10 µm, zoomed 1-2 = 500 nm, zoomed 3 = 200 nm (I).

J-K) Representative images of post-processed two-color DNA-PAINT microscopy experiments of untreated (n = 2 experiments with 6 cells) (J) or apoptotic (n = 2 experiments with 4 cells) (K) U2OS WT cells expressing hGSDME-GFP and ComplexV-HaloTag-AlfaTag. White boxes in the main image (left) correspond to zoomed regions (right). Scale bar: 10 µm, zoomed 1-3 = 100 nm, zoomed 2 = 200 nm (J) and 10 µm, zoomed = 500 nm (K).

L) Full-size image of the central slice (shown in Figure 3L) through a tomographic volume of mitochondria isolated from mouse liver and incubated with recombinant hGSDME-N. The MIM can be distinguished from the MOM by the presence of the ATP synthase. Scale bar: 100 µm.

M) Representative images of central slices through a tomographic volume of mitochondria isolated from *Polytomella* and incubated with recombinant hGSDME-N. The MIM can be distinguished by the presence of the ATP synthase. The zoomed images show GSDME-N pores in the MIM (1-2) from side (up) and top (bottom) views and in the MOM (3) from side view. Scale bar: 400 nm; zoom-in: 40 nm.

N) Representative images of tomogram slices of a control (untreated) mitochondrion. The mitochondrial outer membrane (MOM) and inner membrane (MIM) can be distinguished by the presence of the ATP synthase in the MIM. Scale bar: 100 nm; zoom-in: 50 nm.

All statistics were measured by a two-tailed unpaired t-test with p>0.05 = non-significant. All data were tested for normality.

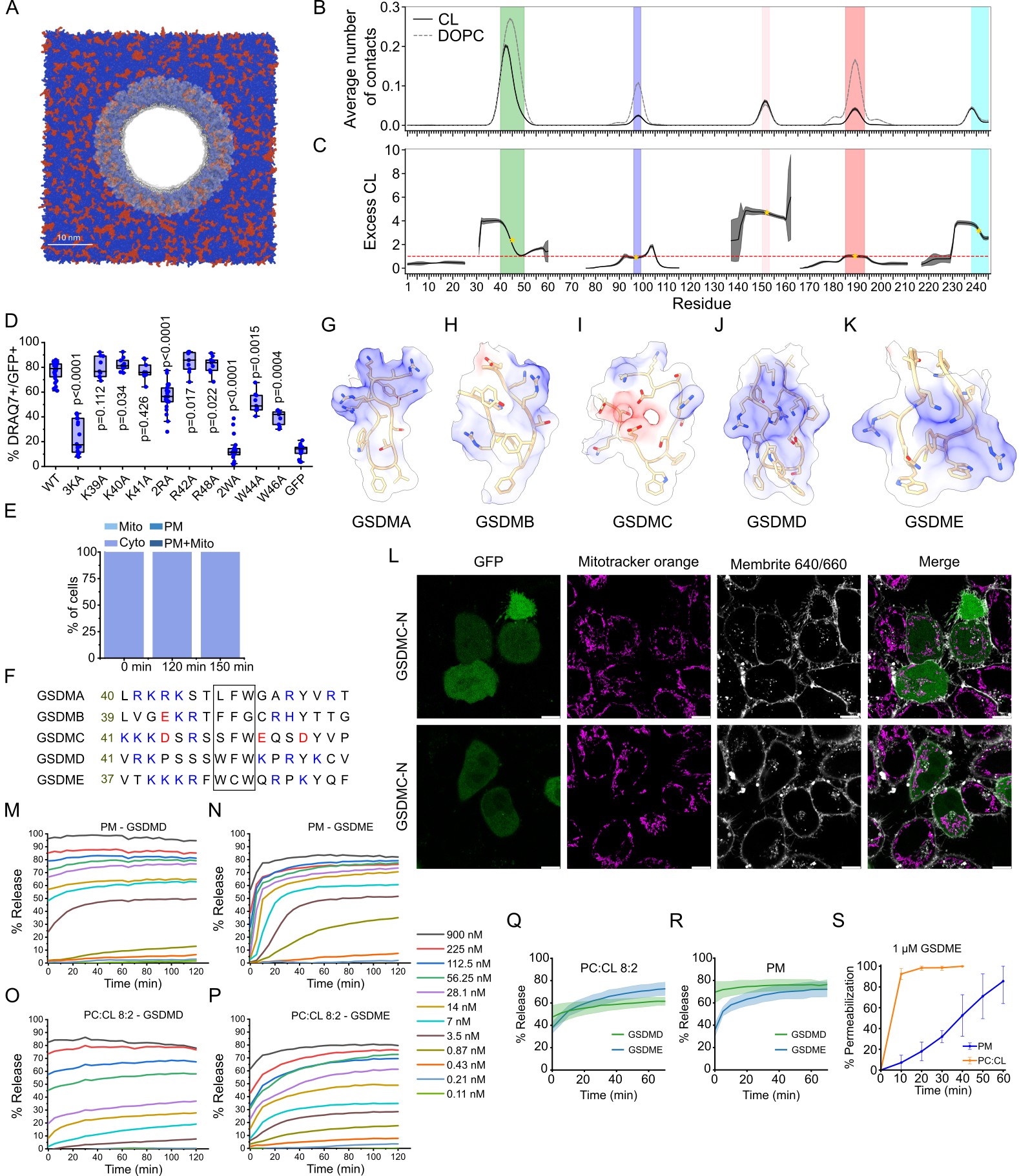

**Suppl. Figure 9: Molecular determinants of GSDME-N interactions with mitochondria.**

A) Representative top-view snapshot from an all-atom molecular dynamics trajectory showing the GSDME-N pore embedded in the membrane. The pore is shown in transparent white, cardiolipin (CL) molecules in red, and di-oleoyl phosphatidylcholine (DOPC) lipids in blue. Water and ions are omitted for clarity.

B-C) Per-residue contact frequency of CL molecules in a coarse-grained simulation of the GSDME-N pore. B) Number of cardiolipins in contact with each residue, averaged over a 10-microsecond Martini simulation, excluding the first 200 nanoseconds for equilibration (dark gray; only lipid headgroup contacts are considered). The shaded area around the curve represents the standard deviation over the trajectory. Colored shaded regions in the background correspond to the binding regions identified in Fig. 4B. The same analysis for DOPC lipids is shown in grey for comparison. C) Relative enrichment of CL molecules per residue, computed as the smoothed CL:DOPC contact ratio multiplied by 4 to normalize for lipid composition (20% CL, 80% DOPC). The red dashed line indicates no enrichment (reflecting membrane composition). Colored background regions match those in A, and golden stars indicate average enrichment within each region. Residues 40–50 (green), 150–153 (pink), and 238–245 (light blue) show higher CL contact ratios than expected from lipid abundance, suggesting preferential binding at these sites.

D) Functional analysis of anchor domain GSDME-N-GFP mutants, overexpressed at similar levels. % DRAQ7-positive HeLa cells from cells transiently transfected with GSDME-N-GFP and indicated mutants or with GFP at 24 h. 3KA (K39A, K40A, K41A), 2RA (R42A, R48A), 2WA (W44A, W46A). The center line of the box represents the median, while the box limits represent 25th and 75th percentiles. Data were analyzed with Mann-Whitney U test, with p-values compared to the WT sample.

E) Quantification of the cellular distribution of hGSDME-2WA at 120 min and 150 min after apoptosis induction, classified as mitochondrial (Mito), at the plasma membrane (PM), or both (n = 3 with>18 cells for each timepoint).

F) Sequence alignment of the anchor regions of the GSDMs shown in panels G–K. Positively charged residues are shown in blue, while negatively charged residues are shown in red. The black box highlights the hydrophobic tip within the anchor region, where aromatic and hydrophobic residues are enriched.

G-K) Ribbon-stick representations of the anchor regions from five human Gasdermin proteins: GSDMA (G, PDB:6CB8), GSDMB (H, PDB:8ET2), GSDMC (I, AlphaFold-predicted model), GSDMD (J, PDB: 6VFE), and GSDME (K, this study). Side chains of residues within each anchor region are depicted (colored according to standard molecular display: O red, N blue, and S yellow). Surfaces are shown with electrostatic potential surface coloring to visualize the distribution of charge on the anchor surface (red represents negative electrostatic potential, blue represents positive potential, and white represents neutral potential).

L) Confocal microscopy images of HeLa cells transfected with GSDMC-N-GFP (green). MitoTracker orange and Membrite640/660 were used to mark the mitochondria and plasma membrane, respectively. Scale bar: 10 µm.

M–P) Representative kinetics of carboxyfluorescein release from LUVs of different compositions induced by GSDMD-N or GSDME-N at varying concentrations as indicated. M, N) Carboxyfluorescein release over time from plasma membrane (PM)-mimicking liposomes upon incubation with GSDMD-N (M) or GSDME-N (N). O, P) Carboxyfluorescein release from phosphatidylcholine (PC):CL (8:2) liposomes in the presence of GSDMD-N (O) or GSDME-N (P).

Q-R) Kinetics of GSDME-N and GSDMD-N-dependent carboxyfluorescein release from LUVs made of PC:CL (8:2) (Q) or PM (PC:PE:SM:Chol:PS:PI:PIP2, 18:26:10:29:10:5:2) (R) at a protein concentration of 112 nM. Solid line corresponds to the mean, with ± SD shown as a shaded area (n=3 independent experiments).

S) Time-dependent permeabilization of co-incubated GUVs composed of either PC:CL (8:2) or PM=PC:PE:Chol:PS (5:3:1:1) after addition of 0.3 µM GSDME-N (n=3 independent experiments).

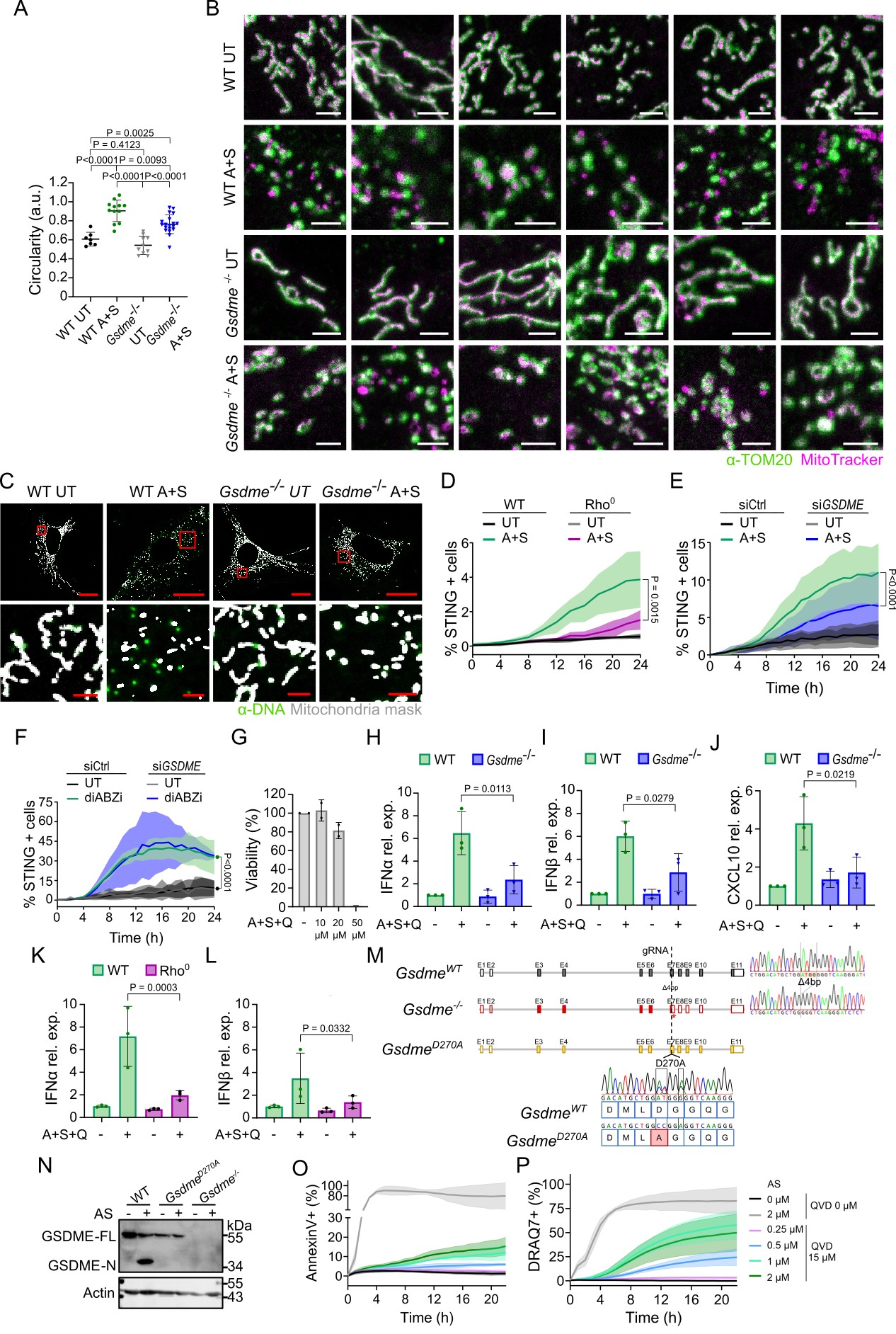

**Suppl. Figure 10: GSDME depletion affects MIM remodeling and mtDNA release in apoptosis as well as STING activation under caspase inhibition.**

A) Quantification of MIM/matrix circularity (MitoTracker signal) using ImageJ/Fiji. Mean ± SD is shown, dots represent mitochondria of individual cells. n = 7-19 independent cells per condition. Statistical analysis via Brown-Forsythe and Welch ANOVA test with p>0.05 = non-significant.

B) Representative gallery of zoomed confocal images of MIM extrusion out of the MOM during apoptosis. The MIM/matrix (magenta) and MOM (green, TOM20) of mitochondria in untreated (UT) and ABT-737 + S68354 treated (A+S) primary WT and *Gsdme^-/-^* lung fibroblasts are shown. Scale bar: 2 µm.

C) Representative overlay of the mitochondria mask (white) with DNA signal (green) of confocal images of untreated (UT) and apoptotic primary WT and GSDME^-/-^ lung fibroblasts is shown. Scale bar: 20 µm, zoom: 2µm.

D) Effect of mtDNA depletion (Rho^0^) on STING activation after 1 µM ABT-737 and S63845 (A+S) treatment without caspase inhibition. All data were normalized to time point 0 min. Mean ± SD is shown. n = 3 independent experiments. Statistical analysis by two-tailed Wilcoxon test with p>0.05 = non-significant.

E) Effect of GSDME knockdown on STING activation after 1 µM ABT-737 and S63845 (A+S) treatment without caspase inhibition. All data were normalized to time point 0 min. Mean ± SD is shown. n = 3 independent experiments. Statistical analysis by two-tailed Wilcoxon test with p>0.05 = non-significant.

F) STING activation in untreated (UT) or diABZi (5 µM) treated HeLa cells expressing the STING biosensor. All data were normalized to time point 0 min. Mean ± SD is shown. n = 3 independent experiments. Statistical analysis by Wilcoxon test with p>0.05 = non-significant. P-value is the same for siCtrl and siGSDME cells.

G) ABT737 (A) and S63845 (S) dose–response viability in wild-type mouse 1380 small cell lung cancer (SCLC) cell line for the indicated time points. Viability was measured by CellTiter-Glo (CTG), all values were normalized to the mean of the DMSO control. One-way ANOVA with Dunnett’s post-hoc test.

H-J) qPCR analysis of *Ifnα* (H), *Ifnβ* (I) and *CXCL10* (J) relative expression (rel. expr.) in WT and *Gsdme^-/-^* SCLC cells after 24 h treatment with 10 µM of ABT-737 (A) and S63845 (S) and 20 µM of QVD (Q). The data were normalized to untreated WT. ΔΔCt method. n = 3. One-way ANOVA followed by Tukey’s post-hoc test with p>0.05 = non-significant.

K-L) qPCR analysis of *Ifnα* (K) and *Ifnβ* (L) expression in mtDNA depleted (Rho^0^) WT SCLC cells after 24 h treatment with 10 µM of ABT-737 (A) and S63845 (S) and 20 µM of QVD (Q). ΔΔCt method. n = 3. One-way ANOVA followed by Tukey’s post-hoc test with p>0.05 = non-significant.

M) Schematic depicting the CRISPR-Cas9-mediated generation of mice that are deficient for GSDME or contain a point mutation preventing caspase-mediated cleavage (D270A mutation). One gRNA targeting exon 7 and one ssDNA HR template were electroporated into C57BL/6N zygotes to generate GSDME-deficient mice and GSDME non-cleavable mice in one experiment. Resulting in an offspring with a 4bp deletion in exon 7 and therefore a frameshift and a premature stop codon, indicated with a star (*), were selected as *Gsdme* knockout founders. Mice with the correct mutation at the caspase cleavage site were chosen as founders of the *Gsdme^D270A^* line. An additional silent mutation was introduced to remove the PAM sequence and aid in genotyping.

We have described *Gsdme^-/-^* previously in <https://pubmed.ncbi.nlm.nih.gov/40996439/> and include it here for completeness.

N) Representative Immunoblot showing GSDME in WT, *Gsdme^-/-^* and *Gsdme^D270A^* primary lung fibroblasts under healthy and apoptotic (2 h AS) conditions to verify the mouse models described in M.

O-P) Titration of BH3-mimetics (AS) at the indicated concentrations in combination with constant caspase inhibition (QVD, 15 µM) in HeLa STING Biosensor cells to measure PS exposure via Annexin V (O) and plasma membrane permeabilization via Draq7 (P). n = 3 independent experiments.

**Supplementary Movie 1.** Movie of representative atomistic molecular dynamics simulation of 30-mer GSDME-N rings embedded in a lipid bilayer composed of 20% CL and 80% di-oleoyl phosphatidylcholine (DOPC) with explicit solvent (with systems of >4 million atoms).

**Supplementary Table 1: Data collection, processing and refinement statistics for the cryo-EM structure of the GSDME-N pore complex.**

|  | Gasdermin E |
| --- | --- |
| EMDB-ID | 51889 |
| PDB-ID | 9H5M |
| Data Collection |  |
| Microscope | Titan Krios |
| Detector | Gatan K3 |
| Magnification | 105,000 |
| Voltage (kV) | 300 |
| Electron Dose (e^-^/Å^2^) | 65 |
| Defocus range (µm) | -1.6 - -2.4 |
| Pixel size (Å) | 0.837 |
| Movies | 32,004 |
| Initial particles | 288,669 |
| Final particles | 88,579 |
| Symmetry | C30 |
| Map resolution (Å) | 3.6 |
| FSC threshold | 0.143 |
| Refinement |  |
| Map sharpening B-factor (Å^2^) | 119.2 |
| Model composition |  |
| Non-hydrogen atoms | 57,540 |
| Protein residues | 7350 |
| Ligands | 0 |
| B factors (Å^2^) |  |
| Protein | 73.1 |
| R.m.s deviations |  |
| Bond lengths (Å) | 0.004 |
| Bond angles (°) | 0.645 |
| Ramachandran plot |  |
| Favored (%) | 86.80 |
| Allowed (%) | 10.32 |
| Disallowed (%) | 2.88 |
